# Discovery of new mycoviral genomes within publicly available fungal transcriptomic datasets

**DOI:** 10.1101/510404

**Authors:** Kerrigan B. Gilbert, Emily E. Holcomb, Robyn L. Allscheid, James C. Carrington

## Abstract

The distribution and diversity of RNA viruses in fungi is incompletely understood due to the often cryptic nature of mycoviral infections and the focused study of primarily pathogenic and/or economically important fungi. As most viruses that are known to infect fungi possess either single-stranded or double-stranded RNA genomes, transcriptomic data provides the opportunity to query for viruses in diverse fungal samples without any a *priori* knowledge of virus infection. Here we describe a systematic survey of all transcriptomic datasets from fungi belonging to the subphylum Pezizomycotina. Using a simple but effective computational pipeline that uses reads discarded during normal RNA-seq analyses, followed by identification of a viral RNA-dependent RNA polymerase (RdRP) motif in *de novo* assembled contigs, 59 viruses from 44 different fungi were identified. Among the viruses identified, 89% were determined to be new species and 68% are, to our knowledge, the first virus described from the fungal species. Comprehensive analyses of both nucleotide and inferred protein sequences characterize the phylogenetic relationships between these viruses and the known set of mycoviral sequences and support the classification of up to four new families and two new genera. Thus the results provide a deeper understanding of the scope of mycoviral diversity while also increasing the distribution of fungal hosts. Further, this study demonstrates the suitability of analyzing RNA-seq data to facilitate rapid discovery of new viruses.

## Introduction

Advances in low-cost, high-throughput sequencing technologies has revolutionized our ability to query the molecular world. Transcriptome projects are of particular use by providing targeted information about gene expression and improving the annotation of genome projects. Indeed, a number of “1K” transcriptome projects are underway, including the 1000 Plant Genome project, Fish-T1K project, 1K Insect Transcriptome Evolution (1KITE) project, and 1000 Fungal Genome and Transcriptome project. These large scale, collaborative projects have contributed to the vast amount of sequencing data available. As of February 2018, more than 16,000 petabases have been deposited in the Short Read Archive (SRA) at NCBI, with over 6,000 petabases available as open-access data.

Beyond revealing gene expression patterns within the target organism, these datasets are a source for additional insights as novel RNA sequences, specifically viral genomic sequences, have been identified within them. In an early example, analysis of RNA-seq data from a soybean cyst nematode revealed the presence of four different RNA viruses [1]. A more broad scale analysis of 66 non-angiosperm RNA-seq datasets, produced as part of the 1000 Plant Genome project, identified viral sequences in 28 plant species [2]. Additionally, the first viral sequences for any member of the family of spiders *Nephilidae* were identified through a secondary analysis of the RNA-seq data produced during the spider genome sequencing project [3].

While over 200 mycoviruses have been described, the full depth of the mycoviral landscape remains to be determined as certains biases have impacted the discovery of mycoviral sequences. Commonly infections of fungi appear cryptic, having little to no apparent impact on fungal health, thus occluding the presence of the virus. Additionally, many of the fungi studied are important human or plant pathogens, but this represents only a fraction of the known diversity of fungi. Advancements in mycovirus discoveries have been aided by screening the transcriptomes of various collections of fungi; study systems include those fungi associated with the seagrass *Posidonia oceanica* [4], five common plant-pathogenic fungal species [5], and soybean phyllosphere phytobiomes [6].

In the biosphere, fungi are not found in isolation; instead, they are members of complex environments that include members from some or all other kingdoms of life. Characterizing the responses and contributions that fungi make to these environments also includes a full understanding of the dynamics at play within the fungi. We hypothesized that mycoviral infections are more pervasive than currently known, and as such, undertook a systematic approach to identify new mycoviral species by querying the publicly available fungal transcriptomic datasets at the Short Read Archive hosted at NCBI. Our bioinformatic approach provides a robust method for the identification of new viral species from raw sequencing data as we identified 59 viral genomes, 53 of which are described here for the first time.

## Results and Discussion

Viral RNA-dependent RNA polymerases (RdRP) are essential enzymes in the genomes of RNA viruses that have no DNA stage. The domain organization of this protein is described as a right hand, where three subdomains are the palm, fingers, and thumb [7]. While some of the motifs found within these subdomains are conserved among all polymerase enzymes, specific structures and motifs are found only in RNA-dependent RNA polymerases. In addition to unique signatures within the sequences themselves, virus-like RdRPs are not encoded in the DNA genomes of eukaryotic organisms, thus viral RdRPs can be specifically targeted as a marker for sequence-based analyses.

We previously determined that contigs assembled from fungal RNA-seq data can be queried for viral RdRP-specific domains [8]. As such, we undertook a systematic analysis of all RNA-seq datasets from Pezizomycotina fungi (division: Ascomycota; subkingdom: Dikarya) available at the Short Read Archive (SRA) at the National Center for Biotechnology Information (NCBI) on July 21, 2017. An initial list of 1,127 unique BioProjects was identified, which was manually curated to identify 569 BioProjects suitable for this analysis (S1 Table). Samples were excluded for a variety of reasons including: individually listed BioProjects were determined to belong to larger groups of experiments using the same fungal strain and were therefore combined and a single representative sample was selected; samples containing a mix of fungi and RNA from other kingdoms were excluded as the host of any putative viral sequences could not be determined; samples where the read sequences were too short for assembly or unavailable for download.

### Virus-like RdRP Discovery Pipeline

A computational pipeline was created to automate the steps following selection of a raw reads SRA file available and identification of putative viral RdRP sequences. The majority of the steps used publicly available programs in conjunction with a small number of custom Perl scripts. The overall structure of the pipeline was optimized to efficiently work within a HTCondor-managed system including the use of the DAGMan (Directed Acyclic Graph Manager) meta-scheduler to coordinate the required dependencies between pipeline steps.

Due to the highly specific nature of viral RdRP protein sequences, the key feature of this pipeline was the use of a customized Pfam database which included the viral RdRP families: RdRP_1, RdRP_2, RdRP_3, RdRP_4, RdRP_5, and Mitovir_RNA_pol. To err on the side of over-inclusiveness, HMMER *hmmscan* was run with an e-value cut-off of 10.0, allowing for the identification of any sequence with limited similarity to the viral protein domains. Putative RdRP sequence(s) were identified in a total of 1067 contigs from 284 of the 569 samples. The actual number of viruses was expected to be less than this total number; reasons include a false positive match to the viral RdRP signatures, contigs that are the forward and reverse strand sequence of a single virus, or contigs that are partial matches along a single virus sequence. To confirm similarity to known viral sequences, contigs were evaluated via a BLASTX search versus the ‘nr’ database at NCBI followed by manual evaluation and curation of contigs with hits to known mycoviral sequences.

### Summary of identified viral sequences

In total, 59 complete, RNA mycoviral genomes were identified in 47 of the 569 SRA samples analyzed. A global analysis of the sequencing data available and the resultant virus sequences along with the set of known mycoviruses available at Virus-HostDB (version March 23, 2018, http://www.genome.jp/virushostdb) demonstrates a number of patterns and trends (Fig 1). There is an uneven distribution of data being generated within the Pezizomycotina subphylum whereby certain Classes are more abundantly represented than others (illustrated by the width of the band going from “Fungal Class” to “Fungal Genus”). For instance, the Sordariomycetes and Eurotiomycetes host 77% of the sequencing projects while the remaining eight Classes host the other 23% of datasets. Unsurprisingly, fungi known to be key pathogens of plants and humans are present within the Sordariomycetes and Eurotiomycetes, including the well-studied genera *Fusaríum, Aspergillus*, and *Penicillium.* Indeed, seven genera alone account for 51% of all sequencing projects analyzed in this study, with the remaining 49% of datasets spread among 176 other genera.

**Fig 1.**
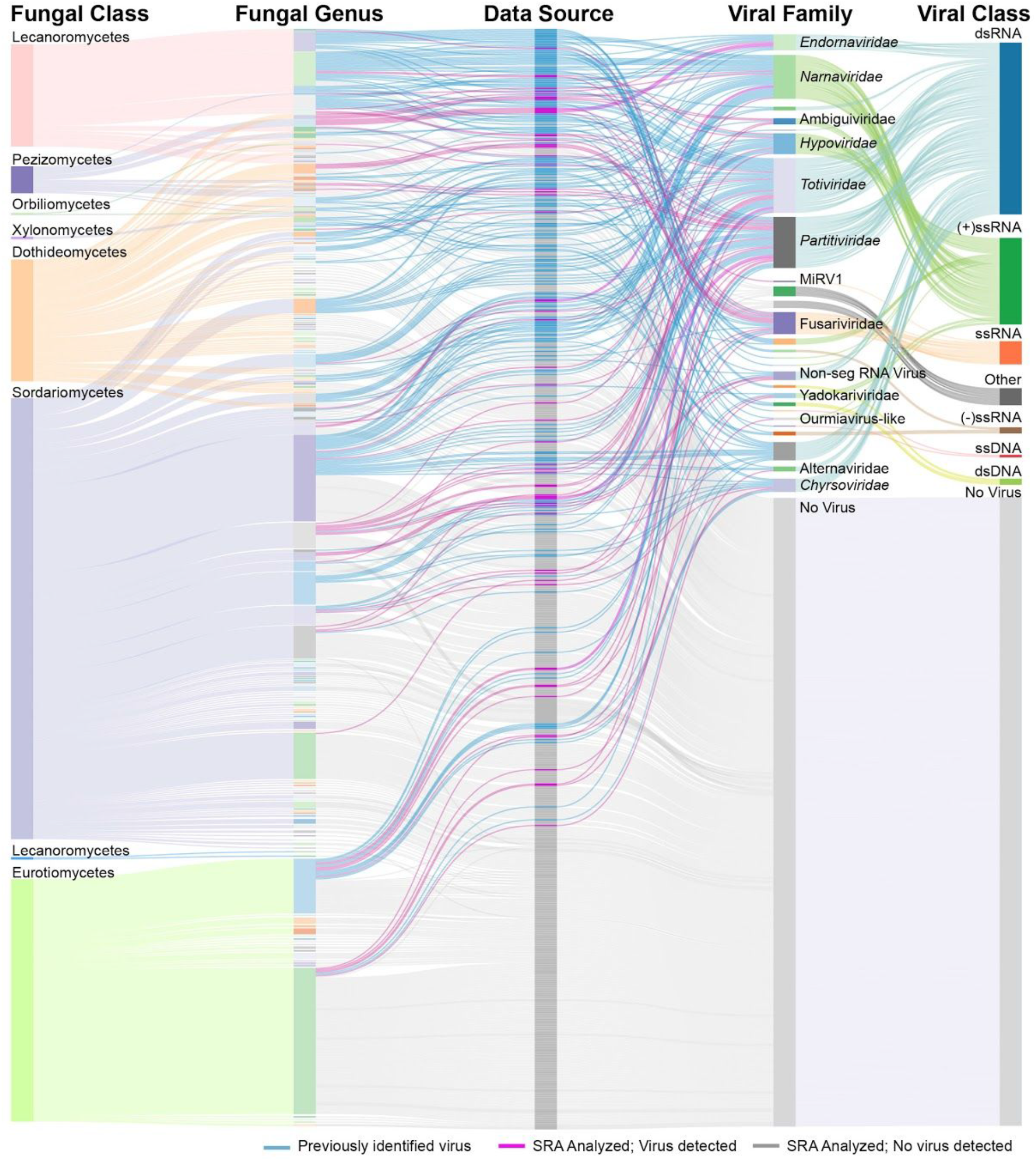
Sankey diagram summarizing both the SRA samples queried in this study and recognized mycoviral sequences. The height of the bars in each column is indicative of the number of samples. Classification of fungal host is summarized in the two, leftmost columns. Column “Fungal Class”: fungal species are sorted into one of eight taxonomic classes, all genera from the same class are the same color; Column “Fungal Genus”: further separates the fungi into their respective genera; colors used in this column are to highlight the individual genera but similar colors have no relation to one another. The third column, “Data Source” is color-coded by the origin of the sample where blue indicates a known mycovirus, while pink and grey indicate an SRA sample was analyzed and either a virus sequence was or was not detected. The two rightmost columns summarize the information related to virus taxonomy. Column 4: “Viral Family” organizes the samples with either a known or discovered virus by viral family. The names of families and groups that contain a virus identified in this study are indicated. Column 5: “Viral Class” summarizes the viral genomes by nucleic acid and/or strandedness. Samples without a virus detected are in the category “No Virus”.

Virus discovery as a function of number of sequencing projects favors the less data-rich Classes of fungi. While Sordariomycetes and Eurotiomycetes host 61% of the viruses identified in this study, the discovery rate is 17%. Conversely, the less-studied classes Leotiomycetes, and Pezizomycetes had virus discovery rates of 21%, and 26% respectively. Clearly, however, a certain threshold of data is required for virus discovery, as the Classes Xylonomycetes, Orbiliomycetes, and Lecanoromycetes had a combined total of seven sequencing projects but only one virus was identified.

Comparing the distribution of the known set of Pezizomycotina-hosted mycoviruses with those identified in this study demonstrates similar distribution patterns. Members of the Sordariomycetes and Eurotiomycetes Classes are host to 58% of known mycoviruses, with Leotiomycetes being also abundant. Gains in under-represented Classes are observed, as only two mycoviruses were previously known to be hosted by a Pezizomycetes fungus, and five viruses were identified in this study. Additionally, the first virus from the Class Orbiliomycetes was found in this study.

Classification of the virus sequences identified in this study reveals that 19% of viruses are from unclassified viral families. This is similar to the known set of mycoviruses, where 18% of viruses are also unclassified. Further, dsRNA viruses are the most abundant category within the two viral datasets, with 63% and 54% belonging to those identified in this study and the known set respectively. The ratio of dsRNA to ssRNA viruses in this study is higher than the known dataset, indicating an under-identification of ssRNA viruses (18% versus 27%, this study and known dataset respectively). This result may indicate that the RNA preparation methods for RNA-seq experiments are biased against ssRNA viruses.

An in-depth analysis and characterization of each of the 59 viral sequences is described below (Table 1). Viral genomes have been divided into two major categories: dsRNA viruses and (+)ssRNA viruses then further separated by viral family and, where applicable, genus. In total, viruses were identified from eight of the currently recognized virus families and as such are described within the context of the expected characteristics. The remaining viruses form groups with other recently identified mycoviruses leading to the characterization of new mycoviral families and genera within an existing family.

**Table 1:**
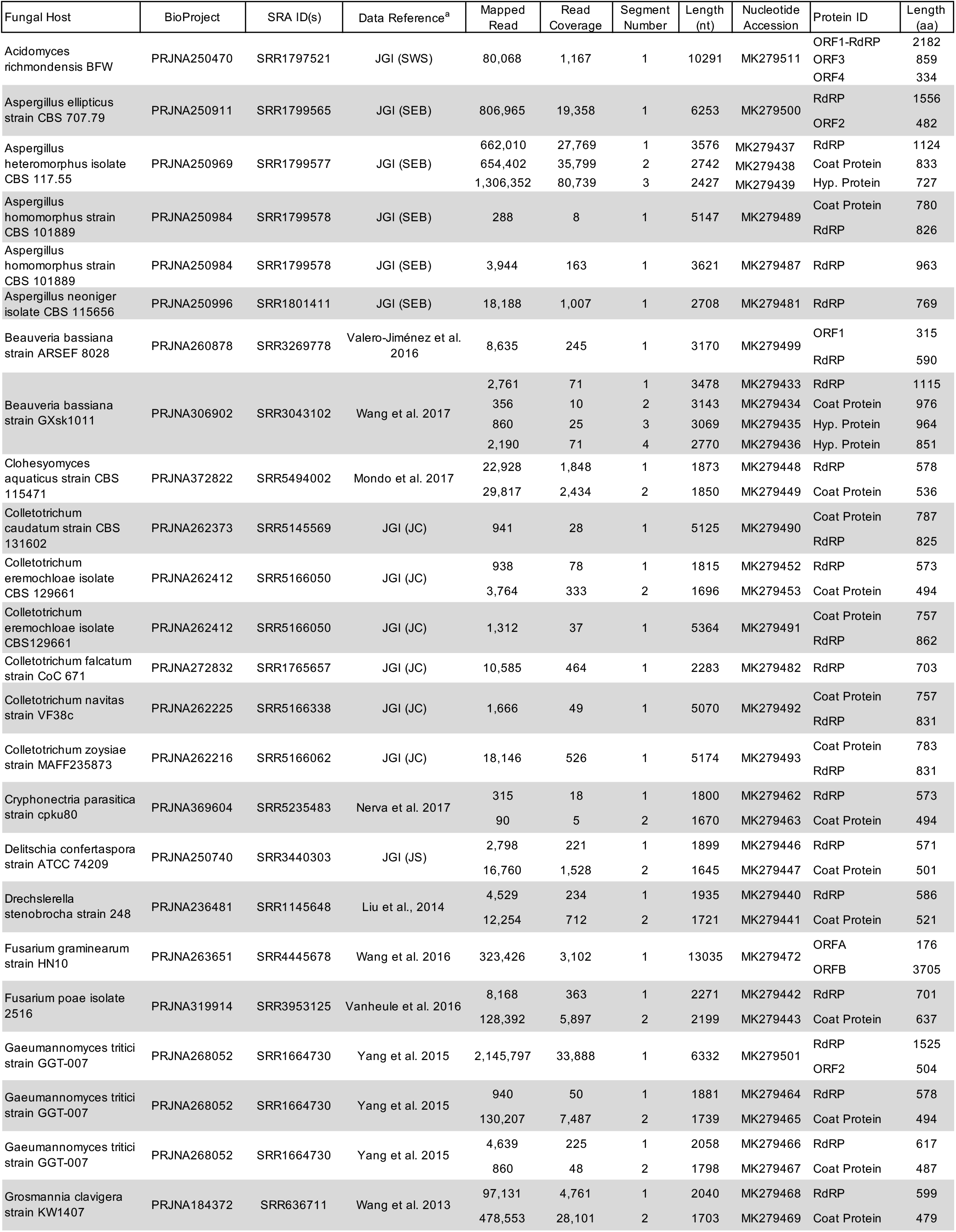
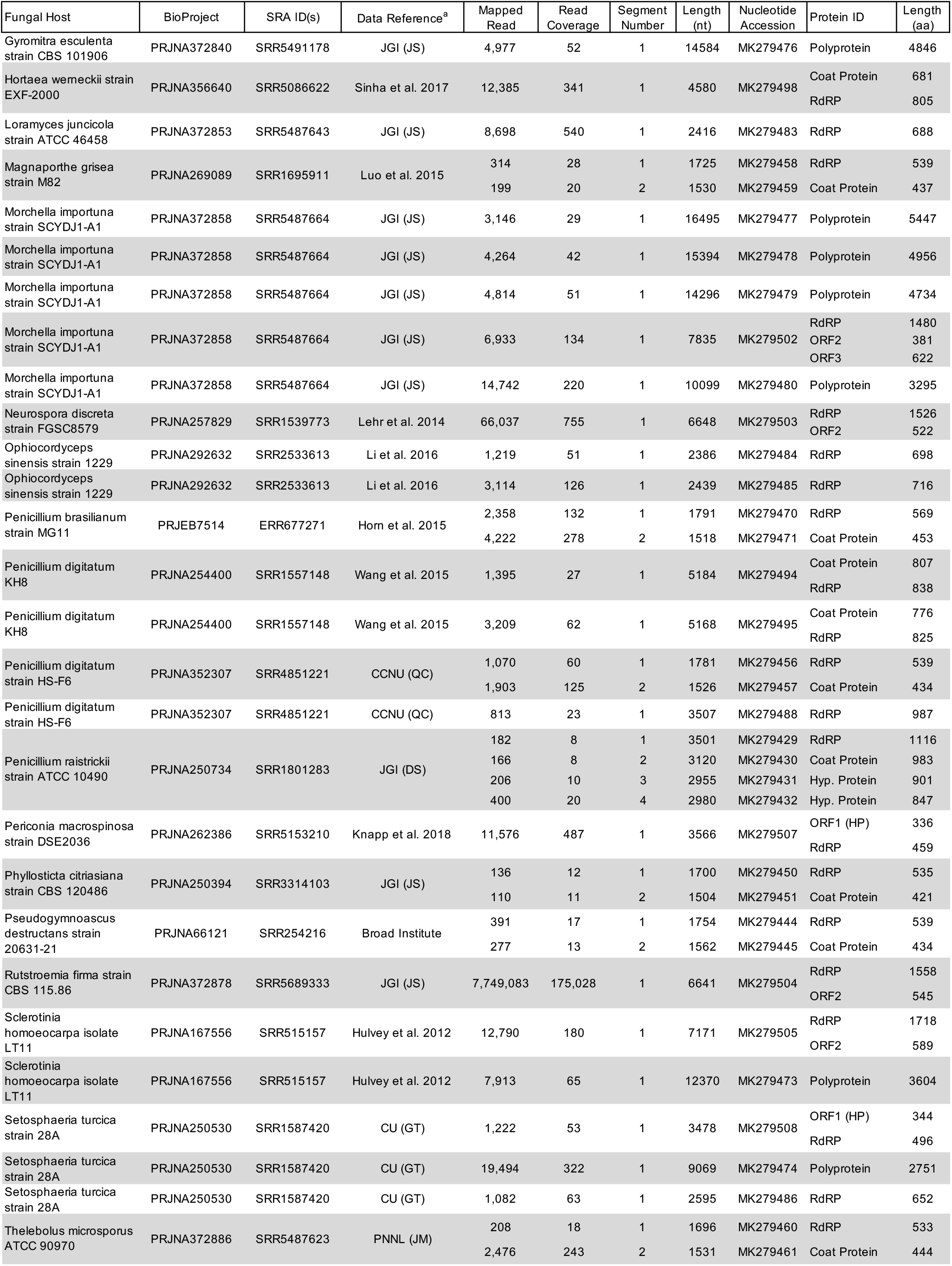
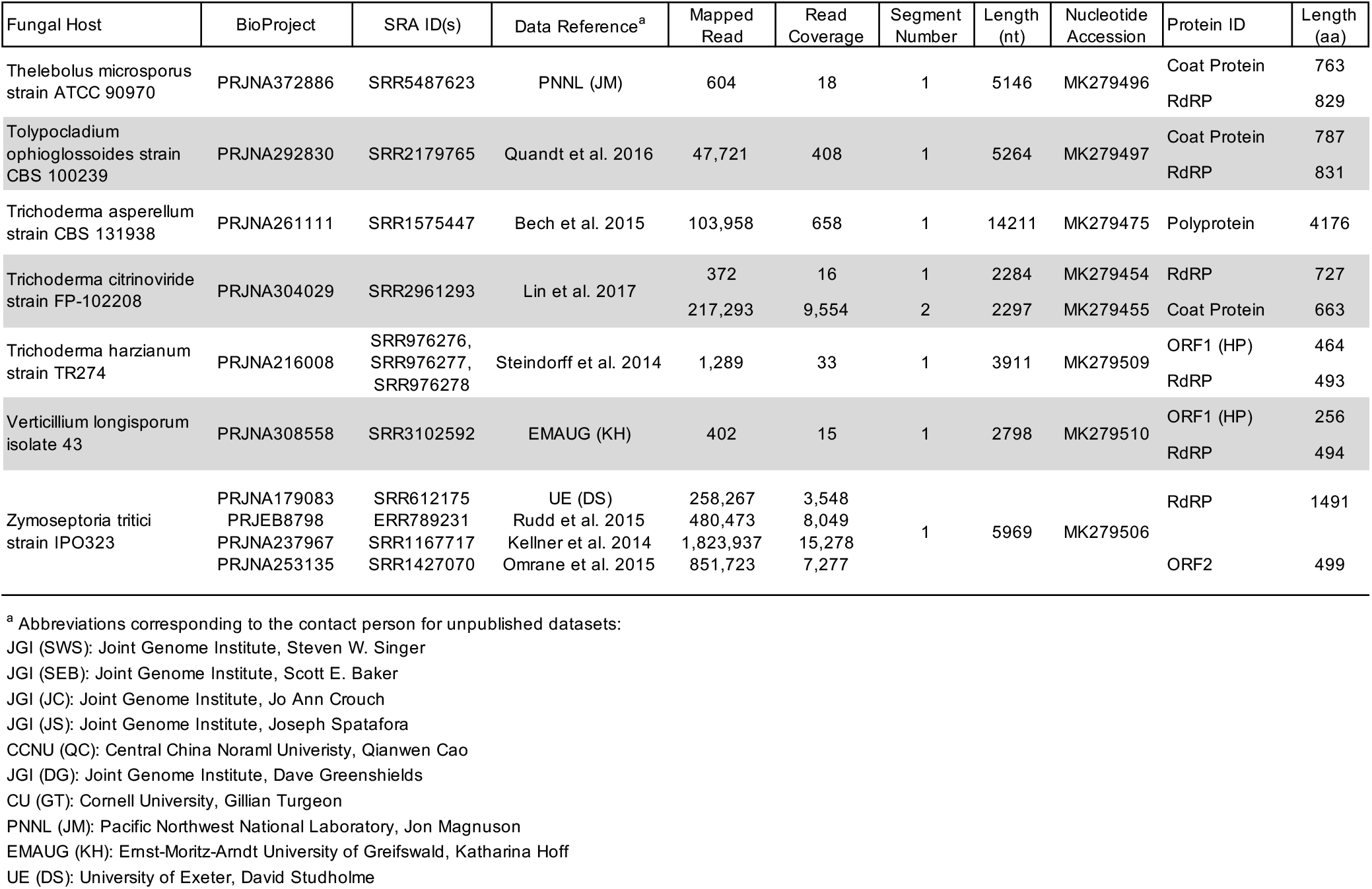
Summary of mycoviral genomes identified from fungal RNA-seq datasets

### Double stranded RNA (dsRNA) Viruses

A total of 34 viruses were categorized as dsRNA viruses due the identity of the top BLASTP match. To further characterize the relationship of these sequences, a phylogenetic tree based on the RdRP amino acid sequences from the 34 identified viruses and the top BLASTP matches was constructed (Fig 2). Globally, five groups are visible, where three are known viral families (*Totiviridae, Partitiviridae, and Chrysoviridae)* and two represent potentially new families. Properties characteristic of each group of viruses are described in detail below.

**Fig 2.**
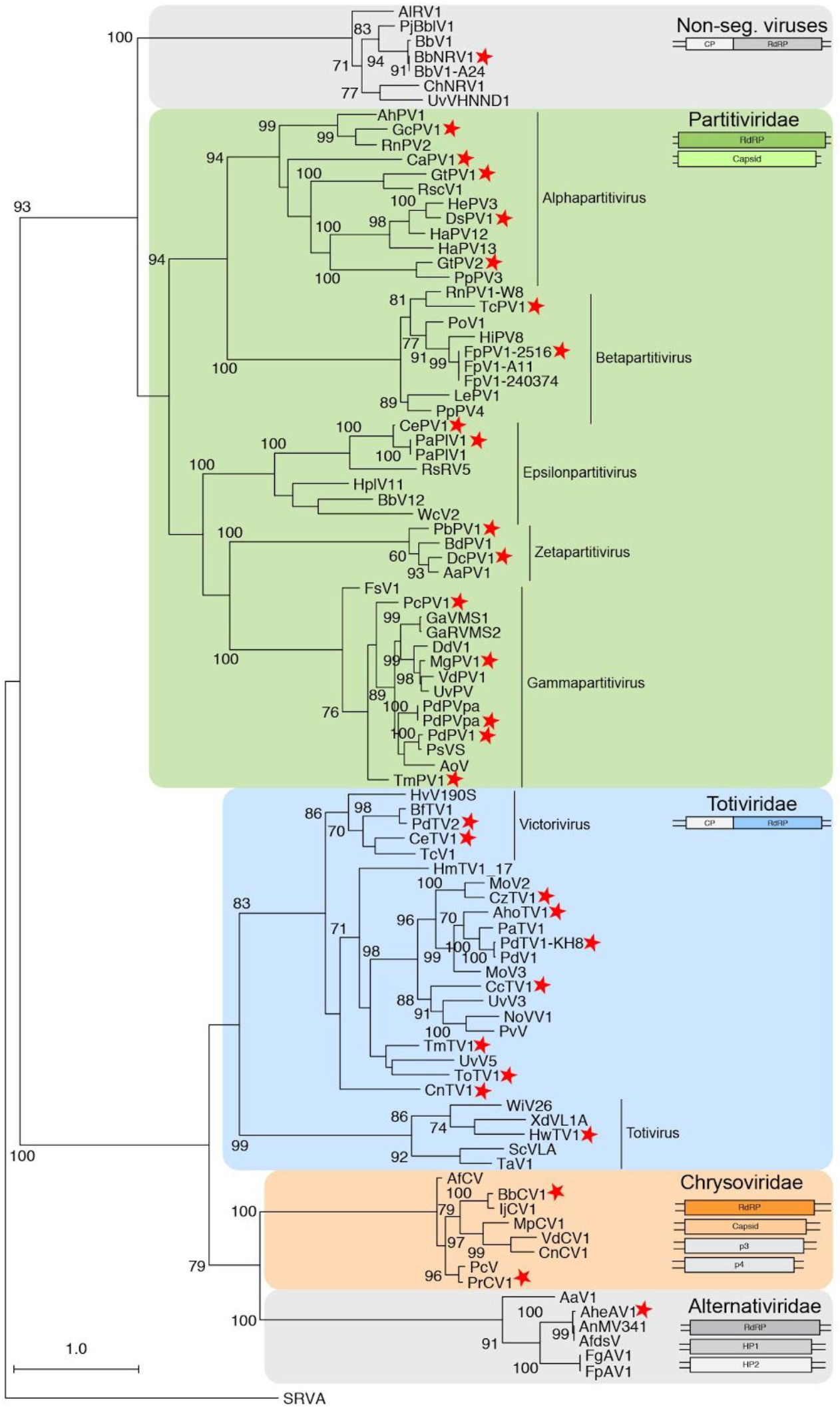
Phylogram of the RdRP- containing ORF for viruses classified as dsRNA viruses. The sequence from *Simian Rotavirus A* (SRVA) serves as the outgroup. Three þ officially recognized families are present: *Partitiviridae, Chrysoviridae*, and *Totiviridae* and are highlighted with a green, orange, or blue box respectively. Gray boxes surround families currently unrecognized. The structure of the genomic segment(s) for each highlighted group is depicted below the family name. Genera within the *Partitiviridae* and *Totiviridae* family are indicated. Viruses identified in this study are noted with a red star. Numbers at the nodes indicate bootstrap support over 70% (1000 replicates). RdRP: RNA-dependent RNA Polymerase; CP: Coat Protein; HP: Hypothetical Protein.

### Non-segmented RNA Virus (BbNRV1)

Recently, dsRNA viruses have been identified from various fungi that encode two ORFs within the single-segment genome, where the RdRP sequences have more similarity to the RdRPs of partitiviruses (multi-segment genomes) than totiviruses (mono-segment genomes). The virus *Beauveria bassiana non-segmented dsRNA virus 1* (BbNRV1), assembled using the RNA-seq data from a transcriptome experiment of the highly virulent isolate *B. bassiana* ARSEF 8028 [9], groups with these previously identified viruses (Fig 2). A second virus from this group was also identified in this analysis: *Colletotrichum higginsianum non-segmented RNA virus 1* (ChNRV1; this virus served as an internal control for the virus discovery pipeline as it was previously identified via a similar approach from two different *Colletotrichum higginsianum* RNA-seq datasets [8].

BbNRV1 and ChNRV1, together with four previously identified viruses, form a strongly supported group that is separate from other characterized fungal viruses (Fig 2). It has been noted by multiple researchers that these fungal viruses share similarities with a family of plant viruses, the *Amalgaviridae*, specifically the mono-segmented genome structure and partitivirus-like RdRP sequence. Therefore, a phylogenetic analysis of the RdRP protein sequence was performed which demonstrates that, while the mono-segmented genome structure is shared, this new clade of fungal viruses is also distinct from the plant amalgaviruses (Fig 3A). Indeed, the amalgavirus group of plant viruses branches off before the BbNRV1-related and *Partitivirus* clades. A percent identity matrix generated with RdRP and CP coding sequences further demonstrates a separation between this new group and existing virus families (Fig 3B). For the RdRP pairwise comparisons, 50% identity easily differentiates the three clades from each other. Additionally, the CP sequence comparisons demonstrate shared similarities within the three clades and less than 20% similarity is shared between the clades.

**Fig 3.**
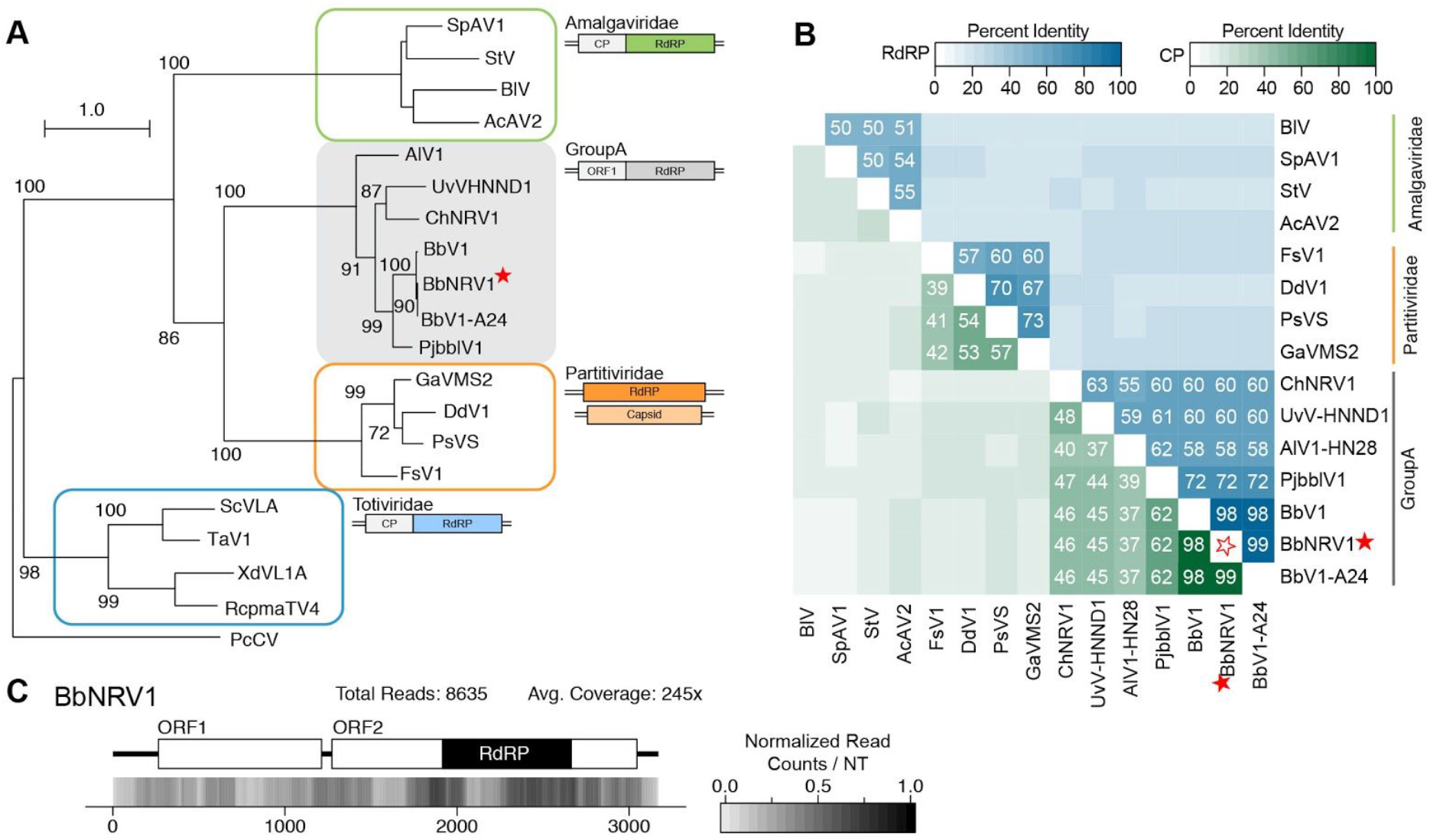
Analyses of identified viral genomes in relation to known *Amalgaviridae* and *Partitiviridae.* (A) Phylogram of the ORF containing the RdRP motif for members of the new mycovirus family along with select partitiviruses, totiviruses and amalgaviruses; sequence from *Penicillium chrysogenum virus* (PcCV) serves as the outgroup. Full names and accession numbers for protein sequences used in the alignment are in Table S2. Scale bar on the phylogenetic tree represents 1.0 amino acid substitutions per site and numbers at the nodes indicate bootstrap support over 70% (1000 replicates). Diagrams illustrating the genome structure and predicted ORFs are present for each group of viruses. (B) Percent identity matrix, generated by Clustal-Omega 2.1, of new mycoviral family, partitiviruses, and amalgaviruses. The top half of the matrix, in the blue scale, is the percent identity of the RDRP protein coding sequence and the bottom half of the matrix, in the green scale, is the percent identity of the putative CP. For sake of clarity, the 100% identity along the diagonal has been removed and outlined stars are used to note junction point for the row and column of the virus identified. Where the identified viral sequences have significant similarity to known viral sequences is indicated, specifically RdRP sequences with similarity ≥50% and CP sequences with similarity >30%. (C) Density plot of read counts per nucleotide across the viral genome for BbNRV1. Graphical depiction of the genome structure, ORFs and predicted motifs are on along the top and a heatmap of normalized read counts is on the bottom. Read counts are normalized to a 0 −1 scale, where the maximum read count for a given genome equals 1. Total reads mapped to the virus genome and average read coverage are noted. The red star indicates the virus identified in this study. RdRP: RNA-dependent RNA Polymerase; CP: Coat Protein; ORF: Open Reading Frame.

The genome structure of viruses within this group consists of two predicted ORFs surrounded by variable length UTRs (Fig 3C). The 5’UTR lengths fall into two distinct ranges: less than 40 nt for *Penicillium janczewskii Beauveria bassiana-like virus 1* (PjBblV1) [4], ChNRV1 [8], and UvHNND1 [10], or between 260 to 320 nt for the three viruses isolated from *B. bassiana* [11, 12] and *Alternaria longipes dsRNA virus 1* (AIV1-HN28) [13]. The protein encoded by ORF1 is predicted to be from 314 to 394 aa and in a +1 frame relative to the protein encoded by ORF2. The intergenic sequence lengths are generally less than 100 nt, with the exception of AIV1-HN28. ORF2 nucleotide sequences range from 1578 to 1773 nt, and encoded proteins, which encode an RdRP domain, range in size from 525 to 590 aa in length. A slippery site sequence identified between ORF1 and ORF2 of ChNRV1 suggested that an ORF1-ORF2 fusion protein may be made via a −1 ribosomal frameshift; indeed, trypsin digest followed by mass spectrometry of a protein near the predicted weight of the fusion protein returned signatures for both the ORF1 and ORF2 protein sequences [8]. Numerous attempts to isolate virus particles definitely produced by a virus within this group remain unsuccessful [4,8,11]. Due to the shared genome structure and partitivirus-like RdRP protein sequence, early viruses from this group were tentatively grouped with the plant *Amalgaviridae* family. However, with the identification of additional genome sequences, a more in depth phylogenetic analysis demonstrates the mycoviruses described here are separate. A new genus, “Unirnavirus”, has recently been suggested to reflect the single, unified genome segment that contains both ORFs [12]. As percent identity is a common criteria for species demarcation, the pairwise comparisons here provide possible thresholds for determining new species within this group of viruses; excluding the three viruses isolated from different strains of *B. bassiana*, a threshold of 75% identity in the RdRP protein sequence and 70% in the ORF1 coding sequence would be appropriate criteria to distinguish between the viruses isolated from unique fungal hosts.

### Alternaviridae

Initially, a single segment encoding a single gene with an RdRP domain was identified from an *Aspergillus heteromorphus* sequencing sample (PRJNA250969; unpublished). BLAST alignment to ‘nr’ identified five related RdRP mycoviral sequences from viruses with multi-segmented genomes. *Fusarium graminearum alternavirus 1* (FgAV1) includes at least one additional genomic segment while *Alternaria alternata virus 1* (AaV1) [14] and *Aspergillus foetidus dsRNA mycovirus* (AfdsV) [15] each contain a total of four genomic segments. Using the additional sequences from these related viruses, two more contigs were identified from the *Aspergillus heteromorphus* Trinity assembly. Thus, the AheAV1 genome contains at least three genomic segments, the largest of which encodes the RdRP enzyme and the two smaller segments that encode proteins of unknown function (Table 1). Recently it has been proposed that a new family be established, “*Alternaviridae*” [15] we support the formation of this family, and the requisite genus (*Alternavirus*) and propose that AheAV1 is also a member of this new mycoviral family.

Previously it was observed that members of this clade group together, but separate from other families [14, 15]. A phylogenetic analysis using the RdRP coding sequence demonstrates that the group containing AheAV1 forms a well supported clade that appears related to the *Chrysoviridae* and *Totiviridae* families (Fig 2). Members of the Totiviridae family have single-segment genomes containing two ORFs, while members of the Chrysoviridae have multi-segment genomes. While the RdRP segment sizes are similar between alternaviruses and chrysoviruses (~3,500 bp), the remaining two to three segments are hundreds of base pairs shorter than those of chrysoviruses. Further, the proteins of unknown function have no predicted domains or shared sequence homology outside of this group of viruses. The three viral RdRP sequences isolated from different species of *Aspergillus*, including AheAV1, are most closely related; similarly, the RdRP sequences from the two viruses from *Fusarium* species are most related (Fig 2). Indeed, a pairwise percent identity comparison of both the RdRP and HP1 protein sequences demonstrates the very high identity among the viruses from the two *Fusarium* and also from the two *Aspergillus* viruses (Fig 4A). This analysis also illustrates a clear delineation between alternaviruses and chrysoviruses.

**Fig 4.**
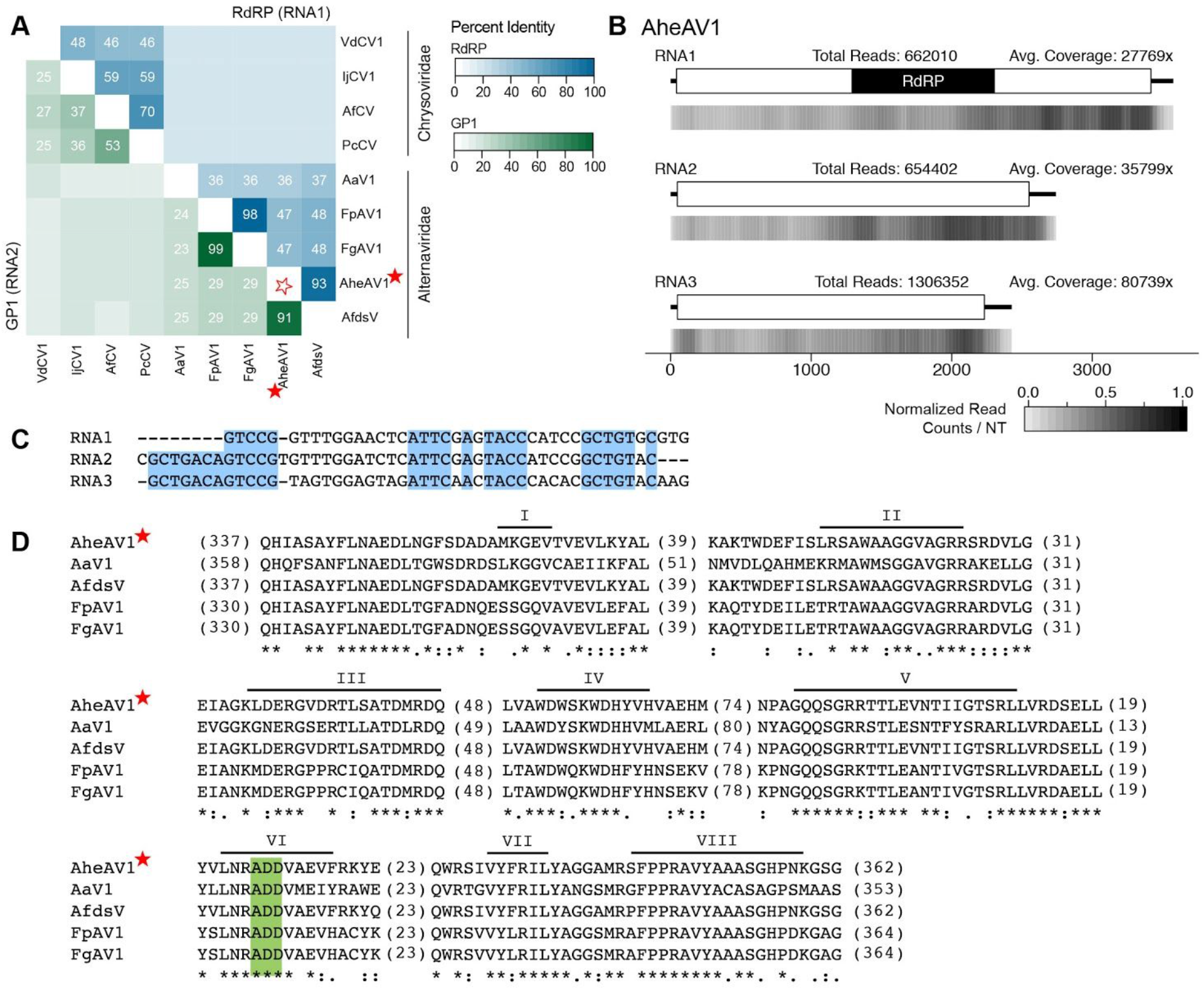
Analyses of *Aspergillus heteromorphus alternavirus 1* (AheAV1), a member of the recently proposed family *Alternaviridae.* (A) Percent identity matrix, generated by Clustal-Omega 2.1, of alternaviruses and members of the *Chrysoviridae* family. The top half of the matrix, in the blue scale, is the percent identity of the RdRP protein coding sequence (from the RNA1 segment) and the bottom half of the matrix, in the green scale, is the percent identity of the second largest genomic fragment, encoding GP1. For sake of clarity, the 100% identity along the diagonal has been removed and outlined stars are used to note junction point for the row and column of the virus identified. Values ž20% identity are noted. (B) Density plot of read counts per nucleotide across the viral genomic segments of AheAV1. Graphical depiction of the genome structure, predicted ORFs and motifs for each segment are on top and a heatmap of normalized read counts is on the bottom. Read counts are normalized to a 0 −1 scale, where the maximum read count for a given genome equals 1. Total reads mapped to the virus genome and average read coverage are noted. (C) Alignment of 5’UTR sequences of the three genomic segments of AheAV1. Conserved nucleotides are highlighted in blue. (D) Alignment of RdRP domains of Alternaviruses found in (A). The eight conserved RdRP motifs of dsRNA polymerases are noted along the top of the alignment and the ADD triad, found only in Alternavirus sequences, is highlighted in green. The number of amino acids not shown in alignment are noted in parentheses. Viruses identified in this study are indicated with a red star. RdRP: RNA-dependent RNA Polymerase; GP1: Gene Product.

A common feature to viral genomes with multiple segments is the presence of a conserved sequence within the 5’ UTRs of each fragment. Sequence conservation was described within the UTRs of the other putative alternaviruses, including AfV [15] and AaV1 [14]. Alignment of the 5’UTR sequences of the three genomic segments of AheAV1 demonstrates sequence conservation (Fig 4C). Alignment of the source RNA-seq reads to the viral genome segments reveals a high amount of total reads, with over four million reads mapped across the three segments (Fig 4B). This represents nearly 25% of the reads used for the analysis (reads that did not align to the fungal genome) and over 5% of the total reads from the sample. In an effort to identify a possible fourth genomic fragment for AheAV1, both the 5’UTR conserved sequence and the high read count were used as criteria to further query the Trinity assembled contigs however no additional contigs were identified.

An alignment of the RdRP protein sequences from the five alternavirus genomes reveals the presence of the expected eight conserved domains found within viral RdRP proteins (Fig 4D). Certain differences are shared among these viruses that are unique when compared to known viral sequences. Specifically, the triad within domain VI has an alanine (ADD) instead of the nearly universally conserved glycine (GDD).

Taken together, a number of criteria identify a new *Alternavirus*: multiple genome segments, where the largest encodes an RdRP protein, and the presence of an alanine within motif VI of the RdRP protein sequence. Classification of a new species within the genus *Alternavirus* remains to be established. To date, the distinguishing factors between viruses within this genus are the fungal host species and the number of genomic segments. Using these criteria, AheAV1 is a new species within the genus *Alternavirus.*

### Totiviridae

Ten of the viral sequences identified in this study are similar to other known members of the *Totiviridae* family, nine within the genus *Victorivirus* and one within the genus *Totivirus.*

#### Totiviruses

*Hortaea werneckii totivirus 1* (HwTV1), the first viral sequence described for this fungal species to our knowledge (PRJNA356640; [16]), is a member of the *Totivirus* genus (Fig 2). Species demarcation within *Totivirus* requires biological characterization of the virus within a host cell; however, isolation of a virus from a distinct host species and less than 50% sequence identity of the RdRP at the protein level is a proxy for identifying a separate species [17]. HwTV1 was identified from a new fungal host, *H. werneckii*, and shares less than 44% sequence similarity to the most closely related known totiviruses (S1A Fig) thus satisfying the proxy criteria for a new virus species. The RdRP protein sequence of HwTV1 shares the greatest similarity to two totiviruses identified within the fungus *Xanthophyllomyces dendrorhous*, XdV-L1-A and XdV-L1-B [18], along with the invertebrate virus *Wuhan insect virus 26* (WiV26), isolated from RNA-sequencing data of a mixed sample of flea and ants isolated in Hubei, China [19].

A common feature to members of the *Totivirus* genus is the presence of two partially overlapping open reading frames that produce a fusion protein via a −1 frameshift event with the 5’ ORF encoding the major capsid protein and the 3’ ORF encoding an RdRP. Characteristics of a frameshift site are (1) a heptanucleotide sequence ‘slippery site’ with the sequence X XXY YYZ and (2) a downstream RNA pseudoknot [20]. As expected, the two open reading frames in HwTV1 are frame 3 (CP) and frame 2 (RdRP). A canonical slippery site, G GGU UUU, is present upstream of the ORF1 stop codon at position 1,940 − 1,946 nt, and two predicted pseudoknots are present immediately 3’ of the slippery site (S1B Fig). A conserved domain search within the encoded proteins reveals the presence of the major coat protein (L-A virus) domain in ORF1, in addition to the RdRP domain in ORF2 (S1C Fig) A total of 12,385 reads align to the HwTV1 genome, resulting in an average coverage of 341x, with slightly higher coverage observed across the 5’ half of the genome (S1C Fig).

**S1 Fig.**
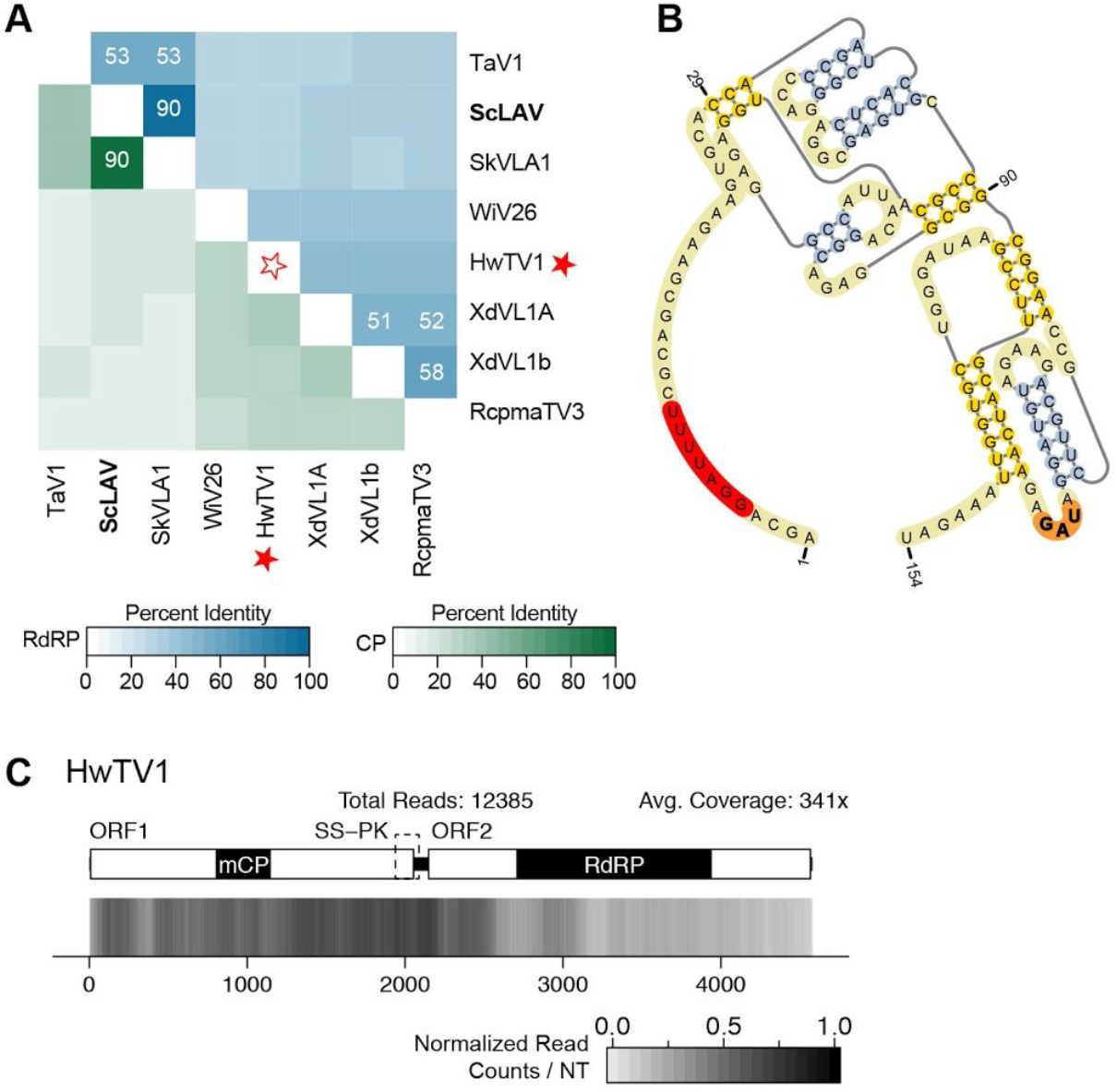
Analyses of *Hortaea werneckii totivirus 1* (HwTV1), a virus from the *Totivirus* genus within the *Totiviridae* family. (A) Percent identity matrix, generated by Clustal-Omega 2.1, to select totiviruses. ICTV-recognized members of the genus are in bold. The top half of the matrix, in the blue scale, is the percent identity of RdRP protein sequences and the bottom half of the matrix, in the green scale, is the percent identity of CP protein sequences. For sake of clarity, the 100% identity along the diagonal has been removed and outlined stars are used to note junction point for the row and column of the virus identified. Identities >50% are noted. (B) Slippery site and two pseudoknot structures predicted using DotKnot and visualized with Pseudo Viewer3. The putative slippery site heptamer (GGAUUUU) is highlighted in red and the CP stop codon (UAG) is in highlighted in orange. (C) Density plot of read counts per nucleotide across the viral genome of HwTV1. Graphical depiction of the genome structure, encoded genes and predicted protein motifs are on top and a heatmap of normalized read counts is on the bottom. The predicted slippery site and pseudoknot region is indicated by the dashed box. Read counts are normalized to a 0 −1 scale, where the maximum read count for a given genome equals 1. Total reads mapped to the virus genome and average read coverage are noted. Viruses identified in this study are noted with a red star. RdRP: RNA-dependent RNA Polymerase; CP: Coat Protein; mCP: major coat protein domain of L-A virus.

#### Victoriviruses

The remaining nine viruses in the *Totiviridae* family were found within two branches of the *Vicŧorivirus* genus. The prototype virus of this genus, *Helminthosporium victoriae virus 190S* (HvV190S) [21], is present within one clade along with *Coiletotrichum eremochioae totivirus 1* (CeTV1) (PRJNA262412; unpublished), *Colletotrichum navitas totivirus 1* (CnTV1) (PRJNA262225; unpublished), and *Penicillium digitatum totivirus 2* (PdTV2) (PRJNA254400; [22]). A second clade contains *Colletotrichum caudatum totivirus 1* (CcTV1) (PRJNA262373; unpublished), *Colletotrichum zoysiae totivirus 1* (CzTV1) (PRJNA262216; unpublished), *Penicillium digitatum totivirus 1-KH8* (PdTV1-KH8) (PRJNA254400; [22]), *Thelebolus microsporus totivirus 1* (TmTV1) (PRJNA372886; unpublished), *Tolypocladium ophioglossoides totivirus 1* (ToTV1) (PRJNA292830; [23]), and *Aspergillus homomorphus totivirus 1* (AhoTV1) (PRJNA250984; unpublished) along with other victoriviruses (Fig 2).

Alignment of the RNA-seq reads to the viral genomes reveals a wide range of average coverage values: from 8x to 526x (S2C and S2D Fig). A peak in coverage observed in CcTV1 and CzTV1 immediately precedes an octanucleotide sequence (AGGGUUCC) which has been observed in the 5’UTR of some other victoriviruses including *Alternaria arborescens vicŧorivirus 1* (AaVV1) [24] and *Epichoë festucae virus 1* (EfV1) [25]. The peak in coverage of CnTV1 immediately precedes an octanucleotide sequence slightly different than previously reported (CGGCCUCC). A similar peak in coverage was not observed for AhoTV1, which also contains the AAGGGUUCC octamer sequence within the 5’UTR, although this viral sequence had the lowest average coverage. The genome structure diagrams drawn above the coverage heat maps highlight features of the viruses that are further described below.

Unlike the frame-shifting observed in totiviruses, the RdRP of Victoriviruses is expressed as a separate protein from the coat protein [26] via a termination-reinitiation strategy [27]. The AUG start codon either overlaps the stop codon of the upstream of ORF1 or slightly precedes it, and is nearly always found in the −1 frame relative to ORF1 [21]. The tetranucleotide sequence, AUGA, is observed for PdTV2, PdTV3, CzTV1, and AspTV1. For CnTV1, TmTV1, and ToTV1, the last nucleotide of the ORF1 stop codon overlaps the ORF2 start codon, creating the pentamer sequences <u>UGA</u>UG, <u>UAA</u>UG, and <u>UAA</u>UG respectively. In each of these seven instances, there is the expected −1 frameshift between ORF1 and ORF2. For CeTV1 and CcTV1, a two nucleotide spacer is observed between the start codon of ORF2 and the stop codon of ORF1 (AUGCA<u>UGA</u> and AUGAA<u>UAA</u> respectively) that is similar to the sequences observed in SsRV1 and SsRV2 (AUGAA<u>UAA</u> and AUGAG<u>UAA</u> respectively) [28]. Thus ORF2 of CeTV1 and CcTV1, as well as SsRV1 and SsRV2, is found in the +1 frame relative to ORF1.

**S2 Fig.**
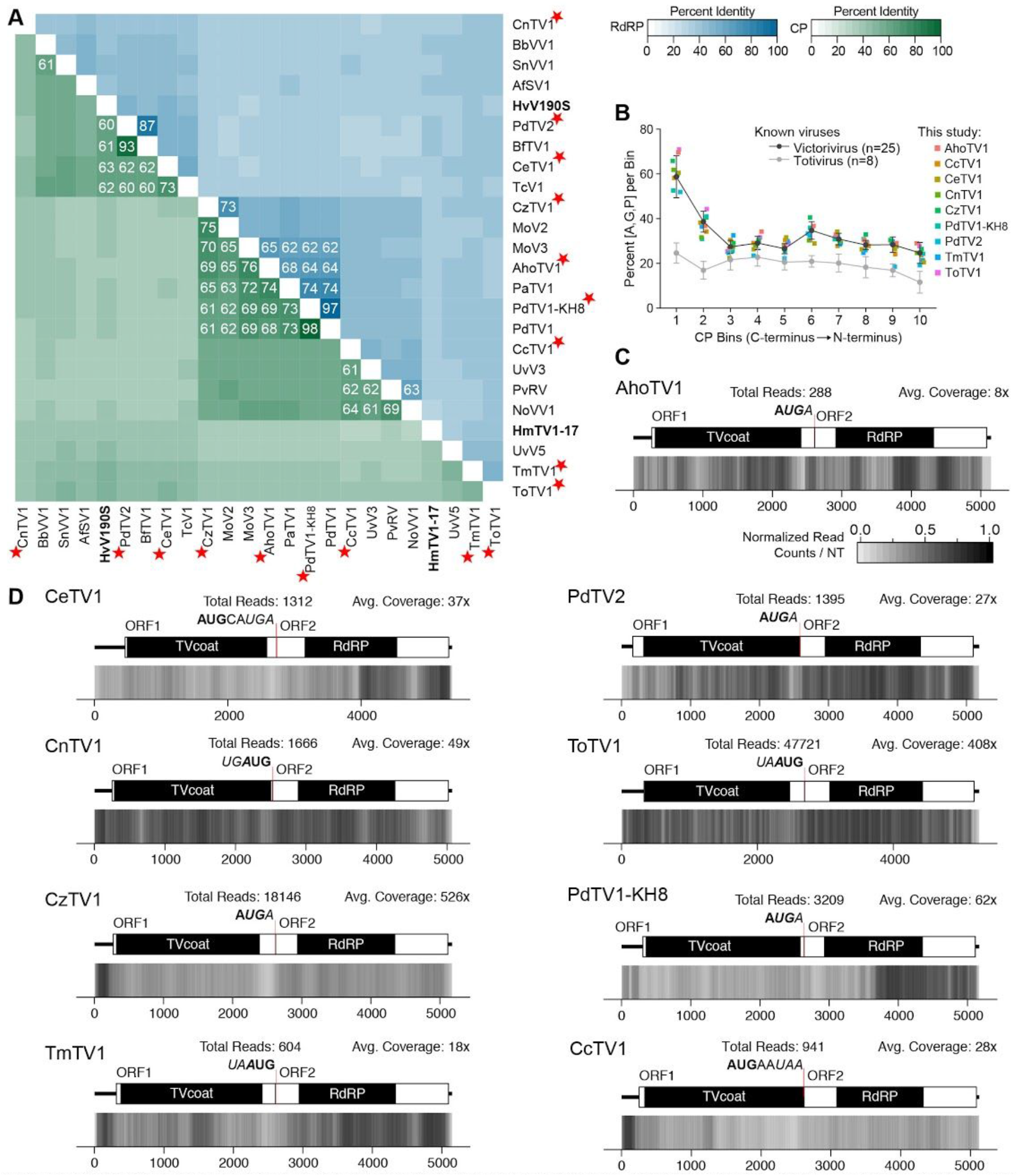
Analyses of identified viral genomes belonging to the *Victorivirus* genus within the *Totiviridae* family. (A) Percent identity matrix, generated by Clustal-Omega 2.1, comparing identified viral sequences to select members of the genus *Victorivirus.* The top half of the matrix, in the blue scale, is the percent identity of RdRP protein sequences and the bottom half of the matrix, in the green scale, is the percent identity of CP protein sequences. For sake of clarity, the 100% identity along the diagonal has been removed and outlined stars are used to note junction point for the row and column of the viruses identified. Where the identified viral sequences have significant similarity to known viral sequences is indicated, specifically RdRP or CP sequences with similarity >60%. Virus names in bold are ICTV-recognized species within the genus. (B) Percent of alanine, glycine, and proline (A,G,P) amino acids in the CP coding sequence. The average values were calculated for twenty-five sequences from known victoriviruses and ten sequences from known totiviruses, shown as dark and light grey lines respectively. The nine putative victoriviruses identified in this study were individually analyzed and plotted. (C and D) Density plot of read counts per nucleotide across the viral genome for the nine viruses identified. Graphical depiction of the genome structure, predicted ORFs, and putative motifs are illustrated on top and a heatmap of normalized read counts is on the bottom. Reads are normalized to a 0 −1 scale, where the maximum read count for a given genome equals 1. Total reads mapped to the virus genome and average read coverage are noted for each virus. The overlap between the CP stop codon (italics) and RdRP start codon (bolded) is also indicated. Red stars indicate viruses identified in this study. RdRP: RNA-dependent RNA Polymerase; CP: Coat Protein; ORF: Open Reading Frame.

A characteristic of the coat protein sequences of victoriviruses is an abundance of alanine, glycine, and proline (A,G,P) residues, particularly towards the C-terminal end. An analysis of between the CP protein sequences from vicŧorivirus species and totivirus species illustrates a generally elevated percentage of these three amino acids across the entire CP coding sequence of victoriviruses in comparison to totiviruses, with a notable increase observed in the two bins closest to the C-terminus of the protein (S2B Fig). The same pattern is also observed for all nine proposed vicŧorivirus sequences identified in this study.

These analyses demonstrate that these nine sequences share conserved characteristics found in members of the genus *Vicŧorivirus.* Criteria for species demarcation within this genus are fungal host species and percent identity of the RdRP and CP amino acid sequences [17], with a cutoff of 60% identity for both. The CP and RdRP sequences of CnTV1, TmTV1, and ToTV1 satisfy both the sequence-based criteria of < 60% identity and unique fungal host criteria to be characterized as new species (S2A Fig).

For a growing number of recently published sequences with strong phylogenetic ties to victoriviruses, the cutoff of 60% identity for the RdRP and/or CP is not satisfied, only the criteria of a differing fungal host (S2A Fig). For five of the six other Victoriviruses identified in this study, a 60% pairwise identity would not distinguish between viral sequences, only the host-based criteria. The sixth virus, PdTV1-KH8, shares 97% (RdRP) and 98% (CP) identity with the already published sequence for *Penicillium digitatum virus 1* (PdV1) [29]thus neither the sequence-based nor host-based criteria are met. Notably, PdTV2 and PdTV1-KH8 were both identified from the same data sample, however they share only 33.5% identity. The coincident infection of *P. dignitatum* KH8 by two victoriviruses is not unprecedented: *Sphaeropsis sapinea RNA virus 1* and *Sphaeropsis sapinea RNA virus 2* were isolated from the same host and share less than 60% similarity [28].

Clearly a precise sequence-based cut-off to distinguish between mycoviral species has not yet been identified. Until further biological studies to query the host range of these similar viruses can be performed, we would recommend more stringent values for pairwise percent identities for species demarcation within the *Vicŧorivirus* genus: 80% for CP sequences and 90% for RdRP sequences in addition to the identity of the fungal host. Following this recommendation, the eight viruses, CnTV1, TdTV2, CeTV1, CcTV1, CzTV1, AspTV1, TmTV1, and ToTV1, would be identified as new viral species within the *Victorivius* genus. To our knowledge, this is the first mycovirus described for *C. navitas, C. eremochloae, C. zoysiae, C. caudatum, Thelebolus microsporus*, and *Tolypocladium ophioglossoides.*

### Partitiviridae

Five genera have been assigned to the family *Partitiviridae*, along with fifteen viruses unassigned to a genus [30]. Genome structure of this family requires two distinct genome segments, each containing a single ORF: dsRNA1 and dsRNA2. The RdRP is found on dsRNA1 while the CP is found on dsRNA2, and highly conserved sequences can be found in the 5’UTR of both viral genome segments [30]. To date, viruses isolated from fungi have been classified into one of three of the *Partitiviridae* genera: *Alphapartitivirus, Betapartitiviruse*, and *Gammapartitivirus.* In this study, sixteen viruses were identified that clearly group among the known partitiviruses (Fig 2).

#### Alphapartitivirus

Five viruses identified in this study phylogenetically group within the *Alphapartitivirus* genus: *Clohesyomyces aquaticus partitivirus 1* (CaPV1) (PRJNA372822; [31]), *Grosmannia clavigera partitivirus 1* (GcPV1) (PRJNA184372; [32]), *Drechslerella stenobrocha partitivirus 1* (DsPV1) (; PRJNA236481[33]), *Gaeumannomyces tritici partitivirus* 1 and 2 (GtPV1, GtPV2) (PRJNA268052; [34]). To our knowledge, the first three are the first viruses identified in these fungal species while the two partitiviruses in *G. tritici* are the first partitiviruses from this fungus. As with other members of this genus, the genomic segment encoding the RdRP is approximately 1.9 to 2.0 kb while the genomic segment encoding the CP is between 1.7 and 1.9 kb (Table 1). Alignment of the RNA-seq reads to the viral genome sequences reveals average coverage per nucleotide ranging from 48x to 28,101x (S3B Fig). In general, the two dsRNA segments do not appear to have similar coverage per nucleotide across the segments, and in all but one instance, the dsRNA2 segment has the higher average coverage. For the two viruses isolated from *G. tritici*, 23x more reads align to GtPV1 than GtPV2 (S3B Fig).

**S3 Fig.**
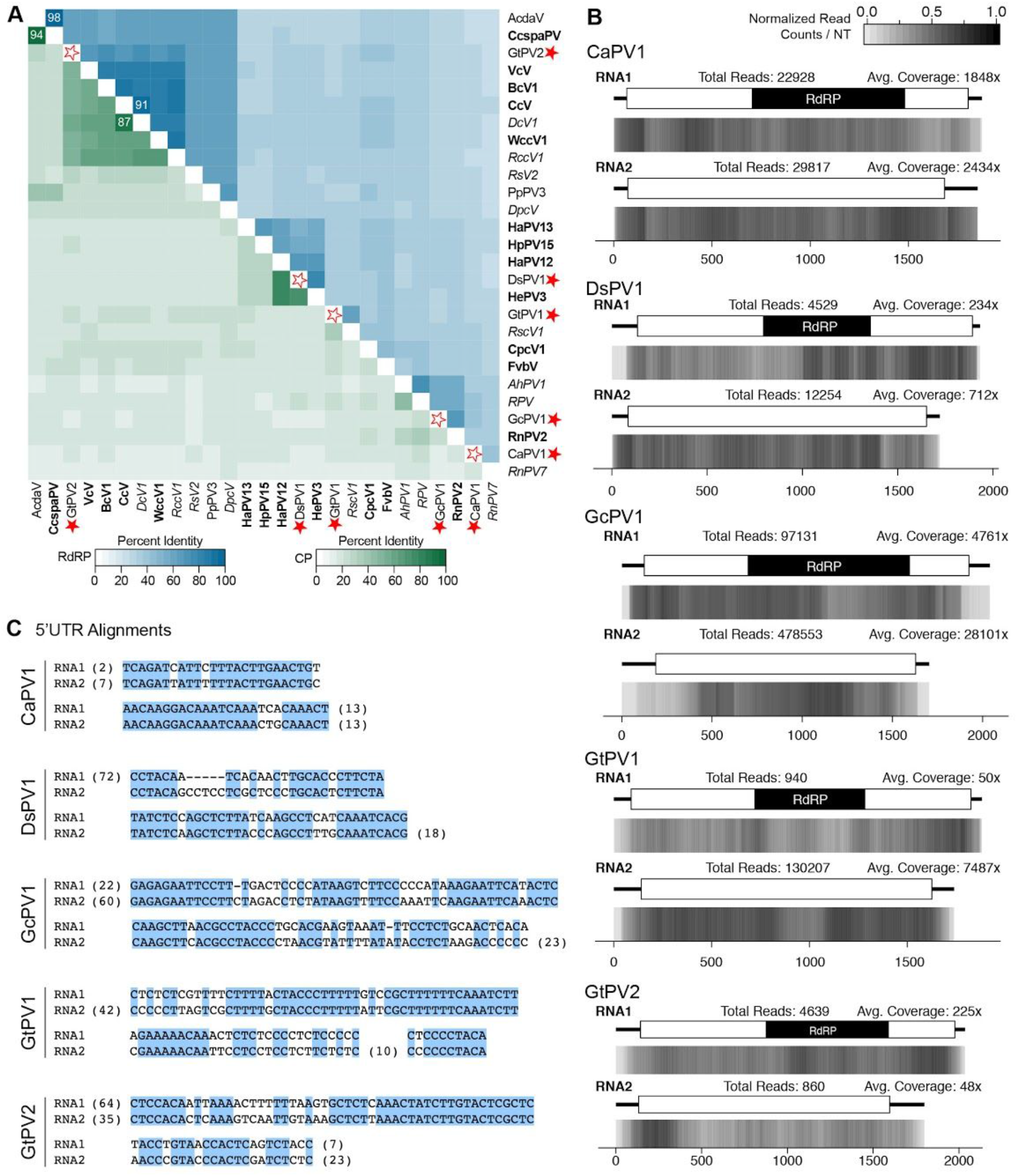
Analyses of viral genomes for members of the genus *Alphapartitivirus.* (A) Percent identity matrix generated by Clustal-Omega 2.1 of viruses identified in this study and select alphapartitiviruses. The top half of the matrix, in the blue scale, is the percent identity of RdRP protein sequences and the bottom half of the matrix, in the green scale, is the percent identity of CP protein sequences. For sake of clarity, the 100% identity along the diagonal has been removed and outlined stars are used to note the junction for the row and column of the viruses identified. Identities above the species demarcation are noted: ž90% for RdRP and >80% for CP. Names in bold are ICTV-member species, while names in italics are related, unclassified viruses. (B) Density plot of read counts per nucleotide across the viral genome for the five alphapartitiviruses identified in this study. Graphical depiction of the genome structures of RNA1 and RNA2, the predicted ORFs and RdRP motif are on illustrated above the heatmap of normalized read counts. Reads are normalized to a 0 – 1 scale, where the maximum read count for a given genome equals 1. Total reads mapped to the virus genome and average read coverage are noted for each virus. (C) Alignment of 5’UTRs of RNA1 and RNA2 genome segments. Residues not shown in alignment are in parentheses. Blue shading indicates regions of 100% identity. Red stars note the viruses identified in this study. RdRP: RNA-dependent RNA Polymerase; CP: Coat Protein.

Multiple sequence alignments of the 5’UTR regions of the two RNA genomic sequences for each identified viral genome revealed highly conserved stretches of nucleotides specific to each virus (S3C Fig). As dsRNA satellite sequences have been previously described for the alphapartitivirus *Cherry chlorotic rusty spot associated partitivirus* [35] the conserved sequences present in dsRNA1 and dsRNA2 were used to query the remaining Trinity contigs, however no putative satellite RNAs were identified.

The criteria for species demarcation within the genus *Alphapartitivirus* are ≤90% identity in the RdRP protein sequence and ≤80% identity in the CP protein sequence. All five viral genomes identified in this study satisfy these criteria and as such are new species of the genus *Alphapartitivirus* (S3A Fig).

#### Betapartitivirus

Two virus genomes were identified as members of the genus *Betapartitivirus*: *Trichoderma citrínoviride partitivirus 1* (TcPV1) (PRJNA304029; [36]) and *Fusarium poae partitivirus 1-2516* (FpPV1-2516) (PRJNA319914; [37]). To our knowledge, this is the first partitivirus identified from *T. citrínoviride* while FpPV1-2516 is the third instance of a species of betapartitivirus described to date from the fungus *F. poae.*

The RdRP coding sequence from FpPV1-2516 shares 99.9% and 97.9% identity with FpV1 (F. *poae* strain A11) and FpV1 (F. *poae* strain 240374) respectively (S4A Fig). Similarly, the CP sequence of FpPV1-2516 shares 99.8% identity with both of the FpV1-CP sequences. Despite the high sequence similarity observed, these three viruses appear to originate from different locales. FpV1-240374 was published in 2016 following deep RNA-sequencing of the fungal strain F. *poae* MAFF 240374, a fungus isolated from a *Triticum aestivum* spikelet in Japan in 1991 [38]. FpV1 (strain A11) was identified in 1998 from the strain F. *poae* A-11 isolated from wheat grain in Hungary [39]. Finally, FpPV1-2516, identified in this study, was found in the RNA-seq data from F. *poae* strain 2516, a fungus isolated in Belgium from wheat [37]. While FpPV1-2516 is not a unique viral species, it does have a unique genome sequence and unique history from the other two known viruses and as such, has been deposited at Genbank (Table 1). A pairwise percent identity analysis of TcPV1 with other betapartitiviruses reveals that this is a unique sequence since identity values for both the RdRP and CP sequences are below the established thresholds for species demarcation of 90% identity and 80% respectively (S4A Fig) [30].

A sequence alignment of the 5’UTR sequences of the two genomic segments of TcPV1 reveals stretches of highly conserved nucleotides, characteristic of partitiviral genomes (S4C Fig). Additionally, the two genomic segments are both 2.3 kb (Table 1), which is within the described range for betapartitiviruses of 2.1 to 2.4 kb [30]. As observed with the majority of the alphapartitiviruses described above, the average coverage per nucleotide for the genomic segments of FpPV1-2516 and TcPV1 was higher for the dsRNA2 segment that encodes the CP than the RdRP segment (S4B Fig).

**S4 Fig.**
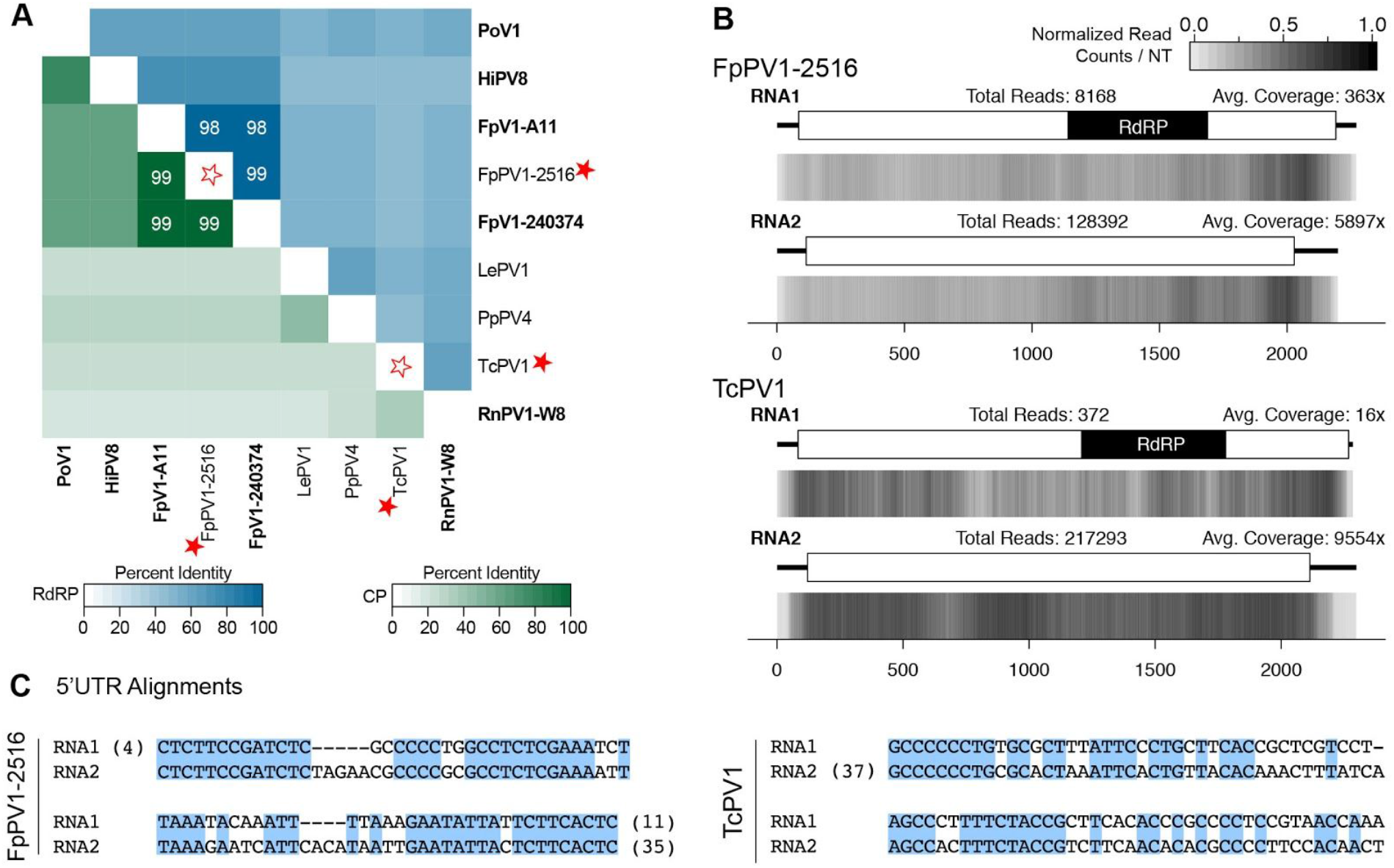
Analyses of Identified viral genomes belonging to the genus *Betapartitivirus.* (A) Percent identity matrix, generated by Clustal-Omega 2.1, between identified viruses and select betapartitiviruses. The top half of the matrix, in the blue scale, is the percent identity of RdRP protein sequences and the bottom half of the matrix, in the green scale, is the percent identity of CP protein sequences. For sake of clarity, the 100% identity along the diagonal has been removed and outlined stars are used to note junction point for the row and column of the viruses identified. Where the percent similarity is above the cutoffs for species demarcation is noted, specifically >90% for RdRP sequences and >80% for CP sequences. Bolded names are ICTV-recognized member species. Full names and accession numbers for protein sequences are in Table S2. (B) Density plot of read counts per nucleotide across the viral genome for the two *Betapartitiviruses.* Graphical depiction of the genome structure of the two RNAs, the predicted ORFs, and the RdRP motif are along the top and a heatmap of normalized read counts is on the bottom. Read counts are normalized to a 0 −1 scale, where the maximum read count for a given genome equals 1. Total reads mapped to the virus genome and average read coverage are noted for each virus. (C) Alignment of 5’UTR sequences from the RNA1 and RNA2 genome segments. Blue boxes highlight conserved nucleotides and numbers in parentheses are residues not shown. Viruses identified in this study are noted with a red star. RdRP: RNA-dependent RNA Polymerase; CP: Coat Protein.

#### Gammapartitivirus

Five viruses identified in this analysis grouped with members of the genus *Gammapartitivirus* (Fig 2): *Penicillium digitatum partitivirus 1-HSF6* (PdPV1-HSF6) (PRJNA352307; unpublished) and *Pseudogymnoascus destructans partitivirus 1-20631* (PdPV1-20631) (PRJNA66121; unpublished) along with *Magnaporthe grisea partitivirus 1* (MgPV1) (PRJNA269089; [40]), *Thelebolus microsporus partitivirus 1* (TmPV1) (PRJNA372886; unpublished), and *Phyllosticta citriasiana partitivirus 1* (PcPV1) (PRJNA250394; unpublished). The latter three viruses are, to our knowledge, the first mycoviruses described for these fungal species and, as described below, satisfy the criteria to be identified as new viral species.

The dsRNA1 genomic segment encoding the RdRP for all five viruses is approximately 1.7 kb while the dsRNA2 genomic segment encoding the CP is between 1.5 and 1.6 bp (Table 1) both within the expected range of other known members of the genus *Gammapartitivirus* of 1.6 to 1.8 kb for dsRNA1 and 1.4 to 1.6 kb for dsRNA2 [30]. A multiple sequence alignment analysis of the 5’UTR sequences from the two genomic segments of each virus revealed stretches of conserved nucleotides within four of the five viruses; PcPV1 was the exception, however the 5’UTR sequence of dsRNA1 is only 17 nt (S5C Fig). Attempts to extend this sequence by manually querying the RNA-seq data were unsuccessful.

Total reads aligned and average coverage per nucleotide were calculated following alignment of the RNA-seq reads to the viral genome sequences. In general, the average coverage per nucleotide for dsRNA1 is similar to that of dsRNA2 (S5B Fig), a departure from the alignment results for the alphapartitiviruses and betapartitiviruses described above.

A pairwise percent identity analysis compared the RdRP coding sequences and the CP coding sequences for the five viruses to other sequences within the genus *Gammapartitivirus* (S5A Fig). The current criteria for species demarcation is 90% for the RdRP sequence and 80% for the CP sequence [30]. Genic and protein sequences from PdPV1-20631 share 100% identity with the virus PdPVpa from *Pseudogymnoascus destructans* 80251, although differences were observed in the length of the 5’ and 3’ UTR sequences. The RdRP and CP protein sequences of PdPV1-HSF6, isolated from *Penicillium digitatum*, shares 95% identity with the proteins from *Peniciilium stoioniferum virus S* (PsVS). Due to the high sequence similarity and the relative close relationship between the two fungal hosts from the *Penicillium* genus, this virus sequence does not meet the criteria to qualify as a new viral species. Nonetheless, these viral sequences have been deposited in GenBank due to the differences within the nucleotide sequences from published viral genomes and the identity of the host fungal strain. The RdRP and CP protein sequences from MgPV1, PcPV1, and TmPV1 do not share significant identity with known gammaparitivirus sequences and are isolated from new fungal hosts. As such, these three viruses are characterized as new viral species.

**S5 Fig.**
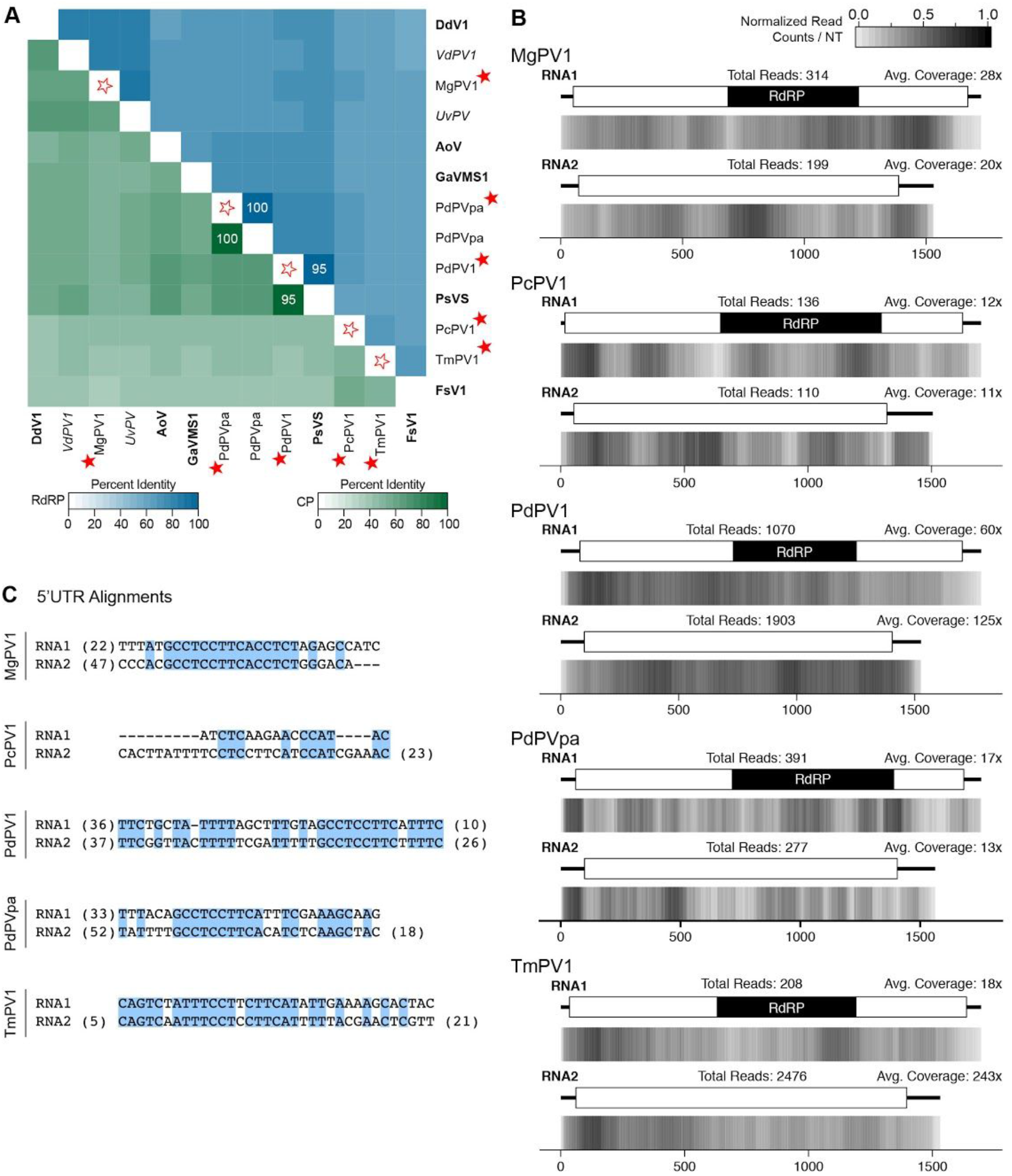
Analyses of identified viral genomes belonging to the genus *Gammapartitivirus.* (A) Percent identity matrix, generated by Clustal-Omega 2.1, between identified viruses and select gammapartitiviruses. The top half of the matrix, in the blue scale, is the percent identity of RdRP protein sequences and the bottom half of the matrix, in the green scale, is the percent identity of CP protein sequences. For sake of clarity, the 100% identity along the diagonal has been removed and outlined stars are used to note junction point for the row and column of the viruses identified. Where a sequences has a value greater than the species demarcation cutoff is noted, specifically >90% for RdRP and >80% for CP sequences. Bolded names are ICTV-recognized member species while italicized names are related, but unclassified viruses. Full names and accession numbers for protein sequences are in Table S2. (B) Density plot of read counts per nucleotide across the viral genome for the five gammapartitiviruses. Graphical depiction of the genome structure of the RNA segments, predicted ORFs, and RdRP motif are on top and a heatmap of normalized read counts is on the bottom. Read counts are normalized to a 0 −1 scale, where the maximum read count for a given genome equals 1. Total reads mapped to the virus genome and average read coverage are noted for each virus. (C) Alignment of 5’UTR sequences of RNA1 and RNA2 for the identified viral sequences. Conserved nucleotides are highlighted with a blue box and number of nucleotides not shown in the alignment are noted in parentheses. Viruses identified in this study are noted with a red star. RdRP: RIMA-dependent RNA Polymerase; CP: Coat Protein.

#### New Genus: Epsilonpartitivirus

Two viral genomes, *Coiletotrichum eremochloae partiŧivirus 1* (CePV1) (PRJNA262412; unpublished) and *Penicillium aurantiogriseum partiti-like virus* (PaPIV) (PRJNA369604; [41]), were identified that are phylogenetically related to an emerging group of partitiviruses that are separate from, but related to, the genus *Gammapartitivirus* (Fig 2); this group has been tentatively named *Epsilonpartitivirus* [41]. Identification of the PaPIV sequences was expected as this sample was RNA-seq data from *Cryphonectria parasitica* infected with PaPIV, previously isolated from *P. aurantiogriseum* by the same researchers [41]. To distinguish this viral genome from that isolated from the original host, the virus has been named PaPV1-cp. Also, the slight modification of the virus name, from partiti-like to partitivirus, demonstrates a recent improved understanding that this is a partitivirus following the identification of a RNA segment encoding a putative CP [41].

Members of this proposed genus have been identified from both fungal and invertebrate hosts, although currently the viral sequences form two sister clades separated by kingdom (Fig 2). Genome sizes of the dsRNA segments of putative epsilonpartitiviruses are from 1.7 to 1.9 kb for dsRNA1 and 1.5 to 1.8 kb for dsRNA2 (Table 1). Using a pairwise comparison analysis, a value of 85% identity for the RdRP protein sequence and 75% for the CP sequence would currently be appropriately stringent for species demarcation of viruses isolated from different fungal hosts (Fig 5A).

**Fig 5.**
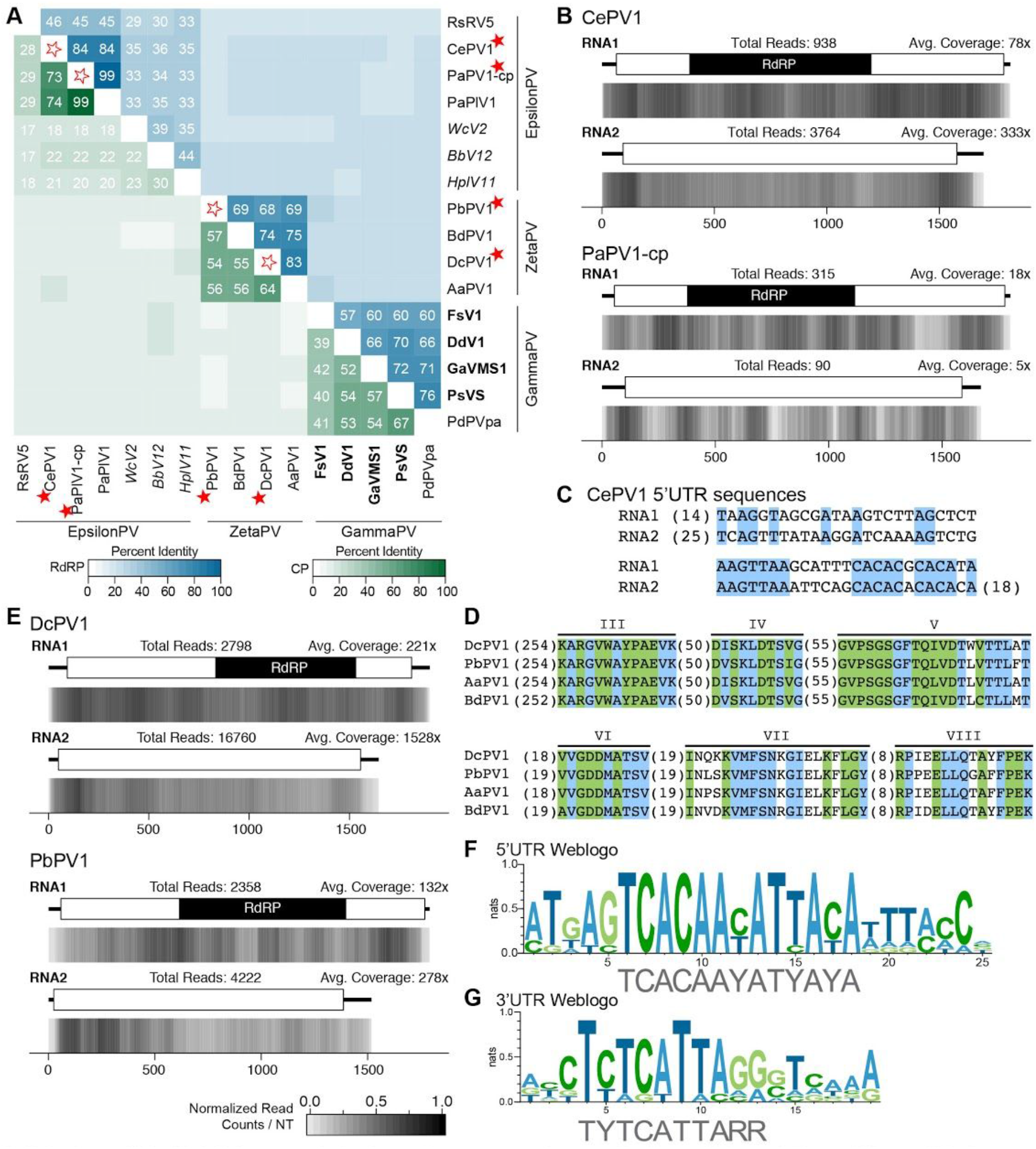
Analyses of identified viral genomes belonging to the two proposed genera, *Epsilonpartitivirus* and *Zetapartĩtivirus*, within the *Partitiviridae* family. Percent identity matrix, generated by Clustal-Omega 2.1, between members of the two new groups and *gammapartitiviruses*, the most closely related, ICTV-recognized genus. The top half of the matrix, in the blue scale, is the percent identity of RdRP protein sequences and the bottom half of the matrix, in the green scale, is the percent identity of CP protein sequences. For sake of clarity, the 100% identity along the diagonal has been removed and outlined stars are used to note junction point for the row and column of the viruses identified. Two cutoff values were applied and as such only percent identities >29% among RdRP sequences and >17% among CP sequences are indicated. Virus names in italics refer to viruses isolated from insect, not fungal, hosts. Bolded virus names are member species recognized by ICTV. (B and E) Density plot of read counts per nucleotide across the viral genome for the two epsilonparitiviruses (B) and two zetapartitiviruses (E). Graphical depiction of the genome structure, predicted ORFs, and RdRP motif are along the top and a heatmap of normalized read counts is on the bottom. Read counts are normalized to a 0 −1 scale, where the maximum read count for a given genome equals 1; heatmap scale found in (E) applies also to (B). Total reads mapped to the virus genome and average read coverage are noted for each virus. (C) Alignment of the 5’ UTR sequences from the two genomic segments of the epsilonpartitivirus CePV1. Conserved nucleotides are highlighted in blue and numbers in parentheses indicate nucleotides not shown. (D) Alignment of the six conserved motifs within the RdRP protein sequence of all four proposed zetapartitiviruses. Amino acids highlighted in blue are only common amon zetapartitiviruses while residues in green are shared with gammapartitiv?ruses. (F and G) Weblogos generated from alignments of the (F) 5’UTR and (G) 3’UTR sequences of both RNA genome segments for all four viruses in the Zetapartitivirus genus. Red stars indicate viruses identified in this study. RdRP: RNA-dependent RNA Polymerase; CP: Coat Protein.

Observed here, sequences isolated from the PaPIV1-infected *C. parasitica* sequencing sample contain a number of SNPs in comparison to the source PaPIV sequence; these changes may reflect an adaptive response by the virus to a new fungal host [41]. A positive effect on fungal growth is observed whereby the virus confers a positive effect on fungal growth of *C. parasitica* in the presence of 2% salts that has not yet been confirmed in the original host *P. aurantiogriseum.* Read coverage along the viral genome segments isolated from *C. parasitica* is on average low, 18x for dsRNA1 and 5x for dsRNA1 (Fig 5B), similar to the published report [41].

Both of the viral genome segments of CePV1 share the greatest percent identity with PaPIV1, but below the threshold suggested above, at 84% and 73% for the RdRP and CP respectively (Fig 5A). Similar to other partitiviruses, a multiple sequence alignment of the 5’UTR sequences of the two genome segments of CePV1 reveals regions of sequence conservation (Fig 5C).

Initially a mycoviral sequence was isolated from a *Colletotrichum navitas* sequencing sample (PRJNA262223; unpublished); however, other than small variances within the 5’ and 3’ UTRs, the nucleotide sequence is identical to that of CePV1. It is possible that two different, closely related, fungi are infected with the same virus, however as noted above with respect to the PaPIV1-infected *C. parasitica*, SNPs can quickly appear within a viral genome sequence when in a new host. Therefore further analyses were performed to determine the likelihood of a CePV1-like sequence hosted by *C. navitas.* First it was determined that, after controlling for the unaligned reads pool size, 2.6x more viral reads were present in the *C. eremochloae* sample than the *C. navitas* sample. Second, to identify possible cross-contamination between the two species within the sequencing data, non-virus-derived reads were reciprocally aligned to the other fungal genome. Thus reads from *C. navitas* were aligned to the *C. eremochloae* genome, and vice versa. Here we observed that 5% of reads from *C. eremochloae* aligned to the *C. navitas* genome while 65% of reads in the *C. navitas* pool aligned to the *C. eremochloae* genome. Evaluating both of these results, we propose that the likely host of this viral sequence is *C. eremochloae* and the identification of the same sequence within the *C. navitas* sample is likely due to cross contamination from the *C. eremochloae* sample. Despite the possible ambiguity in fungal host for this viral sequence, we recommend its inclusion in the recently suggested genus *Epsilonpartitivirus* as a new viral species.

#### New Genus: Zetapartitivrus

The final two viruses identified within the *Partitiviridae* branch of the phylogenetic tree (Fig 2) are *Penicillium brasilianum partitivirus 1* (PbPV1) (PRJEB7514; [42]) and *Delitschia confertaspora partitivirus 1* (DcPV1) (PRJNA250740; unpublished). Both viruses, the first to be described in these fungal species, phylogenetically group with *Alternaria alternata partitivirus 1* (AaPV1) [43] and *Botryosphaeria dothidea partitivirus 1* (BdPV1) [44], a distinct clade within the *Partitiviridae* that is related to, but separate from, the genus *Gammapartitivirus* (Fig 2).

These four viruses share many sequence-based similarities. A 13-nt highly conserved sequence (TCACAAYATYAYA) is present in the 5’UTR sequence of both dsRNA genome segments from all four viruses (Fig 5F). Additionally, a 10-nt conserved sequence (TYTCATTARR) is found within the 3’UTR sequence of both dsRNA segments of all four viruses (Fig 5G). While these four viruses contain the six conserved motifs common to RdRP proteins from other members of the *Partitiviridae*, including the GDD triad in motif VI, there are distinct differences that are shared that distinguish these viruses from the existing genera (Fig 5D).

Further, the range of lengths for the RdRP protein, from 569 to 572 aa, and CP protein, from 453 to 501 aa (Table 1), are unique ranges among the currently described genera [30]. A pairwise percent identity analysis of the RdRP and CP sequences from the four viruses suggests a threshold for species demarcation at 90% for the RdRP sequence and 80% for the CP sequence, similar to other genera within the family (Fig 5A).

Alignment of the RNA-seq reads to the viral genomes demonstrates that dsRNA2 has a higher average coverage per nucleotide for both viruses, with approximately 2x more coverage observed between the segments of PbPV1 and 7x between the segments of DcPV1 (Fig 5E).

We recommend that DcPV1 and PbPV1, along with AaPV1 and BdPV1 belong to a new genus within the family *Partitivirus*, called *Zetapartitivirus.* The genomic segment and protein sequence lengths are consistent and unique within the *Partitivirus* family, and the RdRP protein sequences form a distinct clade that is separate from the closest genus, *Gammapartitivirus* (Fig 2).

### Chrysoviridae

To date, nine species have been classified as belonging to the Chrysoviridae family of dsRNA viruses that infect either ascomycota or basidiomycota fungi [45]. Members of this family have multipartite genomes, consisting of three to five segments. Here we identify two viral genomes with sequence similarity to Chrysoviruses: *Beauveria bassiana chrysovirus 1* (BbCV1) ([46]PRJNA306902) and *Penicillium raistrickii chrysovirus 1* (PrCV1) (unpublished; PRJNA250734) (Fig 2).

Phylogenetically, chrysoviruses are grouped into two distinct clusters, where cluster 1 contains members with either three or four segmented genomes, while cluster 2 contains viruses with either four or five segments [45]. The two viral sequences identified in this study, BbCV1 and PrCV1, belong with cluster 1 chrysoviruses; both genomes are comprised of four genomic segments and the RdRP sequences phylogenetically group with the ICTV-recognized members within cluster 1, *Isaria javanica chrysovirus 1* (IjCV1) [47] and *Penicillium chrysogenum virus* (PcCV1) [48] (Fig 2). Further, as detailed below, we recommend that these are both new viral species within the genus *Chrysovirus.*

To our knowledge, these are the first chrysoviruses identified for both of these fungal hosts, and indeed the first virus from *P. raistrickii* described, thus satisfying the host-based criteria for species demarcation. Following identification of the RdRP-encoding contig, the identity of the other three segments was determined via a BLASTP search using protein sequences of known chrysoviruses against the predicted protein sequences of the Trinity contigs. The combined length of dsRNAs for both BbCV1 and PrCV1 are 12.46 and 12.56 kb respectively (Table 1), within the observed range of other 11.5 to 12.8 kb of other chrysoviruses [45]. In general, chrysovirus genome segments are numbered by size, however segment 4 of PrCV1 is slightly longer than segment 3 (Table 1), similar to segment 4 of *Amasya cherry disease associated chrysovirus* (AcdaCV) which is also larger than its respective segment 3 [49]. A BLASTP analysis confirms that segment 4 of PrCV1 best matches the segment 4 of *Penicillium chrysogenum virus* and thus is numbered accordingly. An alignment of the RNA-seq reads to the viral genome segments reveals uniform average coverage per nucleotide for the four genomic segment of PrCV1 (from 15x to 32x) and slightly more variable average coverage of BbCV1 segments, ranging from 10x to 72x (S6D Fig). The dsRNA genomic segment lengths therefore also satisfy the criteria for species demarcation within the genus *Chrysovirus.*

The 5’UTR of chrysoviruses is generally between 140 and 400 nt in length and contains two regions of high sequence conservation, Box 1 and Box 2 [45], although a number of recently characterized viruses have shorter 5’UTRs, including *Cryphonectria nitschkei chrysovirus 1* (CnCV1) which has 82 nt 5’UTR for segment 1 [50] and AcdaCV which has 5’UTR lengths of 87, 96, 95, and 106 bp [49]. With the exception of the 5’UTR of BbCV1-segment 1 (106 nt), the 5’UTR of all segments discovered for BbCV1 and PrCV1 fall within the published range (S6B Fig and S6C Fig). A highly conserved region is present within area corresponding to “Box 1” within the 5’UTR of the four genome segments of BbCV1 (S6B Fig, top panel) and the expected “CAA” sequence is also observed in all sequences (S6B Fig, bottom panel). Similarly both conserved regions are observed within the 5’UTR of PrCV1 (S6C Fig). Thus the size and composition of the 5’UTR sequences of BbCV1 and PrCV1 also satisfy the criteria for chrysovirus classification.

The final criteria considered for species demarcation is the percent identity of the RdRP and CP protein sequences relative to other members within the genus, where the thresholds are 70% and 53% respectively [45]. BbCV1 RdRP and CP share 83% and 69% identity with the related proteins of IjCV1, while PrCV1 RdRP and CP protein sequences share 88% and 79% identity with those of *Penicillium chrysogenum virus* (PcCV1) (S6A Fig). Since BbCV1 was isolated from a new fungal host, we propose classifying it as a new species of chrysovirus, while the relative relatedness between the hosts *P. raistrickii* and *P. chrysogenum* suggest that PrCV1 is not a new species of chrysovirus. Further identification of additional viruses plus biological characterization of host specificity will provide the needed insights to properly classify viruses within this family.

**S6 Fig.**
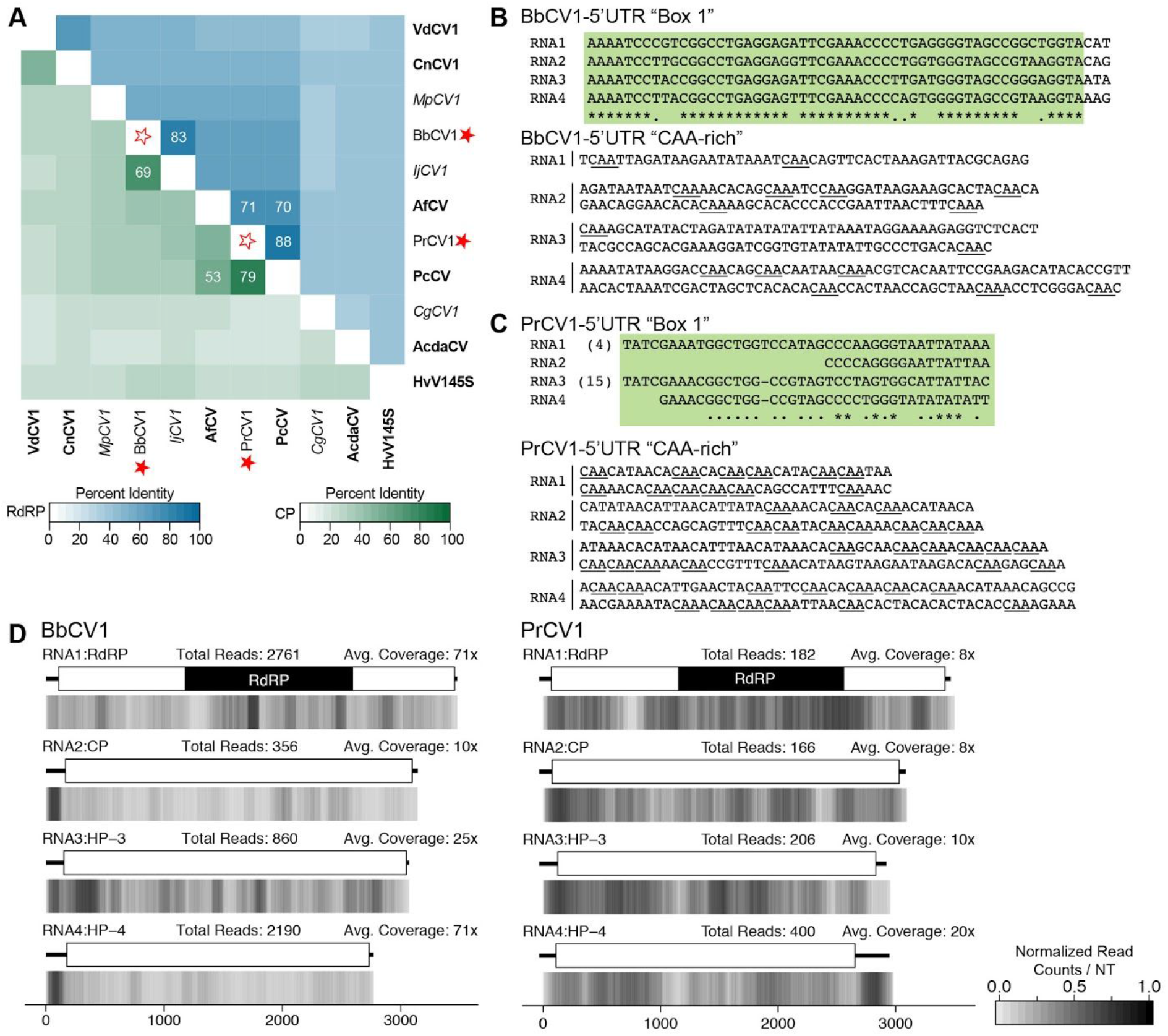
Analyses of viral genomes belonging to the *Chrysoviridae* family. (A) Percent identity matrix, generated by Clustal-Omega 2.1, versus select members of the *Chrysovirus* genus. The top half of the matrix, in the blue scale, is the percent identity of RdRP protein sequences and the bottom half of the matrix, in the green scale, is the percent identity of CP protein sequences. For sake of clarity, the 100% identity along the diagonal has been removed and outlined stars are used to note junction point for the row and column of the viruses identified. Values above 70% for RdRP sequences and 53% for CP sequences are indicated. ICTV-member and unclassified viruses are noted in bold and italics respectively. (B and C) Alignment of the 5’UTR sequences for the four genomic segments of BbCV1 (B) and PrCV1 (C). Two highly conserved regions are found in the 5’UTR; the first is a 40-75 nt region, highlighted in green, while the second region is rich in CAA repeats (underlined). (C) Density plot of read counts per nucleotide across the viral genome segments. Graphical depiction of the genome structure and predicted ORFs and motifs are on top and on the bottom is a heatmap of read counts normalized to a 0 −1 scale, where the maximum read count for a given genome equals 1. Total reads mapped to the virus genome and average read coverage are noted. Red stars indicate viruses identified in this study. RdRP: RNA-dependent RNA Polymerase; CP: Coat Protein; HP: Hypothetical Protein.

### Endornaviridae

Members of the *Endornaviridae* family of viruses have long, single segment, dsRNA genomes, generally encoding a single polyprotein. In addition to the RdRP domain and helicase domain (Viral helicase 1 or DEADx helicase) found in all endornaviruses, viral genomes may also encode a viral methyltransferase domain and/or a glycosyltransferase domain. Recently a two-genus split was proposed [51] where characteristics of clade I (*Alphaendornavirus)* include a plant-host, larger genomes (ranging from 13.8 to 17.5 kb), the presence of a UGT domain and a site-specific nick towards the 5’end of the genome sequence [52], while members of clade II (*Betaendornavirus*) have fungal-hosts and generally have genome sizes ranging from 9.5 to 11.4 kb, encode a viral methyltransferase domain but not a UGT domain, and lack the site-specific nick [51].

Here four new virus sequences were identified that group with known endornaviruses (Fig 6a), *Gyromitra esculenta endornavirus 1* (GeEV1) (PRJNA372840; unpublished) and *Morchella imporŧuna endornavirus* 1, 2, and 3 (MiEV1, MiEV2, MiEV3) (PRJNA372858; unpublished). These are the first viruses, to our knowledge, identified from both *G. esculenta* and the morel mushroom, *M. imporŧuna.*

**Fig 6.**
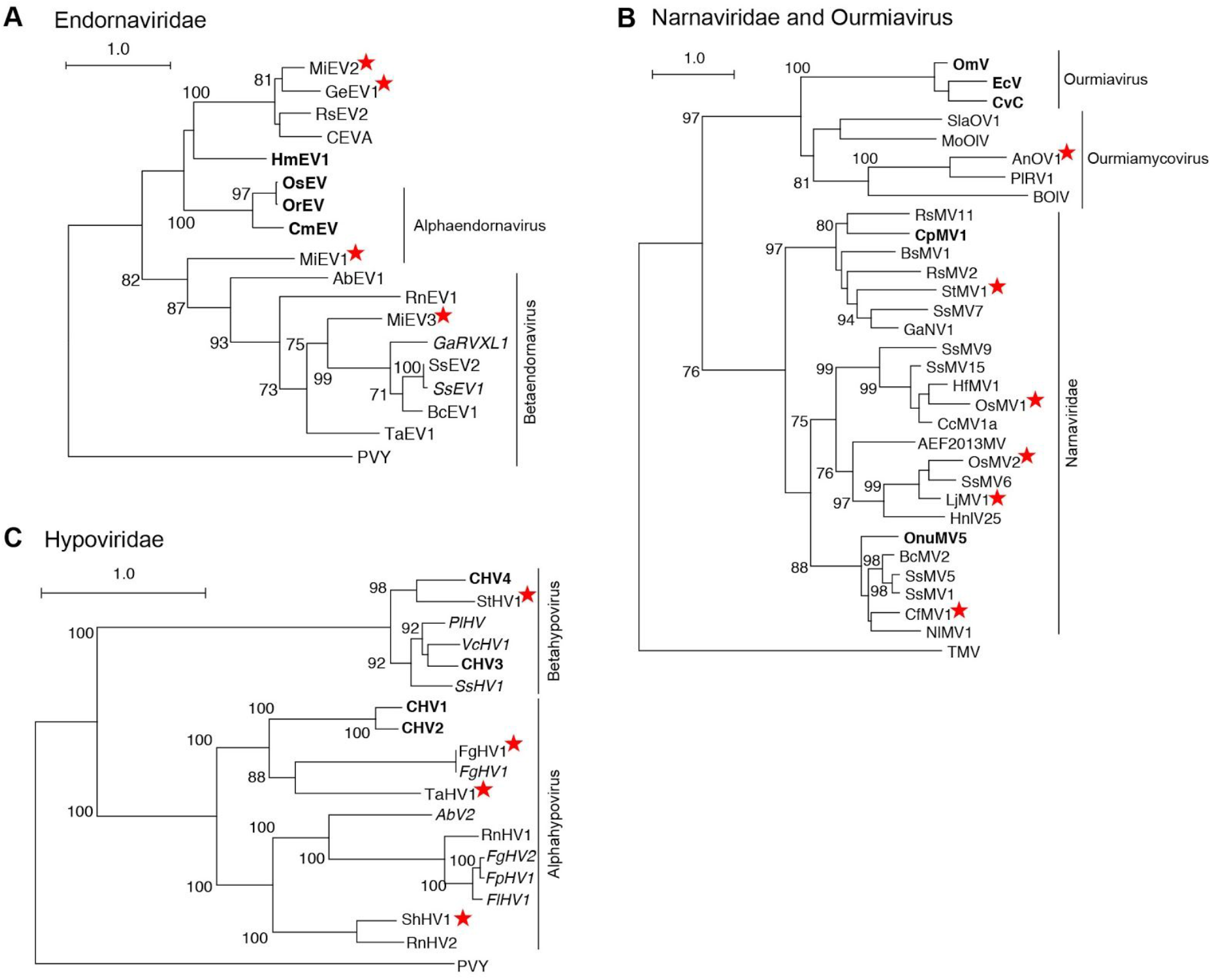
Phylograms based on RdRP motif sequences from viruses from the *Endornaviridae* family (A), *Narnaviridae* and *Ourmiaviruses* (B), and *Hypoviridae* family (C). Full names of viruses and sequences can be found in Table S2, including the outgroup, *Potato Virus Y* (PVY). Bolded names are ICTV-recognized member species while italicized names are related, but unclassified viruses. Full names and accession numbers for protein sequences used in the alignment are in Table S2. Scale bar on the phylogenetic tree represents 1.0 amino acid substitutions per site and numbers at the nodes indicate bootstrap support over 70% (1000 replicates). Viruses identified in this study are noted with a red star.

#### Alphaendornavirus

A phylogenetic analysis of the RdRP motif from related endornaviruses and the viruses identified in this study reveals MiEV2 and GeEV1 form a well supported clade with other fungal endornaviruses, including *Ceratobasidium endornavirus A* (CEVA) [53], and *Rhizoctonia solani endornavirus 2* (RsEV2) [54], that is related to, but separate from alphaendornaviruses isolated from plant hosts, including the type species *Oryza sativa endornavirus* (OsEV) [55] (Fig 6A).

The genome sizes of MiEV2 and GeEV1 are 15.3 and 14.5 kb respectively, well within the observed range of other *Alphaendornaviruses* (Table 1). Additionally, a conserved domain search within the protein coding sequences determined both viruses encode a viral helicase and an RdRP domain, but not a viral methyltransferase domain, nor a UGT domain (S7B Fig). This combination of protein domains appears to be shared among mycoviruses and not plant-hosted endornaviruses.

GeEV1 has a predicted AUG at position 5, resulting in a shorter 5’ UTR sequence than observed in other endornaviruses. The predicted start codon for MiEV2 however is at position 476, resulting in an unexpectedly long 5’UTR; as the entire sequence upstream of this codon is in-frame with the coding sequence, it is possible that this genome sequence is partially truncated and is missing the true AUG sequence. As manual analysis of the RNA-seq data did not result in an extension of the 5’ region of MiEV2, we recommend further analysis of the biological material to confirm the 5’ region of this virus. An alignment of RNA-seq reads to the two viral genomes reveals uniform coverage across both genome sequences, with a total of 4977 reads and 8171 reads respectively (S7B Fig).

**Fig 7.**
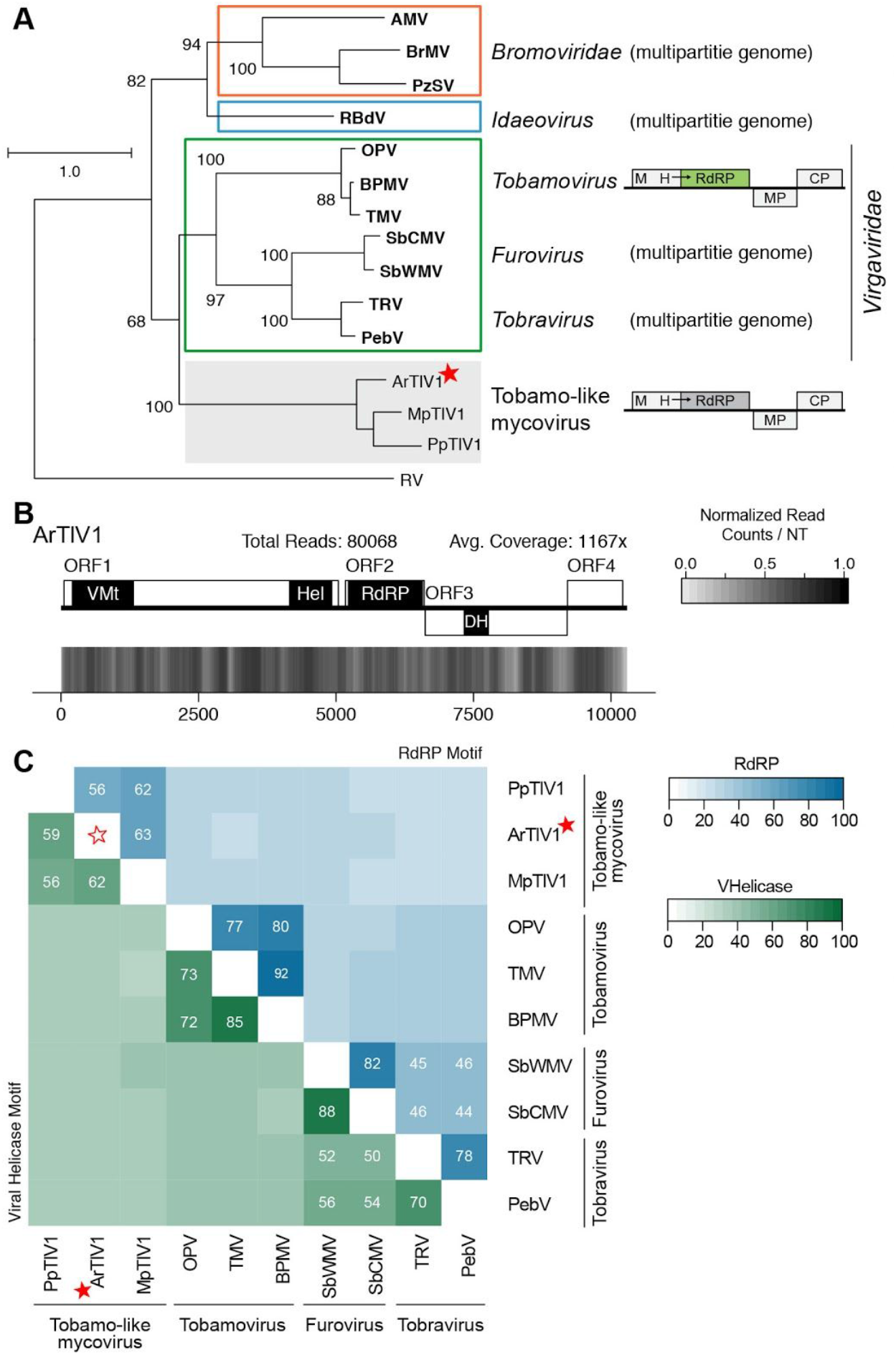
Analyses of *Acidomyces richmonden-sis tobamo-like virus 1* (ArTIV1) and related viruses. (A) Phylogram of the RdRP-motif for ArTIV1, related mycoviruses, and select members of the genus *Tobamovirus* and family *Bromoviridae*;, sequence from *Rubella Virus* (RV) serves as the outgroup. Bolded names are g ICTV-recognized member species. Full names and accession numbers for protein sequences used are in Table S2. Numbers at the nodes S^1^ indicate bootstrap support values ≥65% (1000 replicates). Scale bar on the phylogenetic tree represents 1.0 amino acid substitutions per site. Diagrammatic genome structures are to the right of the phylogram. (B) Density plot of read counts per nucleotide across the ArTIV1 viral genome. Graphical depiction of the genome structure, predicted ORFs, and encoded motifs are along the top and a heatmap of normalized read counts is on the bottom. Reads are normalized to a 0 – 1 scale, where the maximum read count equals 1. Total reads mapped to the virus genome and average read coverage are noted. (C) Percent identity matrix, generated by Clustal-Omega 2.1. The top half of the matrix, in the blue scale, is the percent identity of the RdRP motif sequence while and the bottom half of the matrix, in the green scale, is the percent identity of the residues from Viral Helicase motif. For sake of clarity, the 100% identity along the diagonal has been removed and outlined stars are used to note junction point for the row and column of ArTIV1. Cutoff values of 44% and 40% were applied to the RdRP and Helicase matrices respectively. Viruses identified in this study are noted with a red star. VMt: Viral Methyltransferase; VHel: Viral Helicase; RdRP: RNA-dependent RNA Polymerase; DH: DEXDc Helicase; ORF: Open Reading Frame.

The two criteria for species demarcation within the genus *Alphaendornavirus* are difference in host and an overall nucleotide sequence identity below 75%. Both viruses described here satisfy these criteria. Indeed, nucleotide identity among *Endornaviruses* is generally below 50%. An analysis of the protein sequences of the RdRP motif and the viral helicase motif reveal similar results, as sequences are between 39% and 67% identical for the RdRP sequences and 26% to 51% identical among the helicase sequences (S7A Fig). Together these analyses support the recognition of GeEV1 and MiEV2 as new species within the *Endornaviridae* family, and the genus *Alphaendornavirus* in particular.

**S7 Fig.**
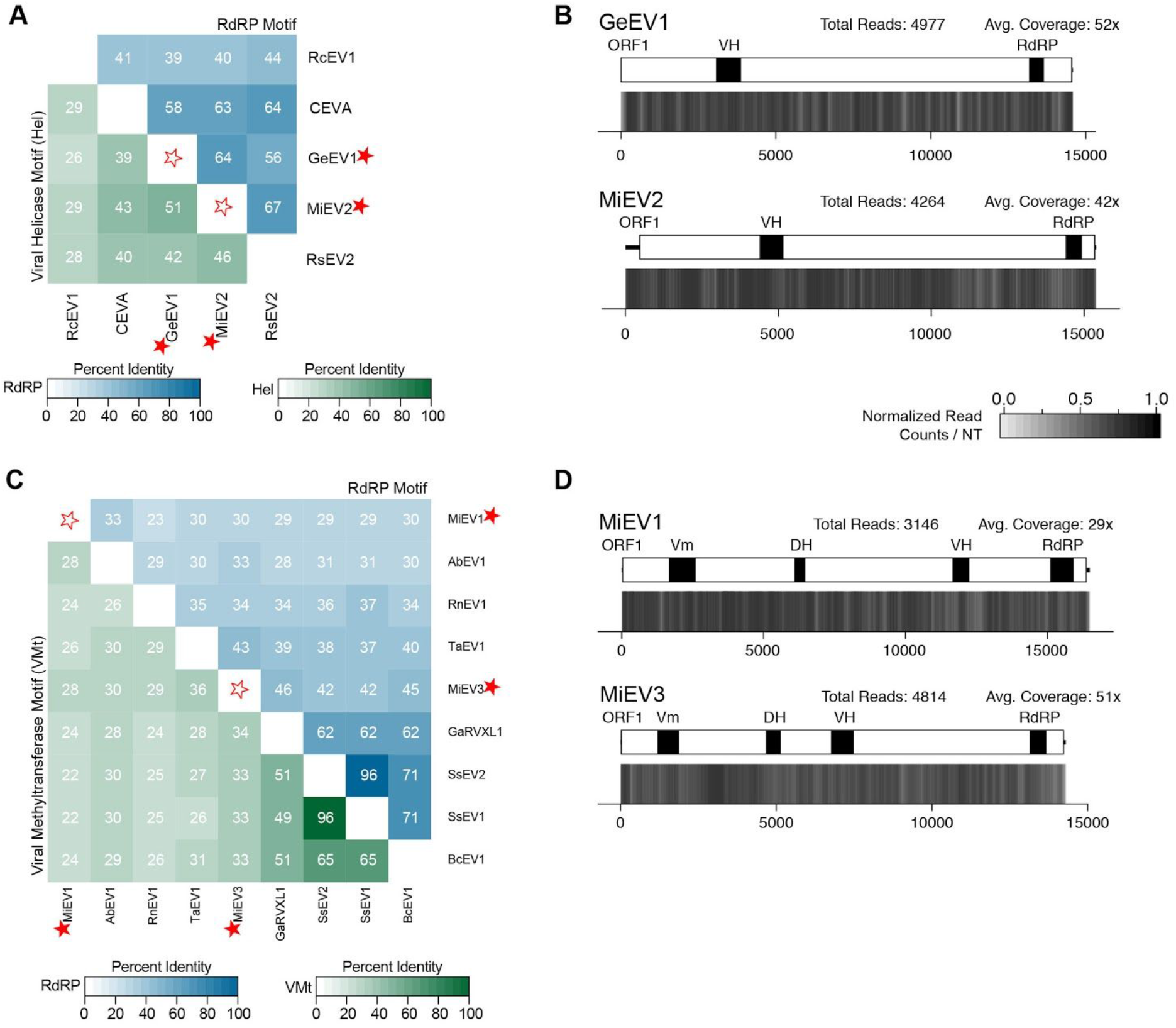
Analyses of identified viral genomes belonging to the family *Endomaviridae.* (A and C) Percent identity matrix, generated by Clustal-Omega 2.1, of mycoviruses in the genus *Alphaendornavirus* (A) and *Betaendornavirus* (C). The top half of both matrices, in the blue scale, is the percent identity of the RdRP motif sequences; the bottom half of the matrix, in the green scale, is the percent identity of the Viral Helicase 1 motif (A) and Viral methyltransferase motif (C). For sake of clarity, the 100% identity along the diagonal has been removed and outlined stars are used to note junction point for the row and column of the viruses identified in this study. (B and D) Density plot of read counts per nucleotide across the viral genome for the alphaendornaviruses, GeEV1 and MiEV2 (B) and betaendornaviruses MiEV1 and MiEV3 (D). Graphical depiction of the genome structure, predicted ORF, and encoded motifs are on top and a heatmap of normalized read counts is on the bottom. Read counts are normalized to a 0 – 1 scale, where the maximum read count for a given genome equals 1. Total reads mapped to the virus genome and average read coverage are noted. Viruses identified in this study are noted with a red star. RdRP: RNA-dependent RNA Polymerase; VH: Viral Helicase 1; Vm: Viral methyltransferase; DH: DEXDx Helicase.

#### Betaendorna virus

The two other endornaviruses identified were MiEV1 and MiEV3, which group with the betaendornaviruses. MiEV3, in particular, forms a strong phylogenetic relationship with other characterized betaendornaviruses, while MiEV1 is the outermost member of this clade (Fig 6A). Unexpectedly, the genome sizes of both viruses are more than 25% larger than other betaendornaviruses; MiEV1 has a genome size of 16.5 kb while MiEV3 has a genome size of 14.3 kb (Table 1). However, the motifs within the protein sequences are as expected, as both MiEV1 and MiEV3 contain a viral methyltransferase domain, helicase domain(s), and an RdRP domain (S7D Fig). Specifically MiEV1 and MiEV3 both encode a DEADx helicase domain and a viral helicase 1 domain, similar to the type species *Sclerotinia sclerotiorum endornavirus-1* (SsEV1) (NC_021706.1). Alignment of the RNA-seq reads to the viral genomes reveals an average coverage per nucleotide of 29x and 51x for MiEV1 and MiEV3 respectively (S7D Fig), and coverage appears uniform across both genomes.

Similar to the alphaendornaviruses, nucleotide identity is generally below 50%, with the exception of SsEV1 and *Sclerotinia sclerotiorum endornavirus 2* strain DIL-1 (SsEV2) which share 82% nucleotide identity. Further analysis of the protein sequences from the RdRP motifs and viral methyltransferase motifs determined that a percent identity threshold for either motif of 75% would be appropriate for species demarcation (S7C Fig). In particular, both MiEV1 and MiEV3 are well below both a nucleotide-based or a protein-motif based threshold. Thus, despite the same fungal host, the uniqueness of the viral genomes sequence for both viruses suggests that both MiEV1 and MiEV3 are new species of the family *Endornaviridae*, and belong within the genus *Betaendornavirus.*

## Positive(+)ssRNA Viruses

### Tobamo-like virus: ArTIV1

A virus sequence was identified in an *Acidomyces richmondensis* sample (PRJNA250470; unpublished) with a single segment genome sequence, 10,291 nt in length, that encoded four predicted ORFs (Table 1). The gene product from ORF2 contains the RdRP domain (Fig 7B) and a BLAST-P search of the ‘nr’ database with this sequence identified two related, full length mycoviruses, *Macrophomina phaseolina tobamo-like virus 1* (MpTIV1) [5] and *Podosphaera prunicola tobamo-like virus 1* (PpTIV1) [56]. As such, this newly identified virus has been named *Acidomyces richmondensis tobamo-like virus 1* (ArTIV1).

For all three mycoviral sequences the first and largest ORF contains a viral methyltransferase and viral helicase domain and ORF2 encodes the RdRP domain (Fig 7B). ORF3 and ORF4 in MpTIV1 have been putatively identified as encoding a movement protein and coat protein respectively [5].

A phylogenetic analysis of the RdRP motif from the three mycoviruses compared to select members of the families *Virgaviridae, Bromoviridae* and genus *Idaeovirus* demonstrated that ArTIV1 is more closely related to the two previously identified mycoviruses than known plant viruses (Fig 7A). Further, the group of mycoviruses form a sister clade to the *Virgaviridae* viruses. Of the viruses currently characterized from this family, only members of the genus *Tobamovirus* have a non-segmented genome similar to ArTIV1 and relatives; other members have multipartite genomes [57].

To further characterize this emerging group of mycoviruses, a percent identity analysis was undertaken, comparing the amino acid sequences from the RdRP and viral helicase motifs to members of the *Virgaviridae* family. The RdRP motif sequences among the mycoviruses share 56 to 63% identity with one another, but less than 44% identity with *Virgaviridae* viruses (Fig 7C). Further, sequences from the viral helicase domain were 59 to 62% identical among mycoviruses, and less than 40% shared identity outside of this clade. Due to these values, it appears that all three viruses are unique viral sequences.

Finally, the RNA-seq data was aligned to the ArTIV1 genome sequence, which revealed uniform coverage along the length of the genome, averaging 1,167 reads per nucleotide (Fig 7B).

To date all viruses of the *Virgaviridae* family infect a plant host, although many members are transmitted by nematodes or plasmodiophorids, a group of parasitic microscopic organisms unrelated to fungi or oomycetes [57]. Tobacco mosaic virus (TMV), the best characterized *Tobamovirus*, can infect, replicate, and persist in the ascomycota fungi *Colletotrichum acutatum* [58]. Further, a virus from the *Bromoviridae* family, Cucumber mosaic virus (CMV) was shown to naturally infect the phytopathogenic fungi *Rhizoctonia solani* and as well as replicate within *Valsa mali* but not *C. parasitica*, and *F. graminearum* [59]. Due to the close contact between phytopathogenic fungi and plants a change in host range of a tobamo-like virus is plausible. Interestingly, the *A. richomondensis* strain analyzed in this study was isolated from acid mine drainage biofilms [60] and is not known to be phytopathogenic. Acid mine drainage biofilms are complex communities comprised of different fungi along with bacteria, archaea and bacteriophages [61, 62]. The role of the mycovirus identified in this study in such an extreme and heterogeneous environment remains to be determined.

### Ambiguiviridae

Four viral genome sequences were identified that share similarity to unclassified (+)ssRNA mycoviruses: *Trichoderma harzianum ssRNA virus 1* (ThAV1) (PRJNA216008; [63]), *Setosphaeria turcica ssRNA virus 1* (StAV1) (PRJNA250530; unpublished), *Verticillium longisporum ssRNA virus 1* (VIAV1) (PRJNA308558; unpublished), and *Periconia macrospinosa ssRNA virus 1* (PmAV1) (PRJNA262386; [64]). To our knowledge, this is the first viral sequence described for the later three fungal species, and the first ssRNA virus isolated from *T. harzianum.*

Six other viruses appear related to these four newly identified viruses: *Diaporthe ambigua RNA virus 1* (DaRV1), *Magnaporthe oryzae RNA virus* (MoRV), *Soybean leaf-associated ssRNA virus* 1 and 2 (SlaRV1, SlaRV2), *Verticillium dahliae RNA virus* (VdRV), and *Sclerotinia sclerotiorum umbra-like virus 1* (SsUIV1) [5,6,65–67]. The RdRP protein sequence of this group of mycoviruses shares some sequence similarity to RdRPs of recently described (+)ssRNA viruses from insects, as well as members of the plant-infecting family *Tombusviridae.* Phylogenetic analysis demonstrates the fungal viruses form a well-supported clade that is separate from both the insect and plant viruses (Fig 8A). A pairwise identity analysis of the RdRP motif region from the viruses analyzed in the phylogenetic tree reveals a range of identities between the fungal viruses, from 36% to 63%, while only sharing 25% to 33% identity with the insect viruses and 26% to 32% identity with the members of the *Tombusviridae* (Fig 8D). A similar analysis using the ORF1 protein coding sequence, from only the mycoviral genomes, demonstrates 18 to 38% identity among sequences (Fig 8D).

**Fig 8.**
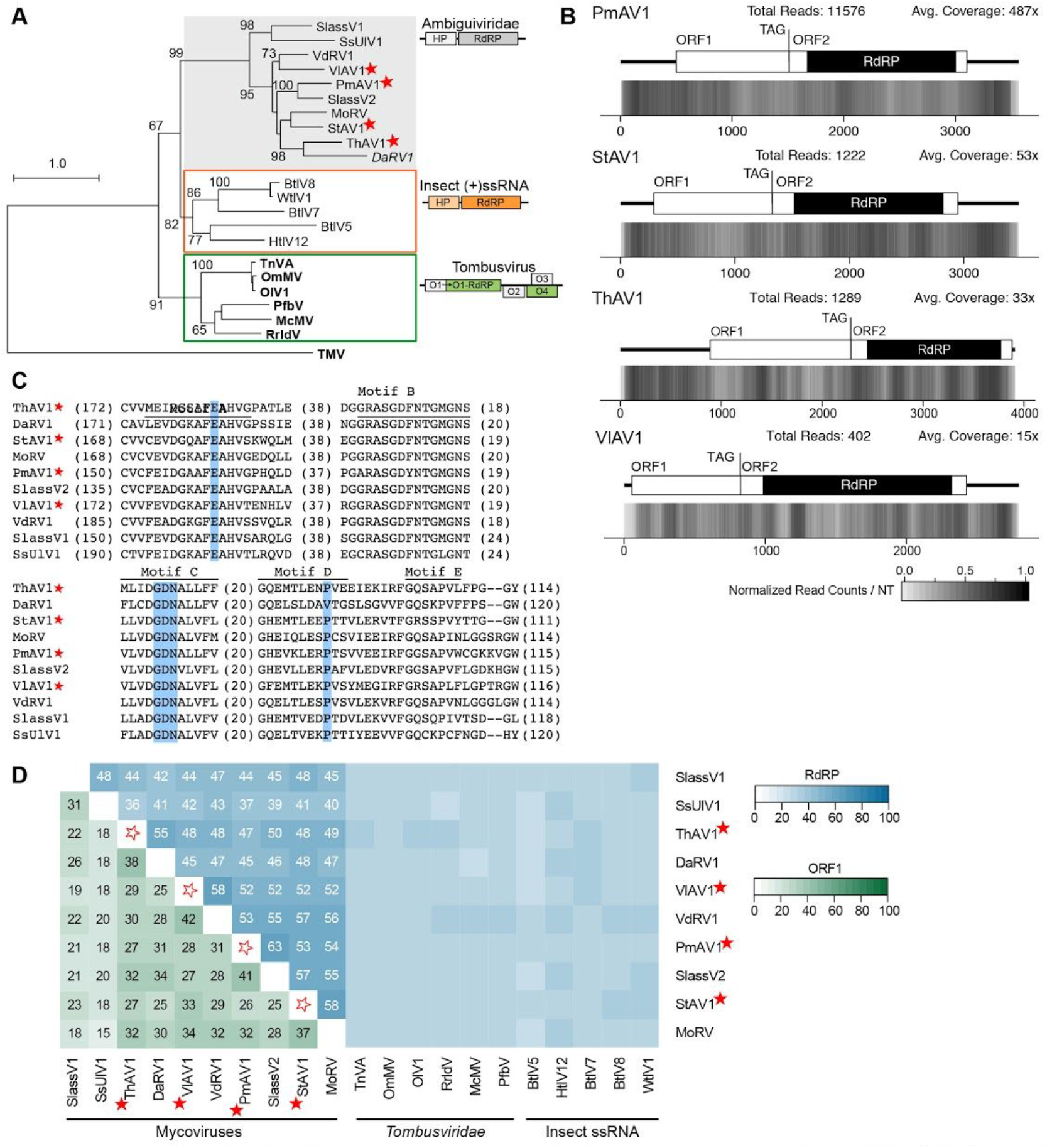
Analyses of four newly identified (+)ssRNA viral genomes and viruses belonging to the proposed family *Ambiguiviridae.* (A) Phylogram of the RdRP-motif for four new viruses and related ambiguiviruses along with select members of the family *Tombusviridae* and members of an unnamed group of (+)ssRNA insect viruses; sequence from *Tobacco Mosaic Virus* (TMV) serves as the outgroup. Bolded names are ICTV-recognized member species while italicized names are recognized, but unclassified viruses. Full names and accession numbers for protein sequences used are in Table S2. Numbers at the nodes indicate bootstrap support values ≥60% (1000 replicates). Scale bar on the phylogenetic tree represents 1.0 amino acid substitutions per site. Diagrammatic genome structures and predicted ORFs for each clade of viruses is to the right of the phylogram. (B) Density plot of read counts per nucleotide across four viral genomes. Graphical depiction of the genome structure, predicted ORFs, encoded motifs and the amber stop codon (TAG) for each genome are along the top and a heatmap of normalized read counts is on the bottom. Reads are normalized to a 0 −1 scale, where the maximum read count for a given genome equals 1. Total reads mapped to the virus genome and average read coverage are noted. (C) Multiple sequence alignment of the RdRP motifs of members of the Ambiguiviridae family. Motifs A-E found in (+)ssRNA viral RdRP sequences are indicated along the top of the alignment. Conserved residues that differ from canonical (+)ssRNA RdRP sequences, including the unusual GDN triad found in Motif C, are highlighted in blue. Numbers in parentheses are residues not shown. (D) Percent identity matrix, generated by Clustal-Omega 2.1, of mycoviruses that comprise the proposed family *Ambiguiviridae.* The top half of the matrix, in the blue scale, is the percent identity of the RdRP motif sequence and the bottom half of the matrix, in the green scale, is the percent identity of the ORF1 amino acid sequence. For sake of clarity, the 100% identity along the diagonal has been removed and outlined stars are used to note junction point for the row and column of the viruses identified. All RdRP motif sequences share ≥35% identity and ORF1 protein sequences share ≥15% identity. Viruses identified in this study are noted with a red star. RdRP: RNA-dependent RNA Polymerase; ORF: Open Reading Frame; HP: Hypothetical Protein.

Together these ten fungal viruses are characterized by a single segment genome, 2.6 kb to 4.5 kb in length, that encodes two ORFs, where the second ORF contains the RdRP domain (Table 1). The amber stop codon ‘UAG’, found at the end of the first ORF, has been suggested to result in readthrough to produce a fusion protein with the downstream RdRP [67]. All members of this group of viruses contain the amber stop codon, and both ORF1 and ORF2 are found in the same frame, a requirement for readthrough. Additionally, all 10 viruses share a number of key differences in residues commonly conserved within (+)ssRNA viruses. For instance, the second aspartic acid residue in motif A, a feature of (+)ssRNA viruses [68], is instead a glutamic acid (Fig 8C). Additionally, these 10 viruses have a GDN triad in motif C versus the canonical GD<u>D</u> found in (+)ssRNA viruses (Fig 8C). This triad has previously been observed within some (-)ssRNA viruses [68], although within (+)ssRNA viruses, modification of the GDD to GDN has repeatedly demonstrated a detrimental effect on enzymatic activity [69–71]. Finally, within motif D, a positively-charged lysine, common to nearly all RdRP sequences [68, 72], is a nonpolar proline or valine residue (Fig 8C).

The 5’UTRs range in size from 283 bp to 889 bp in length for all the mycoviruses in this group, with the exception of VdRV1 and VIAV1, the two viruses isolated from species of *Verticillium*; these 5’UTRs are 11 nt and 54 nt respectively. Similarly, the 3’UTR sequences range in length from 372 bp to 1279 bp, while those of VdRV1 and VIAV1 are 182 nt and 29 nt respectively (Fig 8C). Average coverage per nucleotide across the viral genomes of the four identified viruses is fairly uniform across the genome with ranges from 15x for VIAV1 to 734x for PmAV1 (Fig 8B).

The characteristics of these four identified viruses and their closest mycoviral relatives supports the creation of a new family of viruses within the (+)ssRNA class. We propose the Mycoviruses *lomDusviriaae* insect sskna establishment of the family *Ambiguiviridae* that contains DaRV1 and other recently discovered viruses related to it, including the four described in this study. The proposed name references the host of the first virus identified (*D. ambigua*), while also highlighting the unusual nature of the GDN triad within the RdRP motif.

### Yadokariviridae

Samples *Aspergillus homomorphus* (PRJNA250984; unpublished) and *Penicillium digitatum* (PRJNA352307; unpublished), which yielded a totivirus (AhoTV1) and partitivirus (PdPV1) respectively, each also contained a second, unrelated contig containing an RdRP motif. BLAST-X analysis of these nucleotide sequences determined they were both related to a group of unclassified, (+)ssRNA viruses, that includes four other mycoviruses. As described below, these two viruses are new members of a recently proposed family, *Yadokariviridae* [73].

*Aspergillus homomorphus yadokarivirus 1* (AhoYV1) and *Penicillium digitatum yadokarivirus 1* (PdYV1), along with other members of this group, are characterized by having single segment genomes that encode a single predicted ORF, the RdRP (Table 1). Genome sizes range from 3,144 nt (PdYV1) to 5,089 (*Yado-kari virus 1*, YkV1). The length of the RdRP protein sequence is predicted to range from 962 aa for *Aspergillus foetidus slow virus 2* (AfSV2) [74] to 1,430 aa for YkV1 [75]. The lengths of the 5’UTRs are all greater than 500 nt, except for PdYV1. Attempts to extend the sequence of the 5’UTR of PdYV1 using the RNA-seq data were not successful; however, as the average coverage per nucleotide for PdYV1 was low, just 23x (Fig 9C), additional experiments with the fungal source material may be fruitful. Lengths of the 3’UTRs for all six viruses are also largely similar, between 200 to 400 nt, except for YkV1 which has a 3’UTR of 1,223 nt. A genome segment belonging to a *Yadokarivirus* that encodes a coat protein has not been identified. Alignment of the RNA-seq reads to the genomes of AhoYV1 and PdYV1 reveals an average coverage per nucleotide of 151x and 23x respectively (Fig 9B-C).

**Fig 9.**
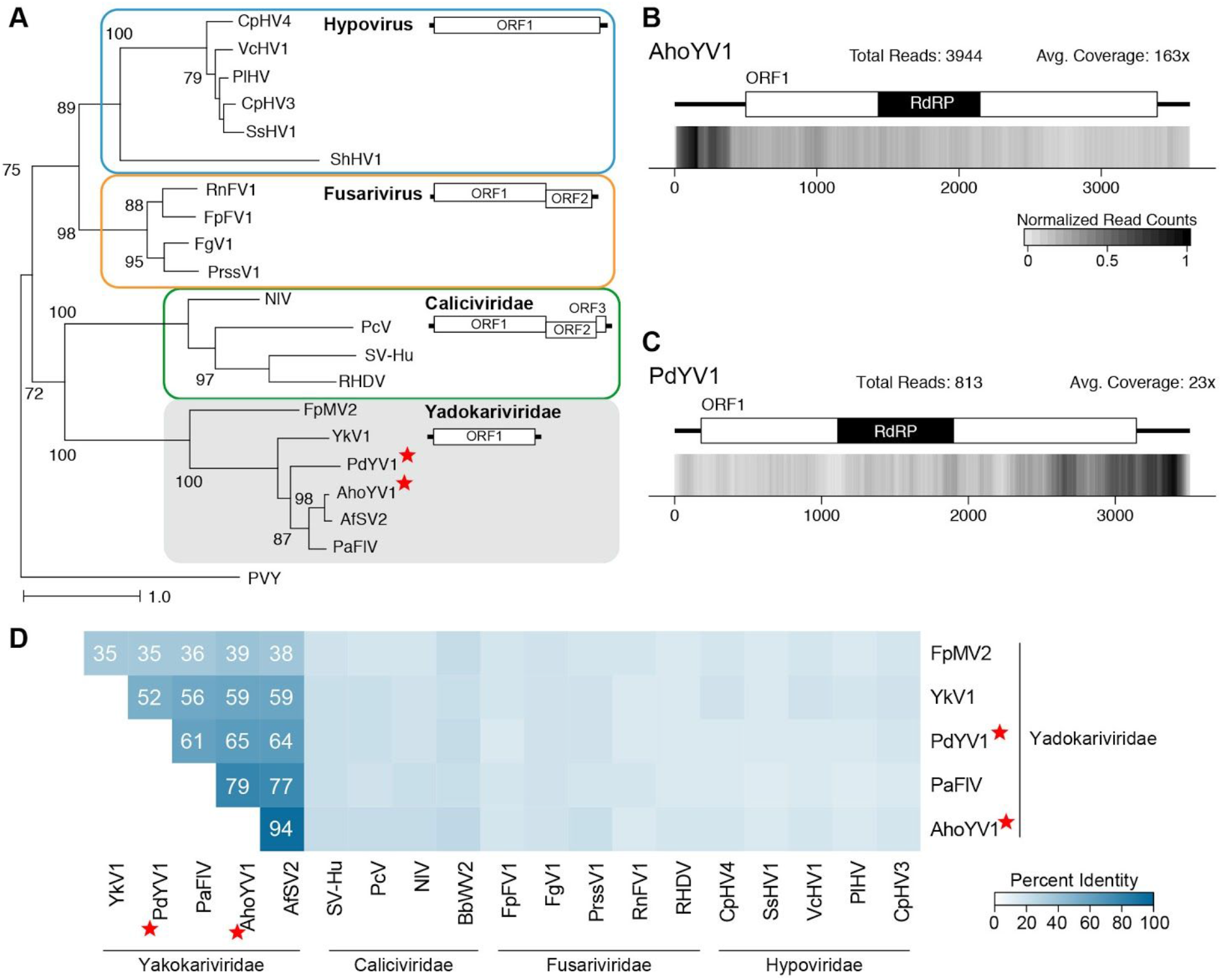
Analyses of *Aspergillus homomorphus yadokarivirus 1* (AhoYV1) and *Penicillium digitatum yadokarivirus 1* (PdYV1) viral genome sequences, members of the proposed family *Yadokariviridae.* (A) A phylogram of the RdRP coding sequence from select (+)ssRNA viruses from the *Yadokariviridae* family, in addition to members of the families *Caliciviridae, Fusariviridae*, and *Hypovirídae*; sequence from *Potato Virus Y* (PVY) serves as the outgroup. Virus names have been abbreviated; full names and accession numbers for protein sequences used in the alignment are in Table S2. Phylogenetic tree was generated with RAxML; scale bar represents 1.0 amino acid substitutions per site and numbers at the nodes indicate bootstrap support over 70% (1000 replicates). To the right of the phylogram are representative diagrams of the genome structures for each family of viruses. (B and C) Density plot of read counts per nucleotide across the viral genome for (B) AhoYV1 and (C) PdYV1. Graphical depiction of the genome structure, the predicted single ORF and the RdRP motif are along the top and a heatmap of normalized read counts is on the bottom. Read counts are normalized to a 0 −1 scale, where the maximum read count for a given genome equals 1. Legend for (C) is the same as the legend found in (B). Total reads mapped to the virus genome and average read coverage are noted for each virus. (D) Percent identity matrix generated by Clustal-Omega 2.1 using the RdRP motif sequences from the viruses analyzed in (A). For the sake of clarity, only the top half of the matrix is included and the 100% values along the diagonal have been removed. Percent identities greater than 25% are indicated. Newly identified viruses, AhoYV1 and PdYV1, are noted with a red star. RdRP: RNA-dependent RNA Polymerase; ORF: Open Reading Frame.

A phylogenetic analysis of the RdRP protein sequence reveals a well supported clade that is most closely related to the *Caliciviridae* and separate from other known mycoviral families including *Hypoviridae* and fusariviruses (Fig 9A). AhoYV1 in particular is most closely related to AfSV2; both viruses were isolated from a member of the genus *Aspergillus* [74].

As percent identity between virus RdRP sequences is a common criteria for defining new groups of viruses, the RdRP motif sequences from the viruses in Fig 9A were analyzed to identify a potential threshold value. Indeed a threshold between 26 to 35% identity easily classifies these viruses together, but separate from other known (+)ssRNA viruses (Fig 9D). An additional threshold of 95% would to distinguish each sequence evaluated here as a unique viral species; however, as the pairwise percent identity between AhoYV1 and AfSV2 of 94% is somewhat higher then is generally observed for unique species, a lower threshold may be more appropriate. Identification of additional sequences belonging to this group of viruses will further refine these sequence-based characteristics.

Another feature common to these viruses is the presence of at least one additional virus within the fungal host. Co-incident infection is not an uncommon occurrence in fungi; however, it has been demonstrated for two yadokariviruses, AfSV2 and YkV1, that isolated virus particles contain a second virus, related to Totiviruses [74, 75]. Further, Zhang and colleagues demonstrated that YkV1 is *trans-e*ncapsidated with the capsid protein from the other virus, YnV1 [75]. As caliciviruses and totiviruses share a distant but strongly supported ancestry following diversification of the picorna-like superfamily [76], and both AfSV2 and YkV1 are present within virions encapsidated by totivirus capsid proteins, it is intriguing to speculate about the mechanisms at play, and the requirements for such an interaction. A totivirus was also identified in the *A. homomoφhus* sample (AhoTV1, discussed above), however only a partitivirus (PdPV1-HSF6) was identified along with PdYV1. For the remaining two members of this group of viruses, it is not possible to know the identity of the putative coat-protein donor, as PaFIV1 was found to co-infect *P. aurantiogriseum* along with a fusarivirus, totivirus, unclassified virus, and partitivirus [4] and FpMyV2 was identified in a fungal strain that contained 13 other viruses from a variety of families [38].

Further study of the fungi that host a member of this group of viruses will answer questions regarding the relationship between the capsid-less virus and the capsid-donor, and may provide an interesting system for studying the evolution of a capsid-less virus. Due to the distinct phylogenetic relationship between these six identified and known (+)ssRNA viruses, along with the unique lifestyle, we support the recommendations to create a new virus family, named *Yadokariviridae* after YkV1, where *yadokari*, is the Japanese word for hermit crab [73, 77].

### Narnaviridae and Ourmia-like virus

Members of the family *Narnaviridae* have single molecule, positive-strand RNA genomes ranging in length from 2.3 to 2.9 kb and encode an RdRP as the only polypeptide [78]. There are currently two genera within this family: *Narnavirus* and *Mitovirus*, where mitoviruses are characterized as viruses that replicate within the mitochondria of filamentous fungal hosts [78]. Indeed, a non-interrupted ORF is only predicted in these viruses when using the mitochondrial codon usage where tryptophan is encoded by UGA [79]. Here we describe five new viruses with similarity to members of the genus *Mitovirus* from RNA-seq datasets from four fungal hosts: *Setosphaeria turcica mitovirus 1* (StMV1) (PRJNA250530; unpublished), *Colletotrichum falcatum mitovirus 1* (CfMV1) (PRJNA272832; [80]), *Loramyces juncicola mitovirus 1* (LjMV1) (PRJNA372853; unpublished), and *Ophiocordyceps sinensis mitovirus* 1 and 2 (OsMV1, OsMV2) (PRJNA292632; [81]).

The five mitoviruses have genome sizes within the expected range (Table 1), and a single ORF is present only when using the mitochondrial codon table. The percent of A+U in the coding strand for these five viruses ranges from 53% to 70%; characteristically, mitoviruses have 62 to 73% A+U [78], although the percent observed in recently identified viruses is as low as 53% [5]. The predicted protein within each genome contains a Mitovir_RNA_pol domain, with a length ranging from 652 to 716 aa. Alignment of the RNA-seq reads to the viral genomes demonstrates end to end coverage for each virus, with average coverage per nucleotide ranging from 51x for OsMV1 to 540x for LjMV1 (S8D Fig).

**S8 Fig.**
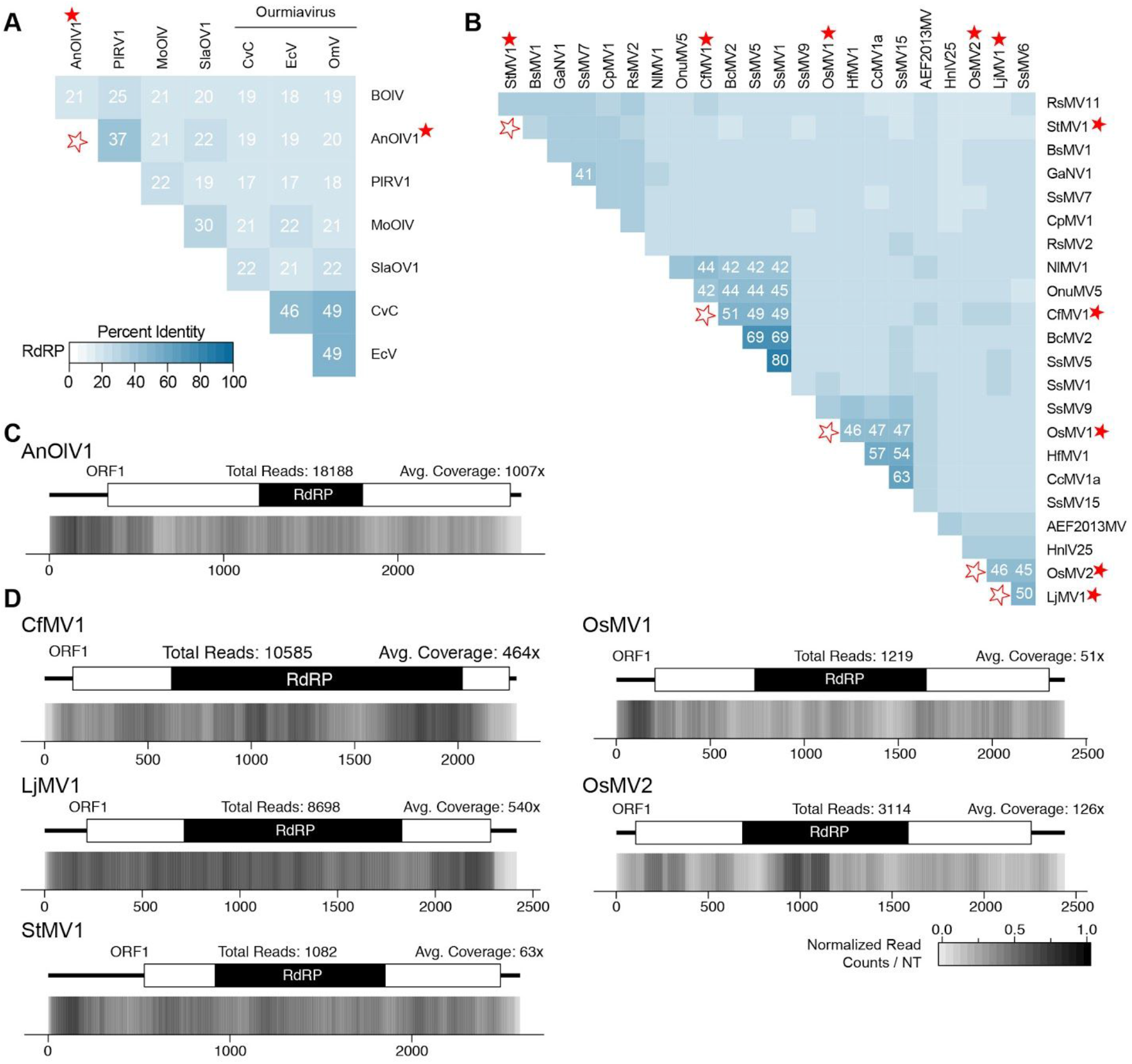
Analyses of identified viruses related to *Ourmiavirus* and *Narnaviridae.* (A and B) Percent identity matrix of the RdRP protein coding sequence, generated by Clustal-Omega 2.1, of ourmia-like mycoviruses (A) and the genus *Mitovirus* from the family *Narnaviridae* (B). For the sake of clarity, only the top half of the matrix is shown and the 100^?^/o identity along the diagonal has been removed; outlined stars are used to note junction point for the row and column of the virus(es) identified. The heatmap scale in (B) is the same as found in (A). Sequences used in analysis can be found in Table S2. (C and D) Density plot of read counts per nucleotide across the viral genomes of the ourmia-like viruses (C) and mitoviruses (D) identified in this study. Graphical depiction of the genome structure, ORF, and predicted RdRP motif for each genome are along the top and a heatmap of normalized read counts is on the bottom. Read counts are normalized to a 0 −1 scale, where the maximum read count for a given genome equals 1. Total reads mapped to the virus genome and average read coverage are noted. Viruses identified in this study are noted with a red star. RdRP; RNA-dependent RNA Polymerase.

A phylogenetic analysis of the RdRP coding sequence demonstrates strong support for all five newly identified viruses among members of the genus *Mitovirus* (Fig 7B). Criteria for species demarcation within the genus *Mitovirus* is not yet full defined, however, viruses with greater than 90% identity within the RdRP protein sequences are considered the same species [20]. Here we observe that StMV1, CfMV1, OsMV1, OsMV2, and LjMV1 have less than 52% identity to any currently available mitovirus sequence (S8B Fig). Taken together, the above characteristics demonstrate that these five viruses are new species within the genus *Mitovirus.*

Ourmiaviruses are also positive-strand RNA viruses, composed of three genome segments that encode an RdRP, movement protein, and coat protein, isolated from plant hosts [19, 82]. Despite usage of standard eukaryotic codons, phylogenetically the RdRP protein sequence is most closely related to that of *Narnavirus* [19]. Recently viruses have been identified from fungal hosts with sequence similarity to that of the plant-hosted ourmiaviruses. One such virus was identified in this study: *Aspergillus neoniger ourmia-like virus 1* (AnOIV1) (PRJNA250996; unpublished). The single segment, 2.7 kb genome encodes a single polyprotein that contains an RdRP motif (Table 1). Alignment of the RNA-seq data to the genome revealed an average coverage of 1007x (S8C Fig).

A phylogenetic analysis of this virus, along with related ourmia-like viruses from other fungi, plant-hosted ourmiaviruses, and mitoviruses illustrates a well supported clade containing AnOIV1 along with *Phomopsis longicolla RNA virus 1* (PIRV1) [83] and *Botrytis ourmia-like virus* HAZ2-3 (BolV) [84] (Fig 7B). A percent identity analysis of the fungal and plant viruses found in Fig 7B demonstrates a percent identity of approximately 50% among the recognized *Ourmiaviruses*, while the fungal-hosted viruses shared from 17 to 37% identity (S8A Fig), with AnOIV1 and PIRV1 the most similar. As there is limited sequence similarity, and AnOIV1 was identified from a new fungal host, it would appear that this is a new species within this yet-to-be fully characterized viral genus. Identification of additional *Ourmiaviruses* and related mycoviruses will shed further light on the specific relationship between these plant and fungal viruses.

### Hypoviridae

The *Hypoviridae* family consists of positive sense, single-stranded RNA viruses that are approximately 9.1 to 12.7 kb in length, containing either one long ORF or two ORFs [85]. Further, these viruses are capsidless and use a host-derived, trans-Golgi network for genome replication [86, 87]. Infection by Hypoviruses can often result in reduced fungal virulence, a hallmark symptom of early members of the *Hypoviridae* family.

Hypoviruses identified from four transcriptomic datasets are described here: *Sclerotinia homoeocarpa hypovirus 1* (ShHV1) (; PRJNA167556[88]), *Trichoderma asperellum hypovirus 1* (TaHV1) (PRJNA261111; [89]), *Setosphaeria turcica hypovirus 1* (StHV1) (PRJNA250530; unpublished), and *Fusarium graminearum hypovirus 1* (FgHV1) (PRJNA263651; [90]). The virus FgHV1 was expected, as the sequencing experiment featured *F. graminearum* strain HN10 infected with FgHV1 [90]; to our knowledge, this is the first mycovirus identified from S. *turcica* and *T. asperellum*, and the first hypovirus from S. *homoeocarpa.*

There are currently four species officially recognized in the *Hypoviridae* family: *Cryphonectria hypovirus* 1, 2, 3, and 4 (CHV1, CHV2, CHV3, and CHV4), although several other related viruses have recently been identified from eight other fungal species. Of the currently recognized hypoviruses two clades form that appear to be distantly related, whereby CHV1 and CHV2 are most closely related and CHV3 and CHV4 are together in a separate grouping. To this end, two new genera have been proposed, *Alphahypovirus* [91] and *Betahypovirus* [92], where CHV1 and CHV2 are classified in the former and CHV3 and CHV4 in the later [85]. A phylogenetic analysis using the RdRP protein sequence of the four viruses identified here, along with known and related hypoviruses, revealed that FgHV1, ShHFV1, and TaHV1 groups with the *Alphahypoviruses* while StHV1 was found with the *Betahypoviruses* (Fig 7C). A further analysis of these viral sequences, described below within the context of the two recently proposed genera, determined that ShHV1, StHV1, and TaHV1 are new species within the *Hypovirus* genus.

#### Aiphahypovirus

Initially, members of *Alphahypovirus* were characterized as having two ORFs, where the second ORF contained protease, RdRP, and Helicase domains. This included CHV1 [93], CHV2 [94], FgHV1 [91], and *Rosellinia necatrix hypovirus 2* (RnHV2) [95]. However, identification of additional viruses has revealed members with a single, large ORF, including *Fusarium langsethiae hypovirus 1* (FIHV1) [96], *Fusarium graminearum hypovirus 2* (FgHV2) [91]. *Fusarium poae hypovirus 1* (FpHV1) [38], and *Rosellinia necatrix hypovirus 1* (RnHV1) [95]. Independent of the number of ORFs, alphahypoviruses have large genomes, with lengths ranging from 12.5 to 14.4 kb. Similarly, newly identified viruses ShHV1 and TaHV1 have genome lengths of 12.4 and 14.2 kb respectively, and single-ORF genome structures (S9A Fig, Table 1).

For a new species to be classified, criteria generally include host species and percent identity with other identified species. Official criteria do not yet exist for the family *Hypoviridae*, and separate criteria for the two proposed genera may be appropriate given the lack of sequence homology between the two groups. Previously it has been observed that CHV1 and CHV2, while both isolated from *Cryphonectria parasitica*, demonstrate only ~67% percent identity within the RdRP amino acid sequence. Expanding this analysis to all viruses within the genus *Alphahypovirus* reveals a maximal identity of 90%, between FgHV2 and FpHV1 (S9B Fig); this excludes the 99% identity observed between the two sequences of FgHV1 which are known to be the same species of virus [90]. Interestingly, there is a cluster of similar hypoviruses from three different *Fusarium* species, FIHV1, FpHV1, and FgHV2, which excludes the fourth *Fusarium* hypovirus, FgHV1. An appropriate threshold for evaluating new species of *Alphahypovirus* remains to be determined; however as ShHV1 and TaHV1 share only 50% and 31% identity with RnHV2 and CHV1 respectively, and were identified from unique fungal hosts, these viruses appear to be new species within the family *Hypovirus.*

**S9 Fig.**
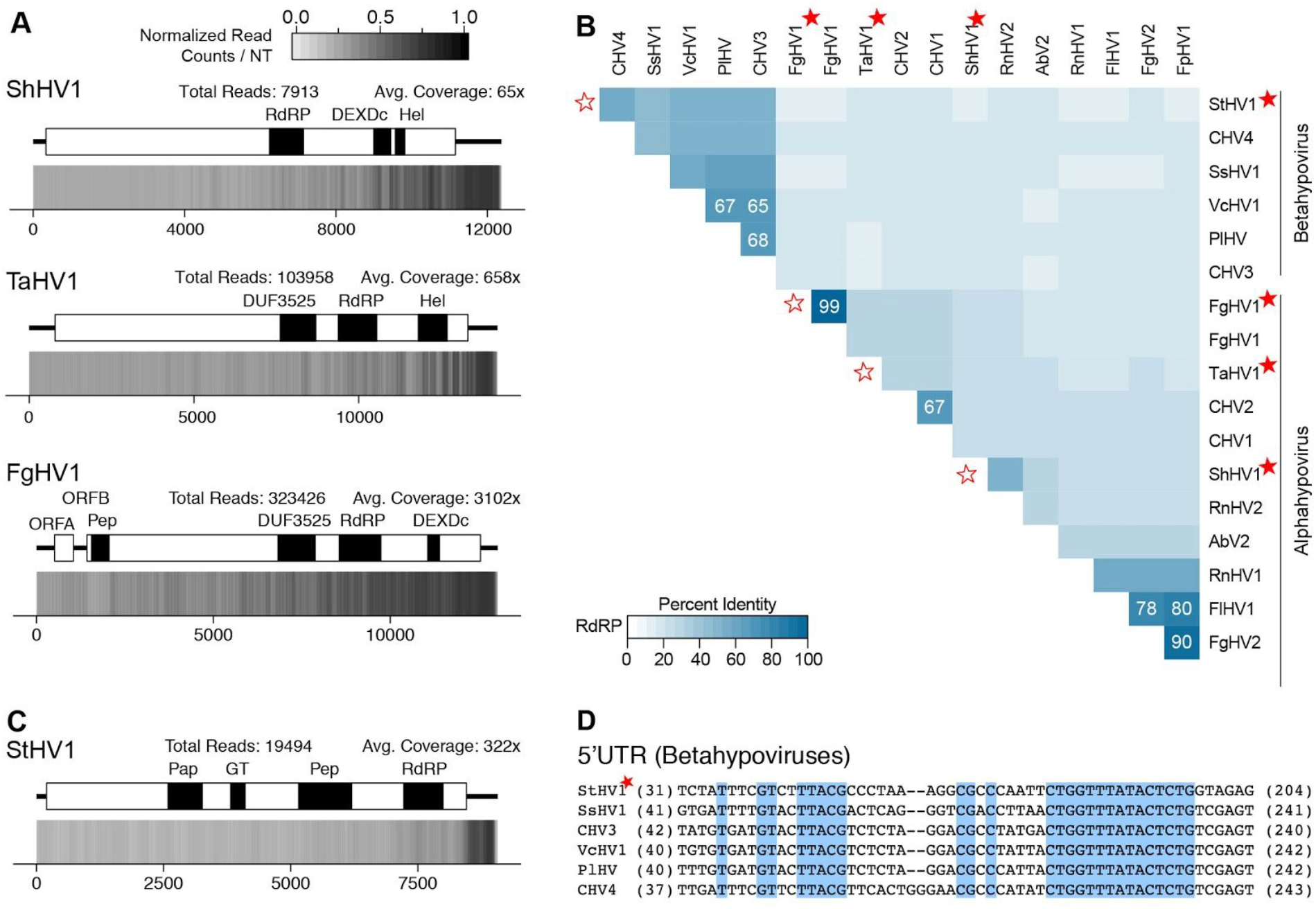
Analyses of identified viral genomes belonging to the family *Hypoviridae.* (A and C) Density plot of read counts per nucleotide across the viral genomes that group with betahypoviruses (A) and alphahypoviruses (C). Graphical depiction of the genome structure, predicted ORFs, and encoded motifs are along the top and a heatmap of normalized read counts is on the bottom. Read counts are normalized to a 0 – 1 scale, where the maximum read count for a given genome equals 1. Total reads mapped to the virus genome and average read coverage are noted. (B) Percent identity matrix, generated by Clustal-Omega 2.1, of the RdRP-motif containing protein coding sequence from the newly identified viruses and related hypoviruses. For sake of clarity, the 100% identity along the diagonal has been removed and outlined stars are used to note the junction point for the row and column of the viruses identified. Where sequences have similarity ≥60% is indicated. Full names and accession numbers for sequences used are in Table S2. (D) Alignment of 5’UTR sequences from alphahypoviruses where conserved nucleotides are highlighted in blue and amino acids not included in the alignment are noted in parentheses. Viruses identified In this study are noted with a red star. Pap: papain-like cysteine protease domain homologs; RdRP: RNA-dependent RNA Polymerase; Hel: Helicase; GT: UDP-glucose/sterol glucosyltransferase; Pep: peptidase; DUF: Domain of Unknown Function.

A final analysis of the three alphahypoviruses identified from this study was the alignment of RNA-seq reads to the viral genomes. Total reads that aligned range from 7,913 for ShHV1 to 323,433 for FgHV1, and in all three instances, the peak coverage is closest to the 3’end of the genome (S9A Fig).

#### Betahypovirus

Viruses in the *Betahypovirus* genus generally encode one ORF that contains protease, RdRp, Helicase, and UGT domains [92] and tend to have shorter genomes than the *Alphahypoviruses*, at approximately 9 to 11 kb in length. These same characteristics are also observed in StHV1, which has a length of 9163 bp and a polyprotein predicted to encode a papain-like cysteine protease domain along with a UGT, peptidase-C97, RdRP, and Helicase-HA2 domain (S9C Fig, Table 1). Additionally, the 5’end of the 5’UTR sequence has high sequence identity across viral genomes from this genus; a multiple sequence alignment of this region from StHV1 and related *Betahypoviruses* reveals similarly high conservation within the StHV1 genome, including a 15-nt sequence that is present in all viruses analyzed (CTGGTTTATACTCTG) (S9D Fig). Characterization of the StHV1 sequence also included alignment of the RNA-seq reads to the viral genome, resulting in a total of 376,89 reads aligned, and an average coverage per nucleotide of 617x (S9C Fig).

A percent identity analysis of the RdRP amino acid sequences of viruses within the genus *Betahypovirus* reveals a maximal identity of 68%, between PIHV and CHV3 (S9B Fig) while StHV1 shares 46% to 54% identity with other viruses. As such, a threshold of 70% identity within the RdRP amino acid is suggested as a criteria to distinguish between different species of *Betahypoviruses.*

### Fusariviridae

Members of the proposed family *Fusariviridae* have mono-segmented, plus strand, single-stranded RNA genomes that encode two to four ORFs. ORF1 is the largest and contains the RdRP and Helicase motifs, while the subsequent and second largest ORF encodes an unknown protein, often containing an SMC motif. One to two additional small proteins have been predicted at the 3’end of some virus genomes. Here we have identified seven new viruses that group with fusariviruses.

*Aspergillus ellipticus fusarivirus 1* (AeFV1) (PRJNA250911; unpublished), *Morchella importuna fusarivirus 1* (MiFV1) (PRJNA372858; unpublished), *Neurospora discreta fusarivirus 1* (NdFV1) (PRJNA257829; [97]), and *Rutstroemia firma fusarivirus 1* (RfFV1) (PRJNA372878; unpublished) are to our knowledge, the first viruses to be identified in these species. *Sclerotinia homoeocarpa fusarivirus 1* (ShFV1) (PRJNA167556; [88]) and *Gaeumannomyces tritici fusarivirus 1* (GtFV1) (PRJNA268052; [34]) are the first fusariviruses identified in these fungi, and this is the first full-length description of *Zymoseptoria tritici fusarivirus 1* (ZtFV1) (PRJNA237967; [98]) (PRJEB8798; [99]) (PRJNA253135; [100]) (PRJNA179083; unpublished).

The genome of MiFV1 encodes three predicted ORFs, where the first contains both the RdRP and helicase motifs, the second contains no predicted domains, and the third ORF contains an SMC (structural maintenance of chromosomes) domain (S10C Fig). The genomic structure of the other six viruses identified in this study encode two predicted ORFs, again with the first encoding RdRP and helicase motifs (S10C Fig). ORF2 of ZtFV1 contains a predicted Spc7 domain (found in kinetochore proteins), a domain also found in *Rosellinia necatrix fusarivirus 1* (RnFV1) [101]. Within ORF2 of AeFV1, GtFV1, NdFV1 and RfFV1 is a predicted SMC domain, while no functional domain is predicted within ORF2 of ShFV1. A previous phylogenetic analysis determined that the closest relatives to the viral SMCs are the proteins from the eukaryotic SMC5 and SMC6 subfamily and that because the SMC domain is widely present throughout members of the *Fusariviridae* family, a common ancestral origin is probable [102].

A phylogenetic analysis of the RdRP amino acid sequence of these seven viruses and twelve other published fusariviruses reveals two strongly supported clusters (S10A Fig). “Group 1” contains 14 viruses while “Group 2” contains the remaining five fusariviruses. The genome structures of the two clusters suggests some general patterns; all members of Group 1 have a predicted functional domain within the second largest ORF, either an SMC or an Spc7 domain as described above. Although, FgV1 encodes three ORFs, unlike related viruses that encode just two. Conversely, viruses within Group 2 generally do not have a functional domain predicted within ORF2, the exception is MiFV1 which contains an SMC domain. Further, three ORFs are predicted for four of the five members, with ShFV1 being the exception. Identification of additional viral genomes that belong to this family will help to elucidate the support for these separate clusters and may reveal additional distinguishing characteristics.

**S10 Fig.**
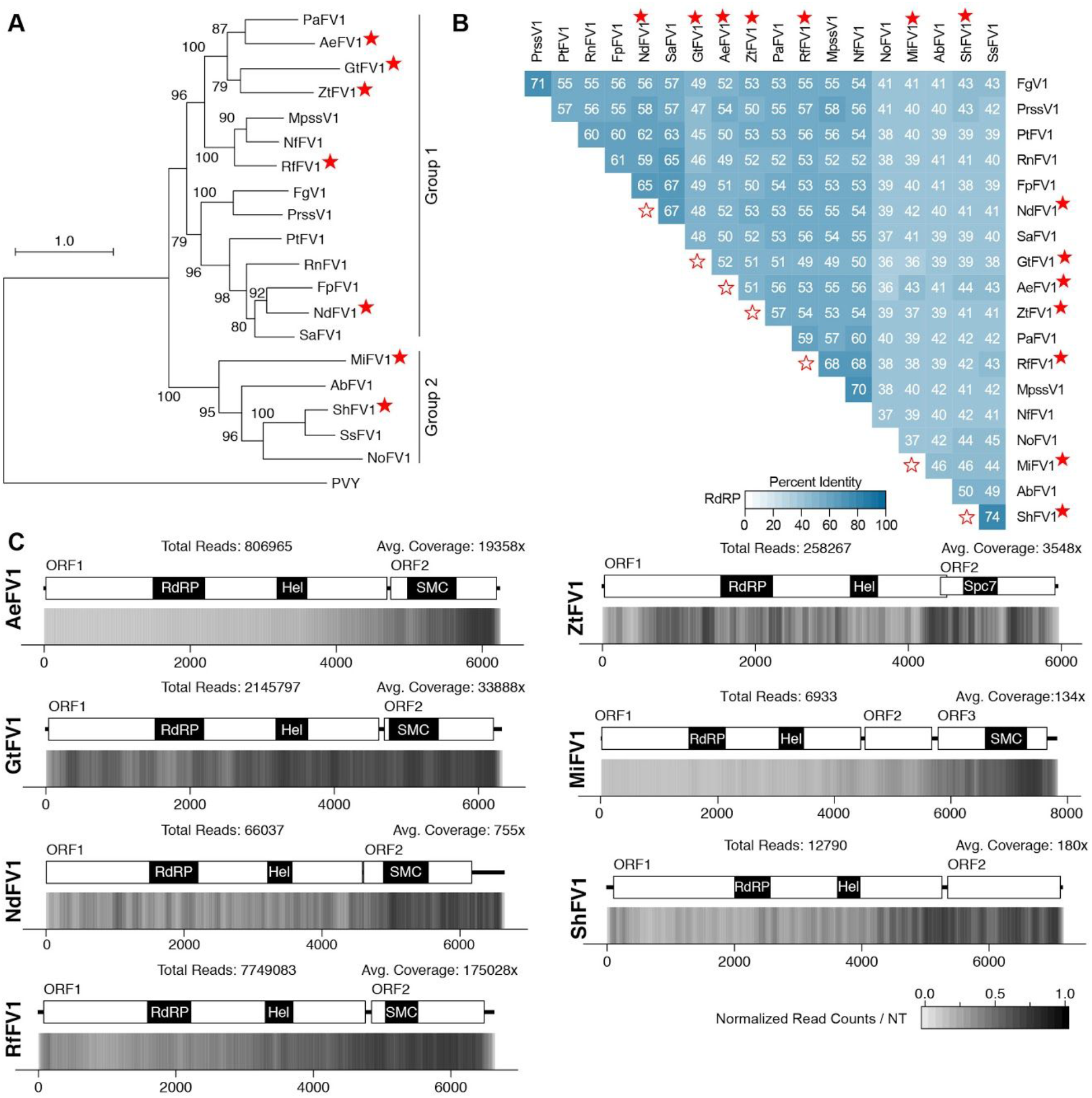
Analyses of viral genomes that belong with known fusariviruses. (A) Phylogram of fusariviruses based on the ORF1 protein coding sequence which contains the RdRP motif. Full names and accession numbers for protein sequences used are in Table S2, including the outgroup Potato Virus Y (PVY). Scale bar represent 1.0 amino acid substitutions per site and numbers at the nodes indicate bootstrap support over 70% (1000 replicates). (B) Percent identity matrix generated from Clustal 2.0 using the RdRP motif sequences from fusariviruses. For sake of clarity, the 100% identity along the diagonal has been removed and open stars note the junction point between the row and column of the viruses identified in this study. (C) Density plot of read counts per nucleotide across the viral genomes identified in this study. Graphical depiction of the genome structure, predicted ORFs and encoded motifs are on top and a heatmap of normalized read counts is on the bottom. Read counts are normalized to a 0 – 1 scale, where the maximum read count for a given genome equals 1. Total reads mapped to the virus genome and average read coverage are noted. RdRP: RNA-dependent RNA Polymerase; Hel: Helicase domain; SMC: SMC N terminal domain; Spc7: Spc7 kinetochore domain.

As *Fusariviridae* is not yet an officially recognized family of viruses, specific characteristics for distinguishing new viral species or genera have yet to be established. In addition to phylogenetic analyses and identity of fungal host species, percent identity across the protein coding sequence of the RdRP-containing ORF is frequently used as a classifier. To date, fusariviruses previously characterized have all been isolated from unique fungal species, including two different species of *Fusarium*, and none share greater than 57% identity with one another (S10B Fig). Viruses identified in this study are also from unique fungal species, including ShFV1 which was isolated from a different species than *Sclerotinia sclerotiorum fusarivirus 1* (SsFV1) [103]. Using the data available to date, two criteria are recommended to qualify viruses as new species: a unique fungal host and a percent identity of the RdRP sequence ≤60% in comparison to existing fungal sequences.

The ZtFV1 sequence described here was identified within the SRA sample PRJNA179083, however interestingly there are three other RNA-seq experiments available for this organism from three other research groups. All four sequencing samples appear to contain the same fusarivirus; indeed a full length sequence was isolated from samples from both the Max Planck Institute for Terrestrial Microbiology (PRJNA237967; [98]) and Rothamsted Research (PRJEB8798; [99]). For the sample from the Institut national de la recherche agronomique (INRA) (PRJNA253135; [100]), a contig covering 76% of the full length was identified, and manual analysis determined sufficient reads to support the full length sequence. The only difference between the four sequences was the length of 5’ poly-G and 3’ poly-A sequences, as such, the most parsimonious version was selected and deposited into GenBank. A bowtie2 alignment of the reads from each sample to this final viral genome sequence did not identify any SNPs for any sample, although some regions of heterozygosity are present among the samples. Nonetheless, the average read coverage per nucleotide using reads from each sample individually was high and ranged from 3548x to 15278x (S10C Fig). Similarly, average coverage per nucleotide was robust for the other fusariviruses identified, ranging from 180x for ShFV1 to 33888x for GtFV1 (S10C Fig).

### MiRV1

In addition to the three endornaviruses and one fusarivirus described above, a fifth viral sequence was identified in the *M. importuna* sample (PRJNA372858; unpublished). The single-segment sequence was determined to be 10,132 nt in length, with a single predicted ORF of 3,295 aa (Table 1). A conserved domain search revealed a viral methyltransferase domain and a viral helicase domain in addition to the RdRP_2 motif similar to positive-strand RNA viruses (S11C Fig). A BLAST-P search with the predicted protein sequence returned a single full length mycovirus sequence, *Sclerotinia sclerotiorum RNA virus L* (SsRV-L) [104], in addition to matches to Hepatitis Virus E and Cordoba virus. Other top hits include partial sequences corresponding to the RdRP domain of *Rhizoctonia solani RNA virus* 1, 2, and 3 (RsRV-1,RsRV-2, RsRV-3) [105]. Both MiRV1 and SsRV-L contain a single ORF with methyltransferase, helicase and RdRP domains, and neither contain peptidase motifs [104]. Previous phylogenetic analyses of SsRV-L and the three viruses from *R. solani* demonstrated a relationship between these viruses and members of the alpha-like virus superfamily, specifically viruses from the *Hepeviridae* and *Alphatetraviridae* families [104, 105]. *Endornaviridae*, whose members infect both fungi and plants, are also a family of alpha-like viruses [106], but appear to be more distantly related to this emerging group of mycoviruses. Expanding the phylogenetic analysis to include MiRV1 confirms that MiRV1 is a member of this new clade of mycoviruses (S11A Fig). Further, while a multiple sequence alignment of the RdRP motif sequences demonstrated the presence of the eight conserved RdRP motifs common among (+)ssRNA viruses, certain residues within the motifs were shared only among the mycoviruses that group with MiRV1 (S11B Fig). The low percent identity between MiRV1 and the related mycoviruses indicates that this is a new species of virus (S11D Fig). Identification of additional related viral sequences will help elucidate the relationship of MiRV1 and the rest of the Alpha-like viruses.

**S11 Fig.**
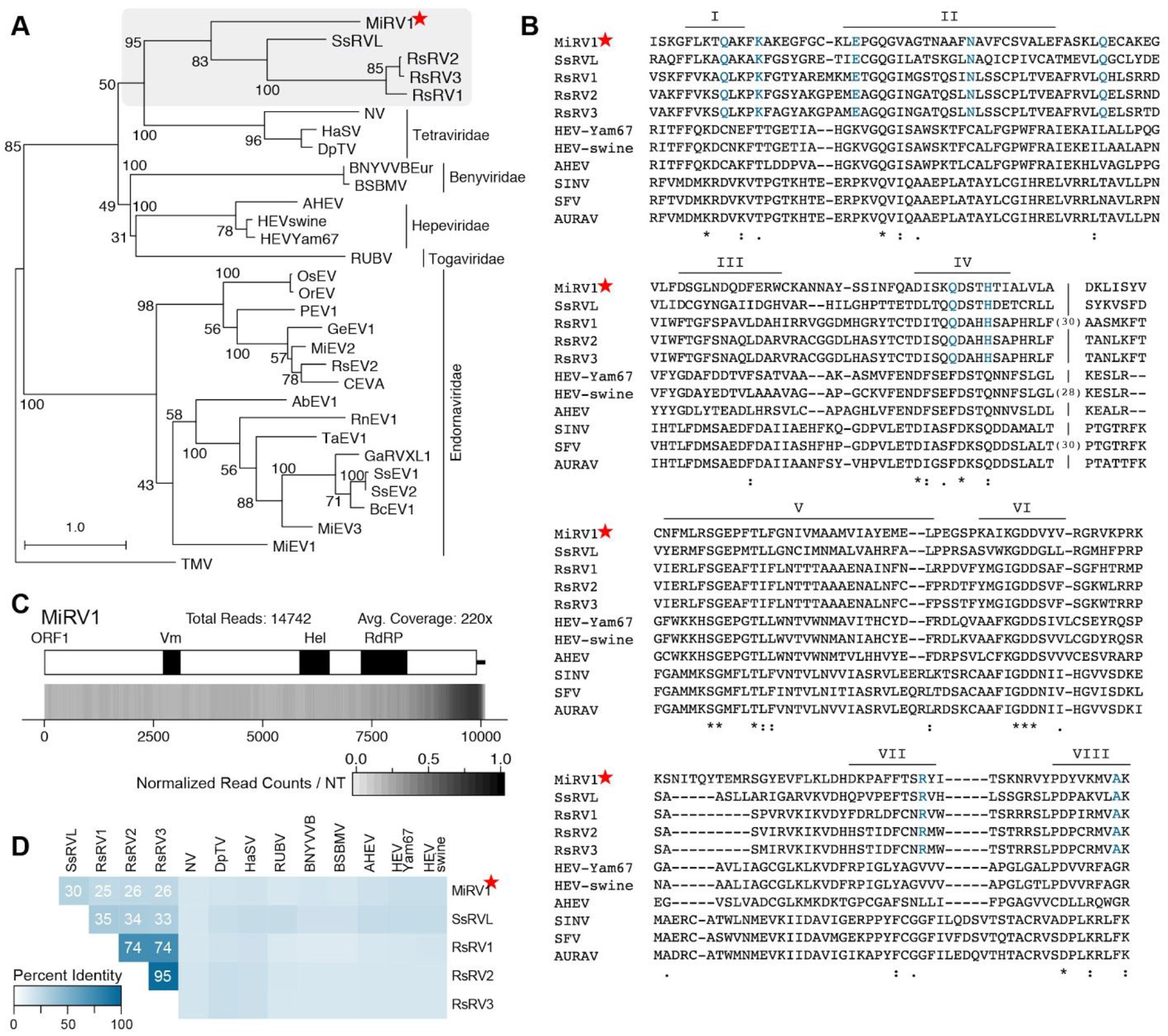
Sequence-based analyses of *Morchella importune RNA virus 1* (MiRV1). (A) Phylogram of the RdRP-motif for MiRV1, related mycoviruses, and Alpha-like viruses. Full names and accession numbers for protein sequences used are in Table S2, including the outgroup Tobacco Mosaic Virus (TMV). Scale bar represent 1.0 amino acid substitutions per site and numbers at the nodes indicate bootstrap support over 70% (1000 replicates). (B) Alignment of the RdRP motif from MiRV1 and related sequences. Conserved motifs within (+)ssRNA RdRP sequences are noted (I to VIII) and amino acid residues found only in the five related mycoviral sequences are noted in blue. Numbers in brackets are amino acids excluded from the alignment. (C) Density plot of read counts per nucleotide across the MiRV1 genome. Graphical depiction of the genome structure, predicted ORF, and encoded motifs are along the top, and a heatmap of normalized read counts is on the bottom. Read counts are normalized to a 0 −1 scale, where the maximum read count equals 1. Total reads mapped to the virus genome and the average read coverage are noted. (D) Percent identity matrix generated from Clustal 2.0 using the RdRP motif sequences from MiRV1, mycoviral relatives and select Alpha-like viruses. For sake of clarity, the 100% identity along the diagonal has been removed and only identities ≥25% are indicated. Red star notes MiRV1, the virus identified in this study. RdRP: RNA-dependent RNA Polymerase; Vm: Viral methyltransferase; Hel: Helicase.

### Unknown comovirus-like sequences

An RdRP-containing contig with a top match to Turnip Ringspot Virus (TRV) was identified in sequencing samples from *Colletotrichum tofieldiae* (;PRJNA287627 [107]) and *Zymoseptoria brevis* (PRJNA277175; [108]). Further inquiry also identified the second genomic segment of TRV, which encodes the movement and coat proteins, from both the *C. tofieldiae* and Z *brevis* Trinity contigs. The coding sequences of RNA1 and RNA2 are 98.5% and 99.7% identical, respectively, suggesting a close relationship between the sequences, despite the unrelated hosts.

As Turnip Ringspot Virus (genus: *Comovirus*; family: *Secoviridae)* can infect *Brassicaceae* [109], and a fungal host has not yet been described for viruses from this family, we postulate that this sequence may be due to cross contamination. The BioProject containing the *C. tofieldiae* sample included both media-grown fungi and an infection series of *Arabidopsis thaliana.* Further analysis of the infection series determined that the viral sequence is present in 19 of 24 (79%) of the mock-infected *A. thaliana* samples, 19 of 24 (79%) of the *A. thaliana* plus *C. tofieldiae* samples, and two of three fungus-only samples. An explanation for the uneven distribution of the viral sequences among these samples remains to be determined. Further, the *Z. brevis* sample was part of a media-grown, fungus-only transcriptomic experiment and as such, a putative source of the TRV sequences is not readily apparent. The *Z. brevis* sample is from Christian-Albrechts University of Kiel (Kiel, Germany) and was sequenced at the Max Planck Genome Center (Cologne, Germany) while the *C. tofieldiae* sample is from the Max Planck Institute for Plant Breeding Research (Kohn, Germany) and was also sequenced at the Max Planck Genome Center Cologne (Cologne, Germany). At this junction, it is not possible to determine the host or source of the viral genome sequences identified in the *C. tofieldiae* and *Z. brevis* samples.

## Summary and Conclusions

Here we describe the power of a systematic approach for analyzing public RNA-seq datasets to identify RNA virus genomes. Application of this bioinformatics pipeline to 569 transcriptomic datasets from 312 species of Pezizomycotina fungi identified 59 full-length viral genomes, 53 of which were determined to be new viruses.

The success of the pipeline was due to (a) the ability to fully assemble a viral RdRP coding sequence from as few as 136 reads (or 12x coverage) and (b) the strong signature of the viral RdRP domain. Benefits of this analysis include suitability to either rRNA-depleted total RNA or poly-A purified RNA. As viral particles enrichment is not required, which would select against non-particle forming viruses, virus identification via RNA-seq data can easily be determined from regular fungal transcriptomic experiments. Notable limitations of this approach include bias for viruses with a recognizable RdRP domain, which automatically excludes DNA viruses and may select against (-)ssRNA viruses. Further, robust analyses are required following the identification of putative viral sequences to confirm homology and relatedness to the suite of known viruses.

This analysis is not just an academic exercise in computational pipeline development and execution. It also works to answer the biological question, what fungi are infected by what viruses? It is becoming increasingly clear that there is a dynamic range of responses due to viral infection, outside of the well documented asymptomatic or hypovirulence phenotypes, that reflect the complex relationships present during multipartite interactions. A recent review by Roossinck [110] highlights a number of mutualistic or symbiogenic relationships between viruses and their hosts. This includes killer yeasts: yeast that are infected with a virus which produces toxins that kill competitors to the host fungi while conferring resistance to the host [111]. Another example is the tripartite interaction between the tropical panic grass *Dichanthelium lanuginosum*, the endophytic fungi *Curvularia protuberata* and *Curvularia* thermal tolerance virus, the virus that infects *C. protuberata.* Thermal tolerance to high soil temperatures is only conferred to the plant when the fungi is present and is infected with the virus [112]. Another phenotype, different from canonical hypovirulence, is the reduction in fungicide resistance observed following viral infection of prochloraz-resistant strain of *Penicillium digitatum* [113].

Multipartite relationship also include virus-virus interactions within a host as many fungi are concurrently infected by more than one mycovirus; here we identified over 10 such fungi. Recently, it was described that a positive effect on accumulation of a viral genome was observed when a second, unrelated virus was present in *Rosellinia necatrix* [75]. Conversely, coinfection of *C. parasitica* by a totivirus and a suppressor-deficient hypovirus lead to induction of antiviral defense genes and ultimately inhibition of totivirus replication [114].

Indeed asymptomatic or latent infections can downplay the active antiviral processes at work within the fungus and the important role a successful antiviral defense has on fungal phenotypes. Small RNA-mediated antiviral defense systems, crucial for preventing a deleterious effect on fungal health, have been identified in *Cryphonectria parasitica* [115], *Colletotrichum higginsianum* [8], and *Sclerotinia sclerotiorum* [116]. Further, it can be assumed that antiviral defenses have contributed to the under-identification of mycoviral infections in fungi, including many of the fungi identified here that are infected with a virus, but lack studies into RNA silencing.

This systematic analysis also highlights the uneven distribution of sequencing data among members of the Pezizomycotina subphylum. While strides are being made, particularly due to large-scale approaches such as the 1000 Fungal Genomes Project, a fuller understanding of the mycoviral landscape and therefore the effect a virus has on fungal phenotype requires continuous inquiry. As the number of Pezizomycotina fungi is estimated to be greater than 32,000 [117] and the current scope of known fungal hosts is less than 500 species, it stands to reason there remains a wide range of viral diversity yet to be identified and characterized. Further, this study focused solely on the subphylum Pezizomycotina, meaning members of the sub-phylums Saccharomycotina and Taphrinomycotina and the unranked group Ascomycota incertae sedis are not included. Beyond the fungal kingdom, what about datasets from the animal and plant kingdoms? While the computational requirements are large to investigate these tens of thousands of RNA-seq datasets, the rate limiting steps are the manual curation required at the start of the pipeline, to analyze metadata and identify suitable samples, and at the end of the pipeline, to evaluate and characterize the identified viral genomes. Other bioinformatic approaches under consideration are (a) downsampling the reads to a suitable sized input for the pipeline and ? or (b) direct translation of the raw reads followed by analysis for RdRP domain signatures. Finally, sequencing technologies that produce reads >1 kb in length provide an even more direct route to virus detection as entire viral genomes could be sequenced as single reads. For all future RNA-seq experiments, we highly recommend that the data be analyzed for the presence of viral RdRP signatures as part of any routine bioinformatic pipeline.

## Materials and Methods

### Identifying publicly available fungal RNA-seq datasets

Programs available via the Entrez Programming Utilities (“E-utilities”) were used to search the NCBI Short Read Archive (SRA) for datasets that matched the search criteria. For ease of searching, the fungal kingdom was divided into three phylum level searches: ascomycota (subphylum Pezizomycotina), basidiomycota, and other. General search criteria used for all phylum-level searches included the phrase “NOT LS454 NOT PACBIO NOT ABLSOLID AND biomol rna [PROP]” to identify RNA-seq datasets from Illumina (or Solexa) instruments. The E-utilities commands esearch and efetch combined to create a comma-separated table of all matching SRA records.

### Pipeline for identification of viral-like sequences

The output file from the E-utilities search command was manually parsed to create a unique list of all BioProjects and fungal species. Multiple fungal species found under one BioProject number were split into separate entries. Most BioProjects included more than one sample; for example different environmental treatments or wild-type versus mutant. In general, the SRA sample selected for analysis was a wild-type-like strain grown on media. When available, the fungal genome sequence was either downloaded from NCBI or Joint Genome Institute (JGI).

The computational steps for identifying viral sequences start with a downloaded raw reads file from SRA. When a fungal genome sequence was available, RNA-seq reads were aligned to the genome using bowtie2 (version 2.2.9) [118], and the unaligned reads were captured in a separate file using the flag ‘--un’. A *de novo* assembly of these unaligned reads was created using Trinity (v2.1.1) [119] and resultant contigs were analyzed for open reading frames (ORFs) by TransDecoder (version 2.1) [120]. Finally, these predicted protein sequences were queried against a custom database of viral RNA dependent RNA polymerases (RdRPs) using HMMscan (version 3.1b2; [121]). In general, when the data was paired end, the data present within pair one was sufficient data for identification of viral contigs. Occasionally read depth from the virus genome was low, resulting in fragmented viral contigs. In these instances, the pipeline was rerun using both pair one and pair two data. Due to the nature of the ‘--un’ flag of bowtie2, the pairs were treated as individual single-end datasets, instead of paired-end data. Assembly of the virus from *Trichoderma harzianum* (PRJNA216008), was particularly recalcitrant and as such, all six SRA paired-end datasets were used in the pipeline for identification of ThAV1.

### Contig extension and identification

Contigs identified containing putative viral RdRP domains were further analyzed by first aligning the nucleotide sequence to the ‘nr’ database at NCBI using BLASTX. A variety of manual annotation steps were required following confirmation of a viral sequence. Some viral genomes, such as those matching the *Partitiviridae* or *Chrysoviridae* families, were predicted to contain additional viral genes encoded on separate contigs; these ORFs encode for coat proteins and/or hypothetical proteins. S2 Table summarizes the viral RdRP sequences identified and, where applicable, which known BLASTX-identified sequences were used to query the Trinity assembly for additional contigs.

Occasionally contigs were identified where the predicted ORF(s) extended to either the 5’- or 3’-most edge; this indicated that the sequence in hand was truncated. In these instances, the RdRP sequence from the best BLAST match was used to query the Trinity contigs to identify putative viral sequences outside the known RdRP contig. Any contigs identified were then stitched together to create a final sequence with a full-length ORF. Start and stop codons were predicted using ORFinder at NCBI [122].

## Bioinformatic Tools and Analyses

### Sankey Diagram

The online tool SankeyMATIC [123] was used generate the Sankey Diagram in Fig 1. Data tables used as input are available through the FigShare FileSet (DOI: 10.6084/m9.figshare.7476359).

### Pseudoknot Structures

The online tool DotKnot was used to predict pseudoknots in certain viral genome sequences [124, 125]. PseudoViewer 3.0 was used for visualization of the predicted pseudoknots [126].

### Read distribution plots

Distribution of reads along the viral genome segment(s) was visualized by first aligning the reads to the viral genome using Bowtie2, then calculating the hits per nucleotide along the viral sequence(s) using the custom Perl script plotRegion_RNAseq_hitsPerNT.pl. The resulting output table was the input for an R script to create hits per nucleotide heat maps for each viral segment. The R code for each plot, along with the input data table, is available through the FigShare FileSet (DOI: 10.6084/m9.figshare.7476359).

### Phylogenetic analysis

To identify related sequences for phylogenetic analysis, the predicted amino acid sequence for the RdRP was used to query the ‘nr’ database at NCBI. Protein sequences of the top hits, generally with e-values approaching or equal to 0.0, were downloaded. Partial sequences were excluded. Known viral sequences that appeared as the top hit for multiple viruses identified in this study were deduplicated. The complete list of Accession IDs used for phylogenetic analysis is available in Supp. Table S2.

To build the trees, amino acid sequences were aligned with MAFFT [127], and resultant alignments were degapped and trimmed using the tool trimAL [128] with a gap threshold of 0.9 (allowing for a gap in up to 10% of the sequences at a given position). Maximum likelihood phylogenetic trees were constructed with RAxML version 8.2.4 [129] using the LG model for amino acid substitution. RAxML tree files, with branch lengths, were viewed in Dendroscope version 3.5.7 [130] and exported as PDF for figure preparation.

### Percent Identity Matrices

Percent identities among protein coding sequences, either full length or motif-specific, were determined via multiple sequence alignment using Clustal Omega [131]. In instances where two different proteins were evaluated and then combined into one matrix, first the RdRP protein sequences were analyzed; the order of the output in the percent identity matrix was then applied to the second set of protein sequences to be analyzed and the option “Order: Input” was selected in Clustal-Omega. Percent identity matrices were converted to heat map plots using a custom R script. This script along with the input matrices are available through the FigShare FileSet (DOI: 10.6084/m9.figshare.7476359).

### Accession numbers

Nucleotide sequences of the mycoviral genomes identified and analyzed in this study were deposited at NCBI; GenBank accession numbers are listed in Table 1.

## Acknowledgments

We are deeply grateful to researchers around the world that have shared their transcriptomic data, both before and after publication. We particularly wish to acknowledge the sequencing data produced by various groups at the US Department of Energy Joint Genome Institute (http://www.jgi.doe.gov). We also thank the Bioinformatics Facility staff at the Danforth Center for excellent computational support.

## Author Contributions

KBG, EEH, and JCC designed the research. KBG, EEH, and RLA performed the research and analyzed the data. KBG, EEH, and JCC drafted the paper; all authors edited and approved the manuscript.

---

**Table S1:**
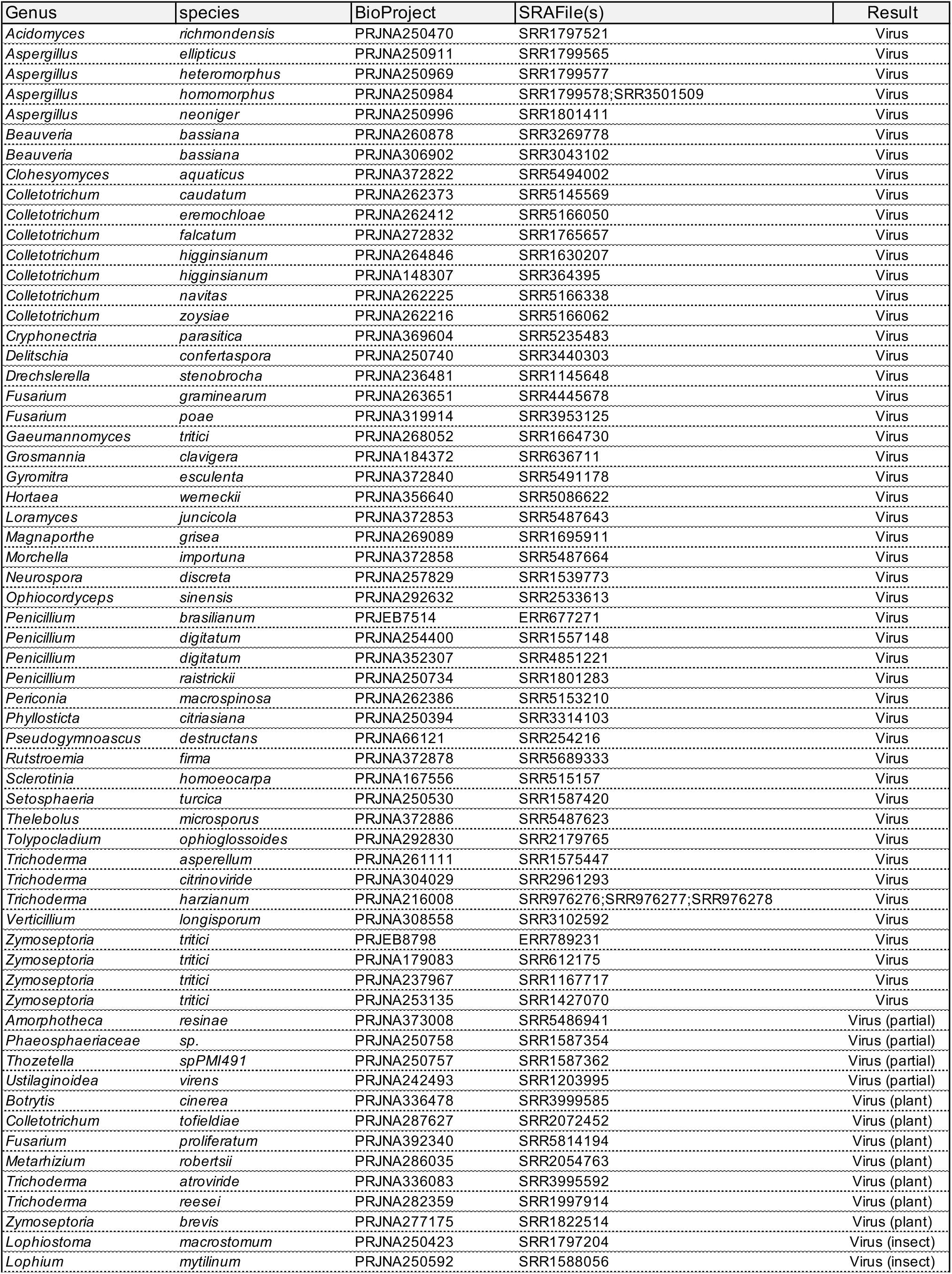
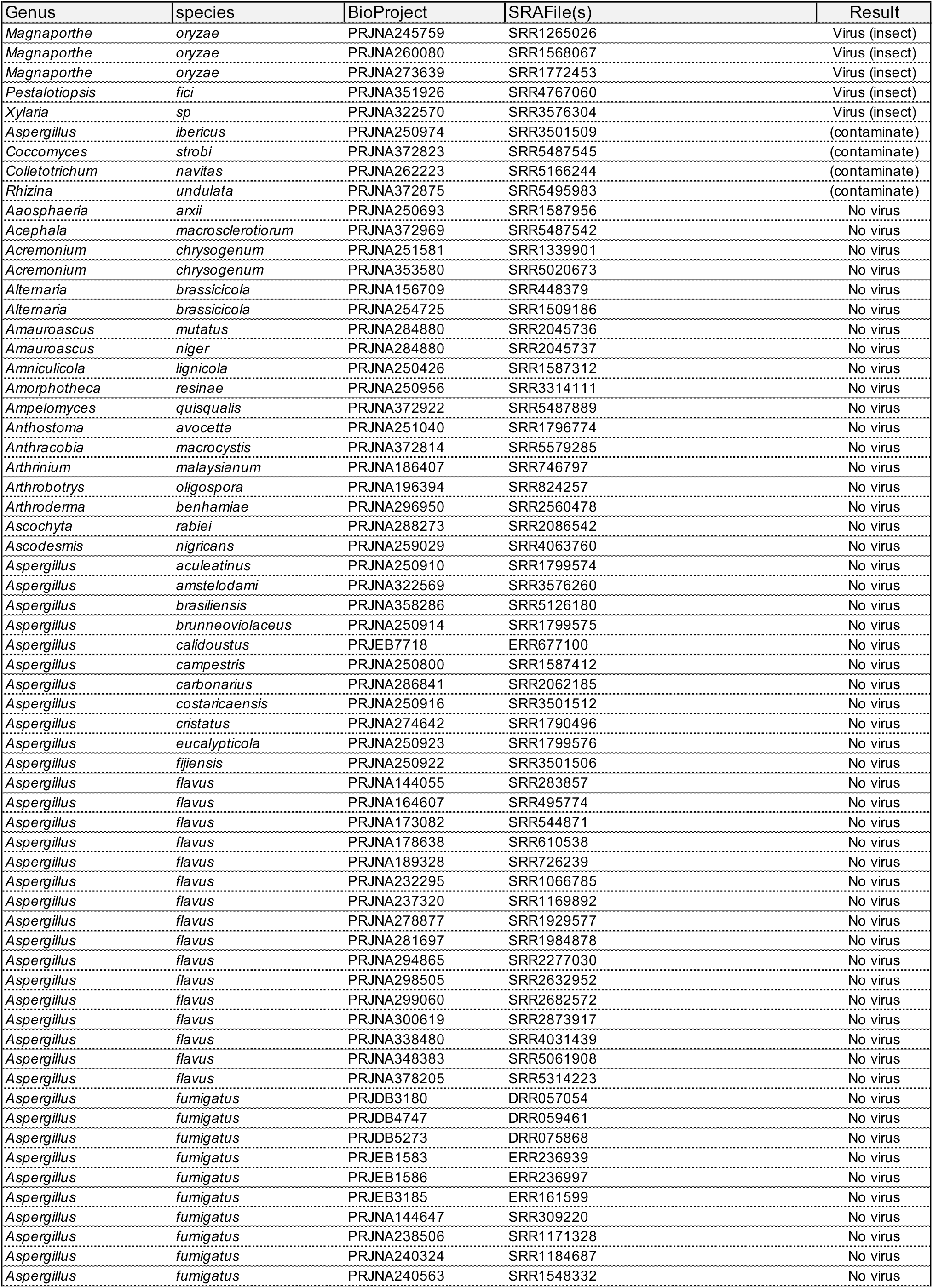
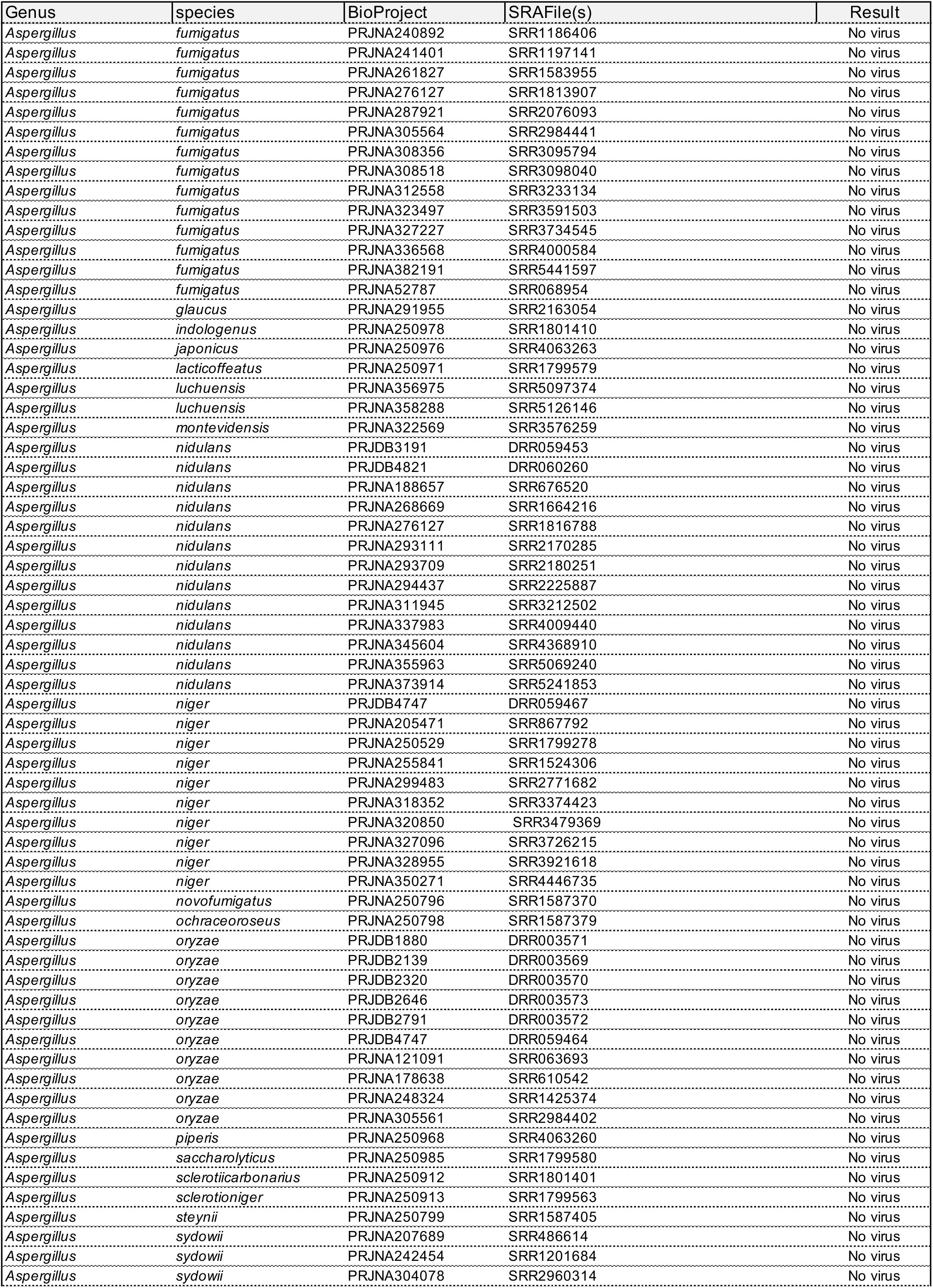
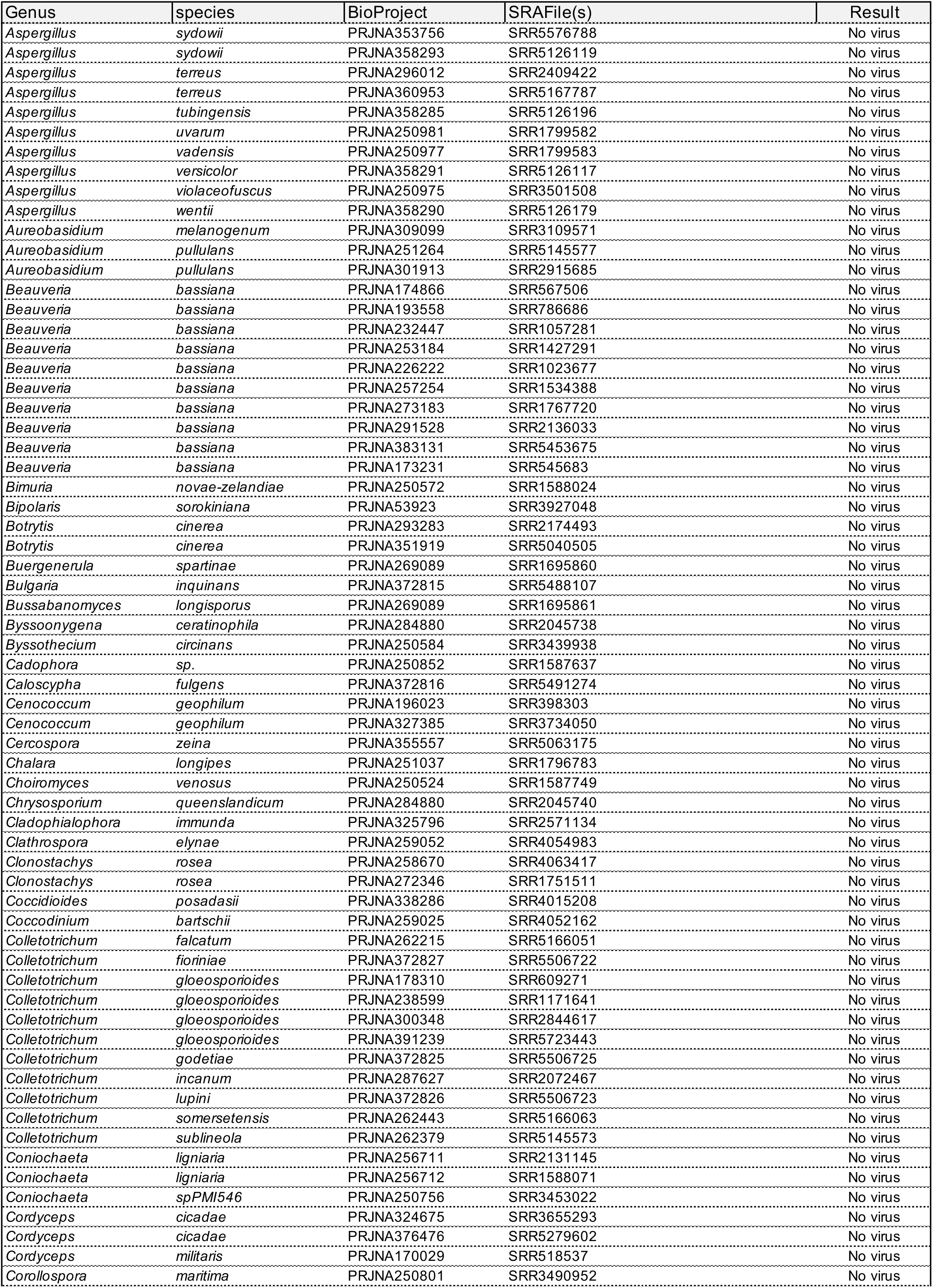
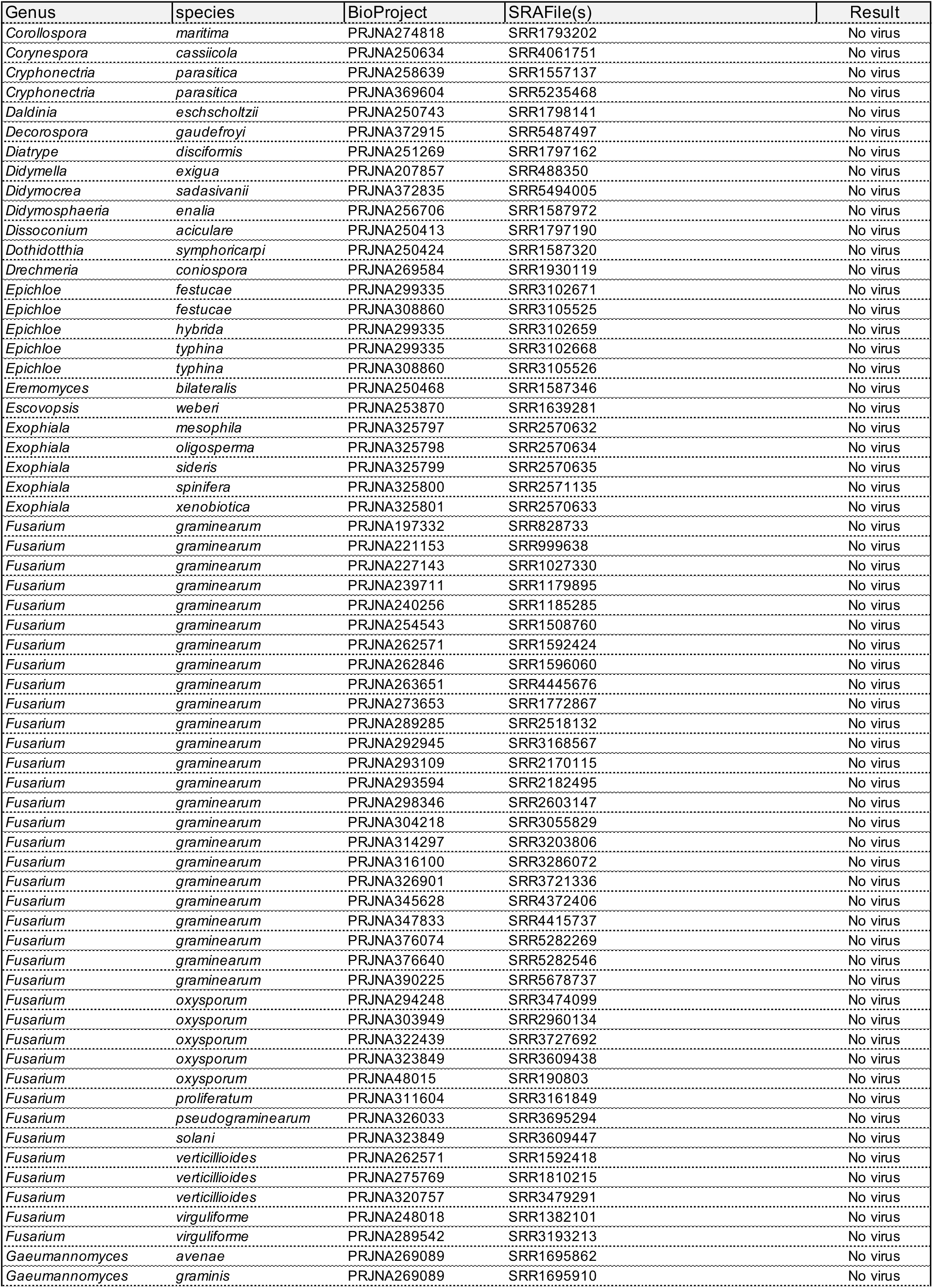
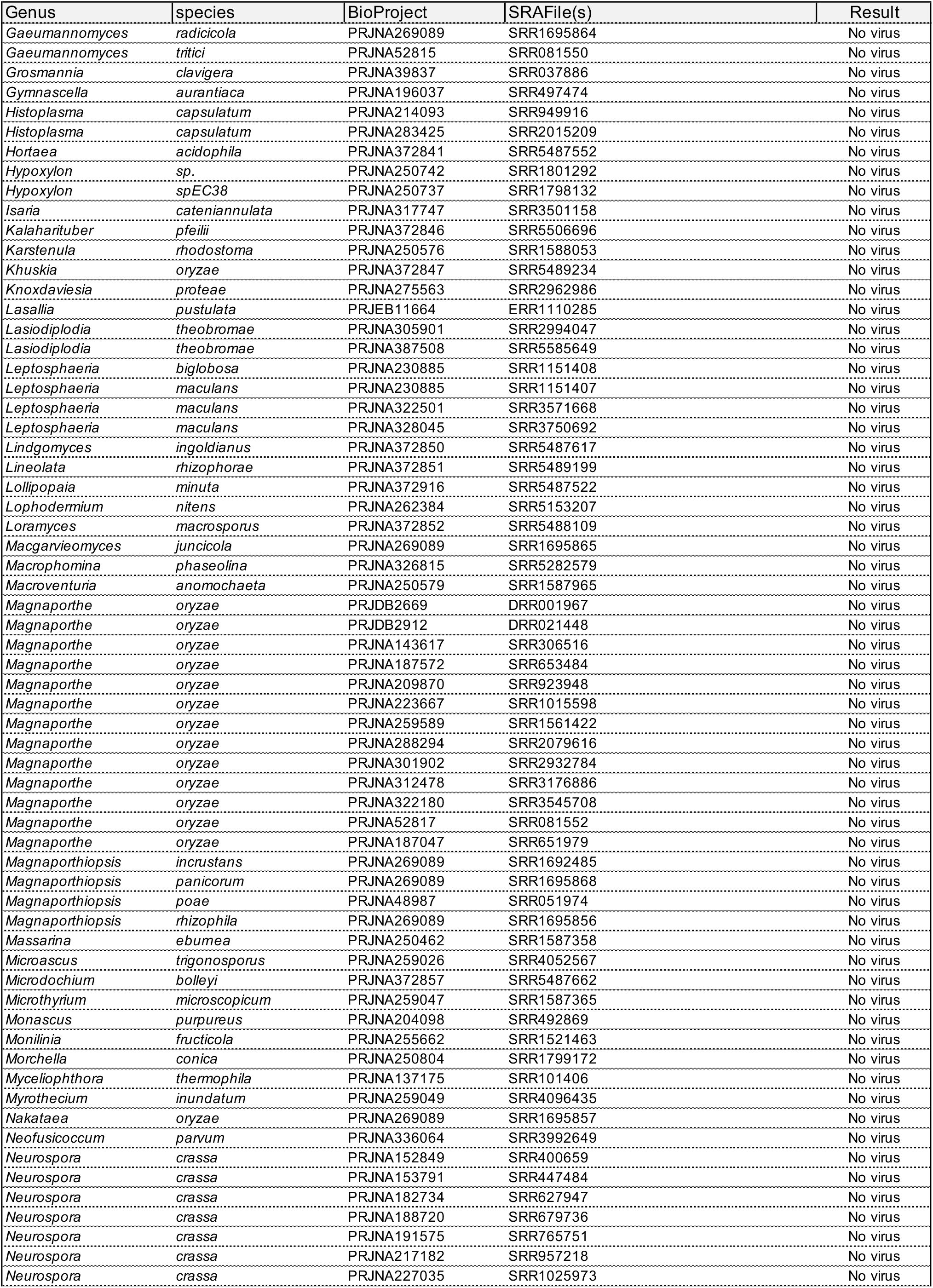
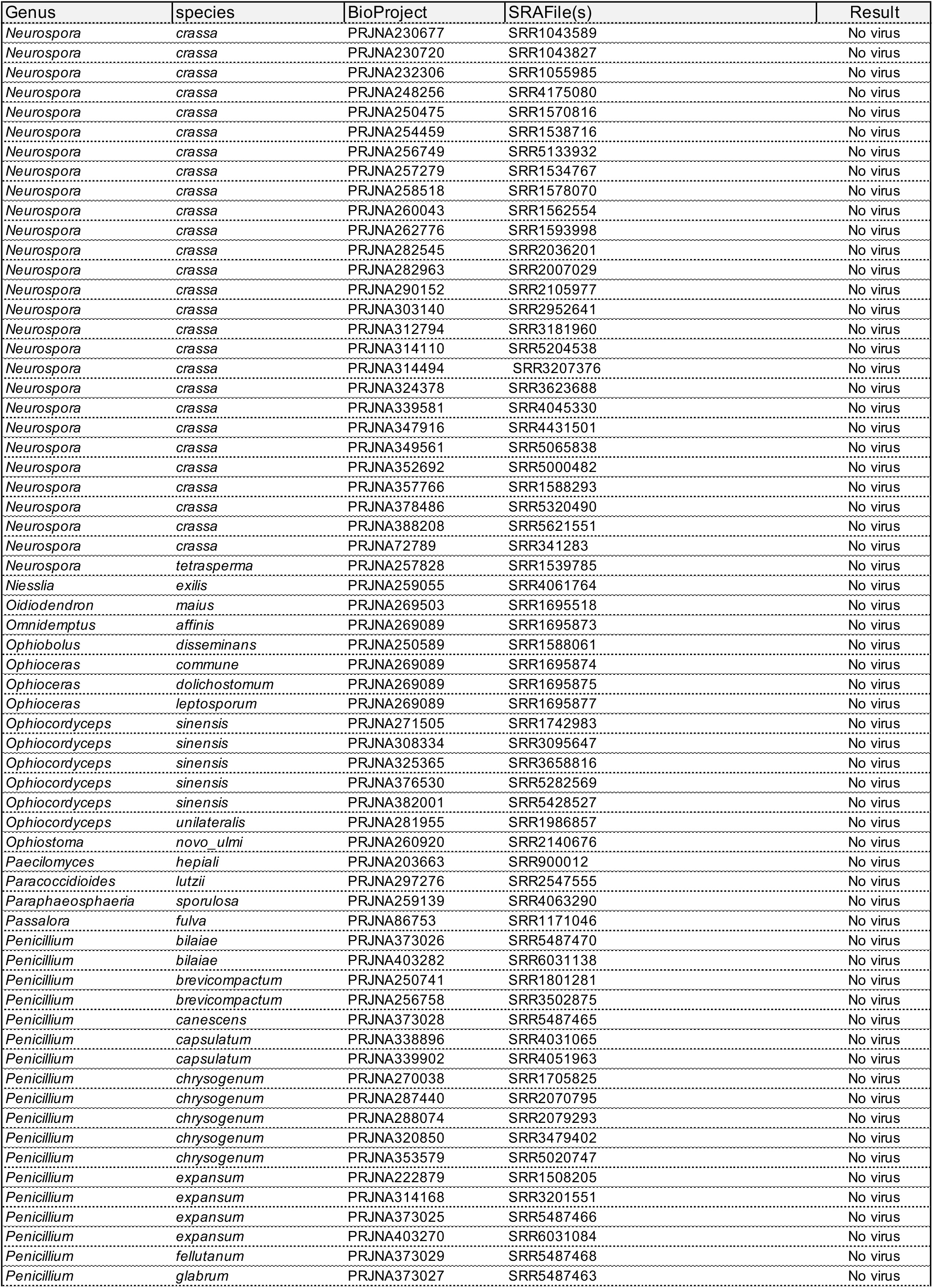
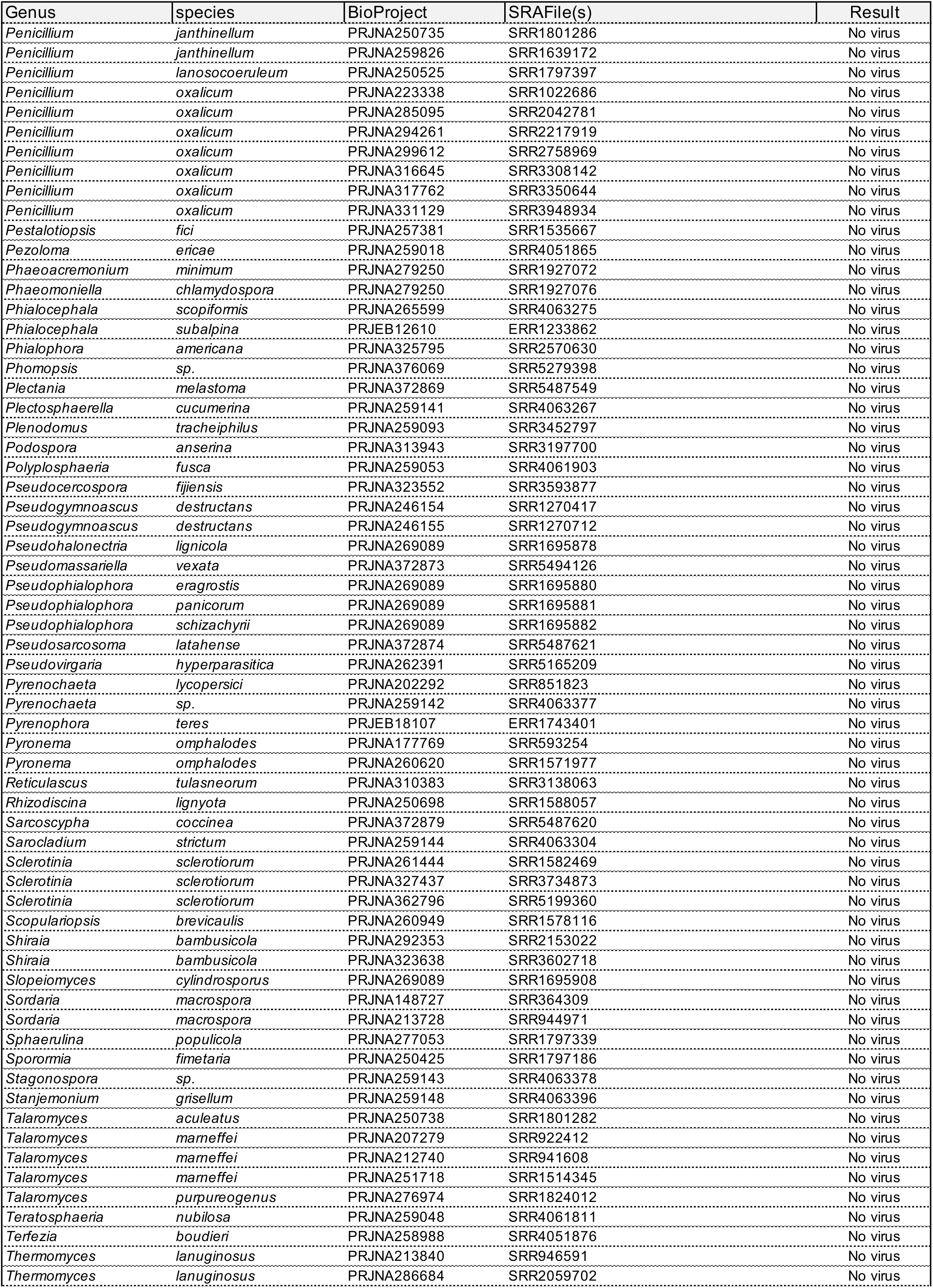
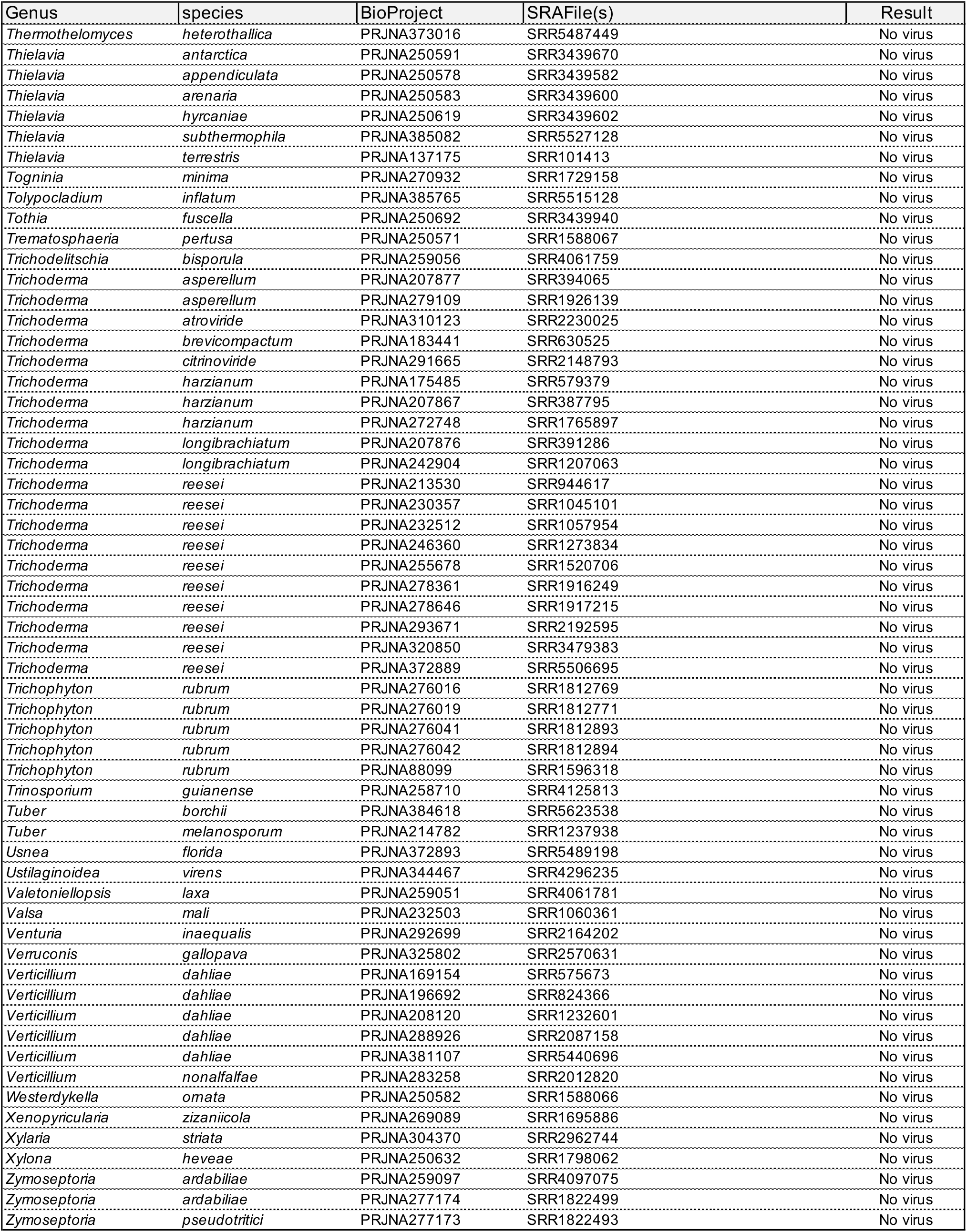
Complete list of BioProjects and SRA datasets analyzed

**Table S2:**
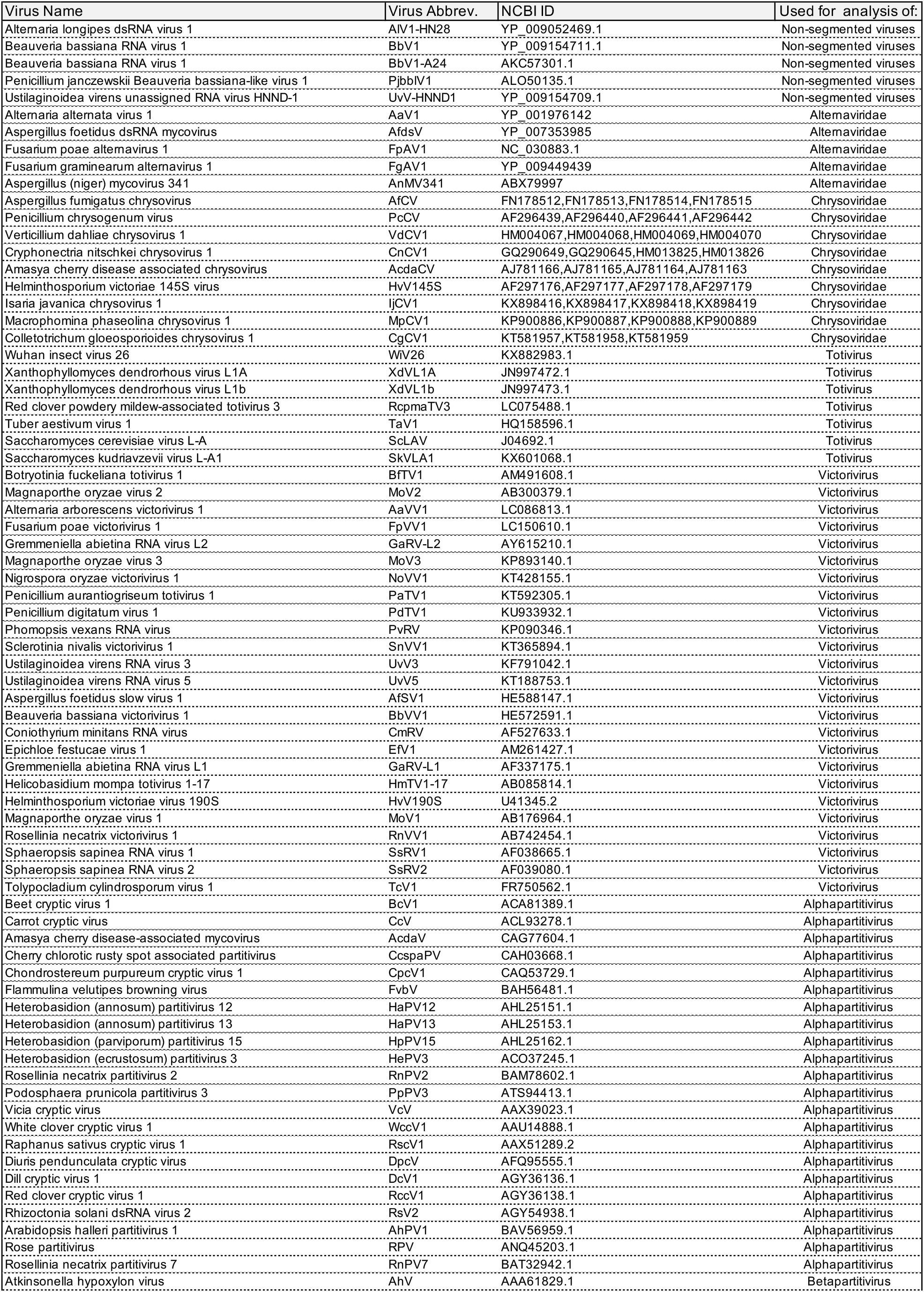
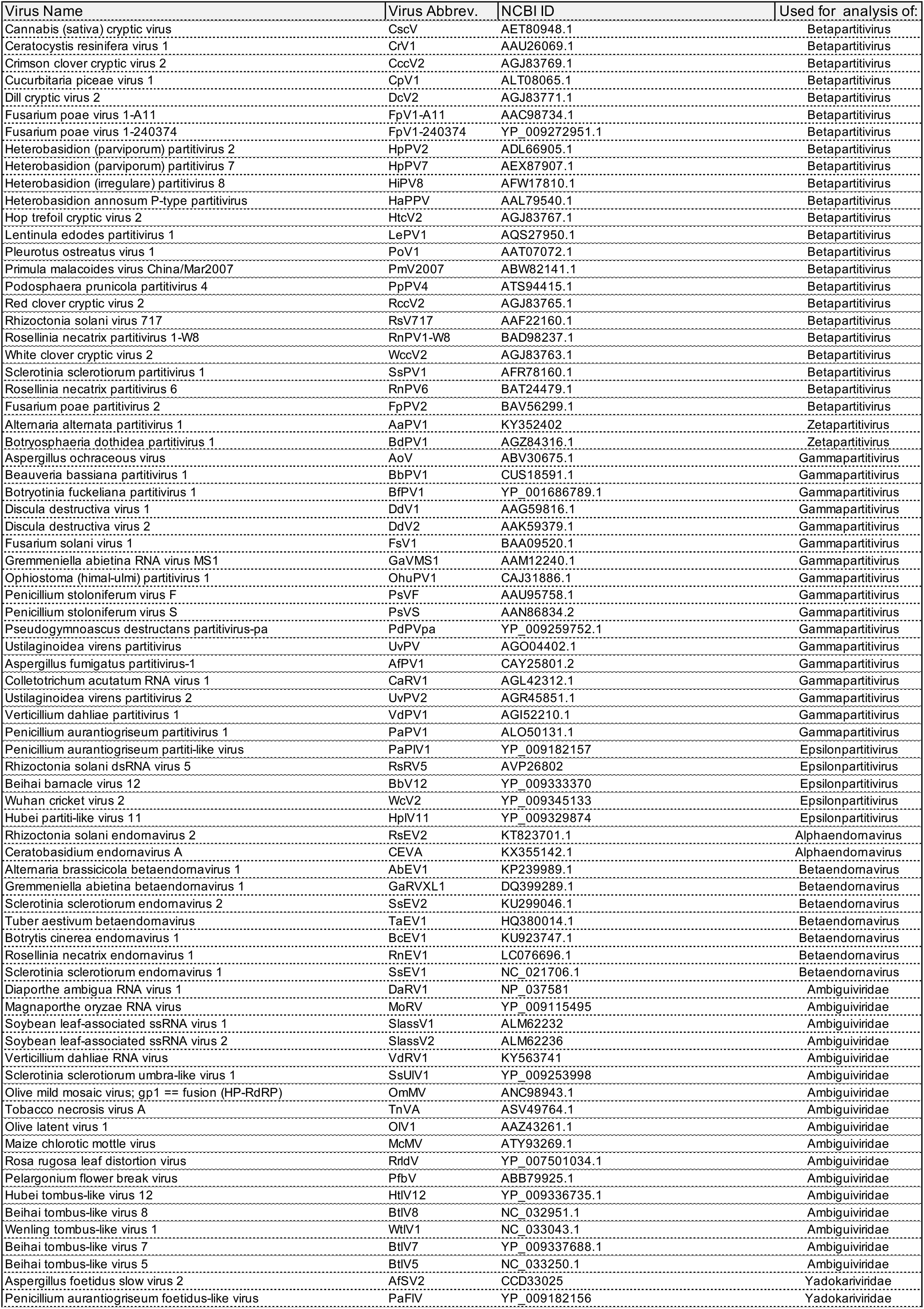
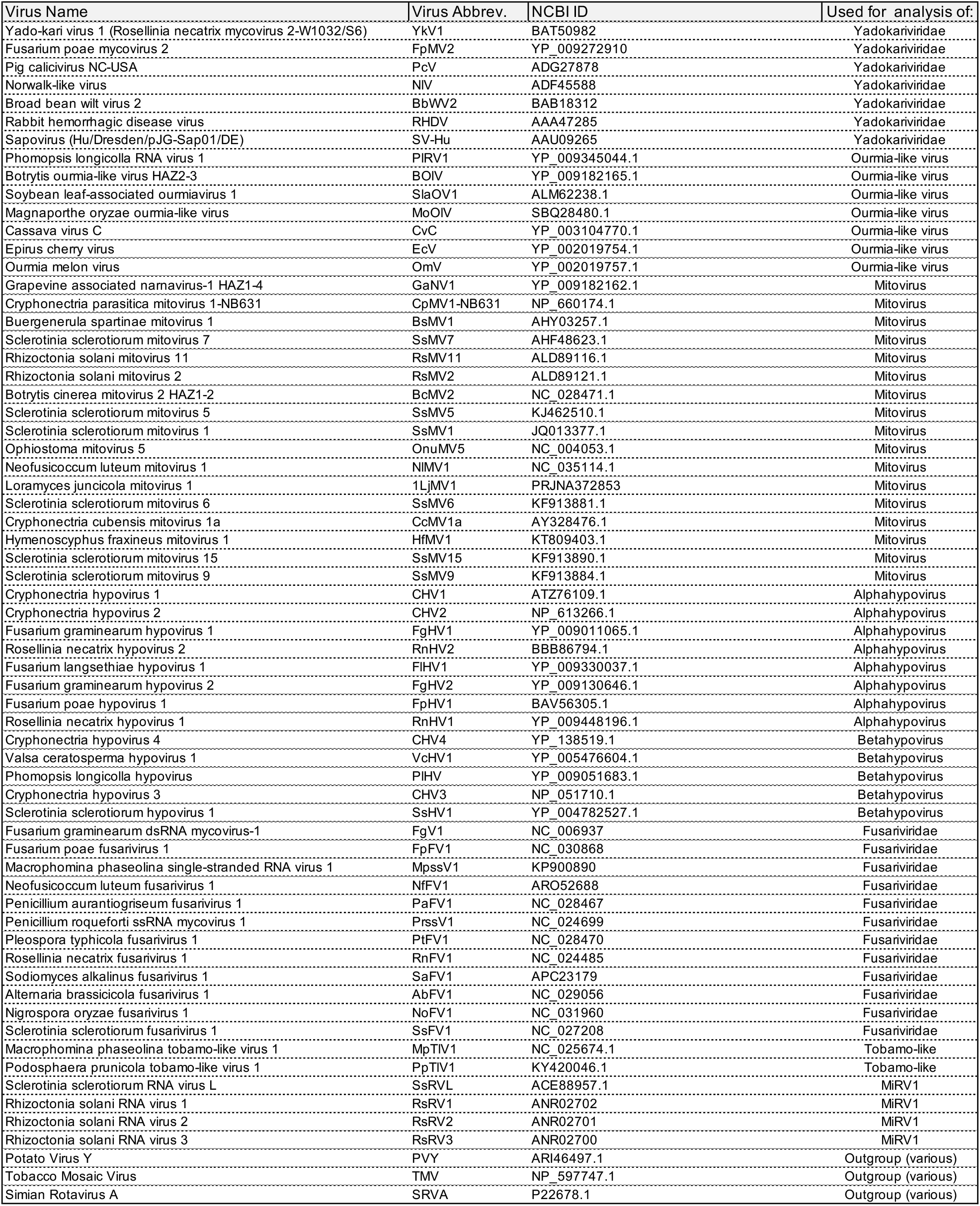
Name and Accession IDs of mycoviruses related to those identified in this study

